# CanFlyet: Habitat Zone and Diet Trait Dataset for Diptera Species of Canada and Greenland

**DOI:** 10.1101/2023.03.15.532812

**Authors:** S.E. Majoros, S.J. Adamowicz

## Abstract

**Background:** True flies (Diptera) are an ecologically important group that play a role in agriculture, public health and ecosystem functioning. As researchers continue to investigate this order, it is beneficial to link the growing occurrence data to biological traits. However, large-scale ecological trait data are not readily available for fly species. While some databases and datasets include fly data, many ecologically relevant traits for taxa of interest are not included. In this study we create a dataset containing ecological traits (habitat and diet) for fly species of Canada and Greenland having occurrence records on the Barcode of Life Data Systems (BOLD). We present a dataset containing trait information for 981 Diptera species.

**New Information:** Diptera were chosen for the dataset based on the occurrence records available for Diptera species from Canada and Greenland on the Barcode of Life Data Systems (BOLD). Trait data were then compiled based on literature searches conducted from April 2021 - April 2024 and assigned at the lowest taxonomic level possible. Three biological traits were included: habitat, larval diet, and adult diet. The dataset contains traits for 981 species across 380 genera, 34 subfamilies, and 61 families. This dataset allows for assignment of traits to occurrence data for Diptera species and can be used for further research into the ecology, evolution, and conservation of this order.

## Introduction

True flies (Diptera) are a diverse, widespread taxonomic group, occurring across a wide range of ecological niches and geographic regions (Adler & Courtney, 2019, Marshall, 2012). This important group occupies various ecological roles, many of which have an impact on the environment and agricultural practices. Fly species can act as pests to crops, such as *Liriomyza bryoniae* on cabbages, lettuce, and tomatoes (Bragard et al., 2020); *Delia radicum* on broccoli and cauliflower (Mesmin et al., 2019); *Dasineura mali* on apples (Wearing et al., 2013); and *Bradysia ocellaris* and *Lycoriella ingenua* on mushrooms (Shamshad 2010). Diptera also includes pests of livestock such as the blood-feeding *Stomoxys calcitrans* (Olafson et al., 2021). Flies are also a group of interest with regard to public health, as some, like *Calliphora vomitoria*, are vectors of pathogenic microorganisms (Bedini et al., 2017).Despite these detrimental examples, flies also provide many important ecological benefits. Flies are an important pollinator group, especially in Arctic and Northern environments (Tiusanen et al., 2016). The family Muscidae includes species that are pollinators of key plant species in this region and play an important role in the ecosystem (Tiusanen et al., 2016). Flies also fill various other ecological roles, such as parasitoids of other insects that act as vectors for pathogenic bacteria (Molinatto et al., 2020). Many species act as ecosystem engineers; changing environments through suspension-feeding, grazing, burrowing, and predation, as well as serving as an important food source for other organisms (Adler & Courtney, 2019). The saprophagous diet of many flies is also important for the breakdown of decaying organic material (Adler & Courtney, 2019). As biocontrol agents and bioindicators of water quality, flies act as a valuable resource for food production and research (Adler & Courtney, 2019).

This ecologically important group has been the focus of large DNA barcoding efforts, such as the Global Malaise Trap Program, which seeks to document Diptera diversity (which is likely under described) and open doors for ecological research and monitoring (deWaard et al., 2018, Geiger et al., 2016, Global Malaise Program, n.d.). DNA barcoding involves using a standardized gene region to identify and delineate species (Hebert et al., 2003), and there is a large amount of DNA barcoding data available on databases such as the Barcode of Life Data System (BOLD; Ratnasingham and Hebert, 2007), which contained 8 591 083 Diptera records representing 42 728 species as of May 28^th^, 2024. There is also a large amount of species occurrence data on databases such as the Global Biodiversity Information Facility (GBIF; GBIF.org, 2023), which contained 17 796 378 occurrences as of May 28^th^, 2024. As Diptera research continues, it is valuable to link such occurrence records with biological data.

However, finding biological trait data for fly species can be a challenge. There are various databases or datasets currently available for Diptera and other insect groups, covering a varying number of traits and taxa (Table 1). There are few databases that focus solely on Diptera, and while flies are included in some databases, not all species and traits of interest are included. A larger focus also appears to be on morphological traits, such as body size, with information on ecological traits, such as habitat and diet, being less readily available. Finally, few databases focus on Diptera from Canada or Greenland. Overall, traits for many Diptera species are very difficult to find and, when known, are typically present in taxon-specific studies rather than compiled datasets. The goal of this study was to create a dataset containing ecological traits (habitat and diet) for selected Diptera species found in Canada and Greenland. Species were selected that have multiple occurrence records and high-quality DNA barcode sequences on BOLD (following criteria outlined in Majoros et al. (2023)), which likely includes many of the most common species in the study region. This dataset can be used in future research to further our knowledge and understanding of this important taxonomic group.

**Table 1:** Overview of some of the databases and datasets that include insects currently publicly available. Number of records are as of Feb 2^nd^, 2023.

| Database Name | Taxonomic Focus | Geographic Focus | Types of Traits | Number of Traits | Number of Records | Data Availability | References |
| --- | --- | --- | --- | --- | --- | --- | --- |
| DISPERSE<sup>1</sup> | Annelida, Mollusca, Platyhelminthes & Arthropoda | Europe | Dispersal | 9 | 480 | Download as xlsx file | Sarremejane et al. (2020) |
| Freshwater Biological Traits<sup>2</sup> | Freshwater macroinvertebrate taxa | North America | Morphology, life history, mobility, environmental tolerance, resource acquisition/preference | 26 | 11912 | Select taxa, region and traits on website. Download as an xlsx file | United States Environmental Protection Agency (2012) |
| Animal Traits<sup>3</sup> | Arthropoda, Chordata, Mollusca and Annelida | Global | Body mass, metabolic rate, brain size | 3 | 3580 | Download xlsx or csv file | Herberstein et al. (2022) |
| CONUS<sup>4</sup> | Arthropoda | United States of America | Life history, dispersal, morphology, ecology | 11 | 2.05 million | Download csv file | Twardochleb et al. (2021) |
| Lotic Invertebrate Traits for | Invertebrates | North America | Ecology, morphology, behaviour, physiology | 62 | 14127 | Download as a txt file | Vieira et al. (2006) |

<sup>1</sup>https://figshare.com/articles/dataset/DISPERSE_a_trait_database_to_assess_the_dispersal_potential_of_European_aquatic_macroinvertebrates/12417251/1
<sup>2</sup> https://cfpub.epa.gov/ncea/global/traits/search.cfm
<sup>3</sup> https://animaltraits.org/
<sup>4</sup> https://portal.edirepository.org/nis/mapbrowse?packageid=edi.481.5

| Database Name | Taxonomic Focus | Geographic Focus | Types of Traits | Number of Traits | Number of Records | Data Availability | References |
| --- | --- | --- | --- | --- | --- | --- | --- |
| North America<sup>5</sup> |  |  |  |  |  |  |  |
| European & Maghreb Butterfly Trait Database<sup>6</sup> | Lepidoptera | Europe and North Africa | Life history, morphology, resource-based, behaviour | 25 | 542 | Download as xlsx file | Middleton-Welling et al. (2020) |
| GlobalAnts<sup>7</sup> | Formicidae | Global | Morphology, ecology, life history | 23 | 82910 | Need to make an account to access. Can search for desired traits and download as a csv file | Parr et al. (2016) |
| The Odonate Phenotypic Database<sup>8</sup> | Odonata | Global | Morphology, life history, behaviour | 35 | 3978 | Download as a xlsx file | Waller et al. (2019) |
| Morphological trait database of saproxylic beetles<sup>9</sup> | Saproxylic Coleoptera | Europe | Morphology | 13 | 1376 | Download as a csv file | Hagge et al. (2021) |
| Data from: Sensitivity of functional diversity metrics to | Carabidae | The Netherlands | Morphology | 7 | 73 | Download as txt file | van der Plas et al. (2017) |

<sup>5</sup> https://pubs.er.usgs.gov/publication/ds187
<sup>6</sup> https://butterflytraits.github.io/European-Butterfly-Traits/index.html
<sup>7</sup> http://globalants.org/
<sup>8</sup> http://www.odonatephenotypicdatabase.org/shiny/odonates/?_inputs_&choose_species=%22%22
<sup>9</sup> https://datadryad.org/stash/dataset/doi:10.5061/dryad.2fqz612p3

| Database Name | Taxonomic Focus | Geographic Focus | Types of Traits | Number of Traits | Number of Records | Data Availability | References |
| --- | --- | --- | --- | --- | --- | --- | --- |
| sampling intensity<sup>10</sup> |  |  |  |  |  |  |  |
| Data from: A summary of eight traits of Coleoptera, Hemiptera, Orthoptera and Araneae, occurring in Grasslands in Germany<sup>11</sup> | Arthropoda | Germany | Morphology, ecology | 8 | 1230 | Download as txt file | Gossner et al. (2015) |
| LepTraits<sup>12</sup> | Lepidoptera | Global | Morphology, habitat, reproduction, hostplant association | 6 | 75103 | Download as csv file | Shirey et al. (2022) |
| The Insect Trait Tool (ITT)<sup>13</sup> | Arthropoda | Germany | Habitat, Diet | 2 | 34085 | Download as a pdf or .xlsx file | Hörren et al. (2022) |

<sup>10</sup> https://datadryad.org/stash/dataset/doi:10.5061/dryad.1fn46
<sup>11</sup> https://datadryad.org/stash/dataset/doi:10.5061/dryad.53ds2
<sup>12</sup> https://springernature.figshare.com/collections/LepTraits_1_0_A_globally_comprehensive_dataset_of_butterfly_traits/5899187/1
<sup>13</sup> https://www.biorxiv.org/content/10.1101/2022.01.25.477751v1.full

### General Description

This study provides a dataset containing the habitat and adult and larval diet for Diptera species of Canada and Greenland having occurrence records on BOLD. The traits were assigned to taxa using the currently available literature. This dataset will aid in further research that involves the ecological traits of Diptera species.

### Project Description

#### Title

CanFlyet: Habitat Zone and Diet Trait Dataset for Diptera Species of Canada and Greenland

#### Personnel

Samantha E. Majoros & Sarah J. Adamowicz

#### Design Description

The dataset provides habitat, larval diet, and adult diet for Diptera species found in Canada and Greenland. Traits were determined based on a literature search.

#### Funding

Funding support for this project comes from the Natural Sciences and Engineering Research Council of Canada; the Government of Canada through Genome Canada and Ontario Genomics; the Ontario Ministry of Economic Development, Job Creation and Trade; and Food from Thought: Agricultural Systems for a Healthy Planet Initiative program funded by the Government of Canada through the Canada First Research Excellence Fund.

### Sampling Methods

Description: The fly species were chosen for inclusion in this dataset by first downloading data for Diptera from Canada and Greenland from BOLD directly into R using BOLD’s application programming interface (API) on June 24^th^, 2021. The records were filtered based on the requirements outlined in Majoros et al. (2023), and the remaining species were chosen for analysis and inclusion in this dataset. In Majoros et al. (2023), species were represented by Barcode Index Numbers (BINS), which are operational taxonomic units (OTUs) used by BOLD that are clusters of barcode sequences similar to species (Ratnasingham & Hebert, 2013). For other portions of the analysis, records were clustered by a 4% clustering threshold, and this was used in the place of BINs. Records needed to possess high quality DNA sequence data to be included in the analysis. BINs and clusters were included in the analysis if they possessed at least 20 records and were found in at least two geographic regions meeting the requirements of Majoros et al. (2023). They also needed to be represented by at least 10 records in each region they were found in. The species represented by these BINs and clusters were chosen and trait data was obtained. Additional species from Greenland were chosen based on occurrence records from GBIF (GBIF.org (June 24th, 2021)) GBIF Occurrence Download (https://doi.org/10.15468/dl.mk52hp) and included in the dataset. The traits were assigned to taxa using a series of literature searches conducted between April 2021 and April 2024.

Sampling Description: The biological traits for each species were determined and assigned through literature searches conducted from April 2021 - April 2024. Through the Omni Academic search tool available through the University of Guelph and Google Scholar, traits were found using the following search terms: trait AND “Taxonomic name”, habitat AND “Taxonomic name”, diet AND “Taxonomic name”, “Feeding mode” AND “Taxonomic name”, biology AND “Taxonomic name”, “Natural history” AND “Taxonomic name”, “Life history” AND “Taxonomic name”, taxonomy AND “Taxonomic name”, catalogue AND “Taxonomic name”, and “Field guide” AND “Taxonomic name”. The title and DOI of each reference are included in the dataset, and the full references are available in Appendix A. Traits were assigned to the lowest taxonomic level possible; however, not all traits could be assigned at the species level. For these species, traits were assigned using data from the next lowest level available, whether genus, subfamily or family. For example, if the authors of a given study mentioned that a particular genus has a specific trait, then species belonging to that genus were assigned that trait. The taxonomic level of the trait data is included in the dataset.

For habitat, species were classified as terrestrial (defined as taxa that live primarily in land habitats), aquatic (defined as taxa that live primarily in water bodies or associated habitats), or semi-aquatic (defined as species that primarily live in wet habitats and require high levels of moisture and some elements of both terrestrial and aquatic habitats). These assignments were made based on the larval habitat requirements. For adult and larval diet, species could be classified as predaceous (taxa who prey on insects or other animals), mycophagous (those who feed on fungi), saprophagous (those who feed on decaying organic matter), nectar/pollen/honeydew feeding (taxa that primarily feed on nectar, pollen and/or honeydew), parasitic (those who live and feed in or on an organism of another species), parasitoid (organism whose young develop within a host, eventually resulting in host death), leaf/root/stem feeding (those who feed on the leaves, roots and/or stems of plants), detritus and algae feeding (those who feed on small particles of algae and detritus), kleptoparasitic (those who take food and resources from another species), polyphagous (those who engage in more than three of the other feeding modes), nonfeeding (those who do not feed at the specified life stage), and unclear (taxa for which there is not enough information to make a trait assignment). Taxa can also be assigned a combination of the diet categories. Many higher taxonomic levels are diverse and contain species with different traits. In this case, the most common category was used. When this was unclear, multiple categories were included separated by “or”.

To check the trait assignments for accuracy and consistency, the dataset was sent to another researcher for review. He randomly selected 20 species from the dataset and performed a literature review to ensure that the literature aligned with the trait assignment.

### Geographic Coverage

Description: This dataset includes Diptera species found in Canada and Greenland that also have multiple occurrence records and DNA barcode sequences publicly available on BOLD.

### Taxonomic Coverage

Description: The dataset contains traits for 981 species across 380 genera, and 61 families. Traits are also provided for 34 subfamilies. The families are represented by varying numbers of species within the dataset (Fig. 1). For several families, the dataset only contains information for one species. The most species-rich family in this dataset is Chironomidae, which is represented by 182 species.

**Figure 1:**
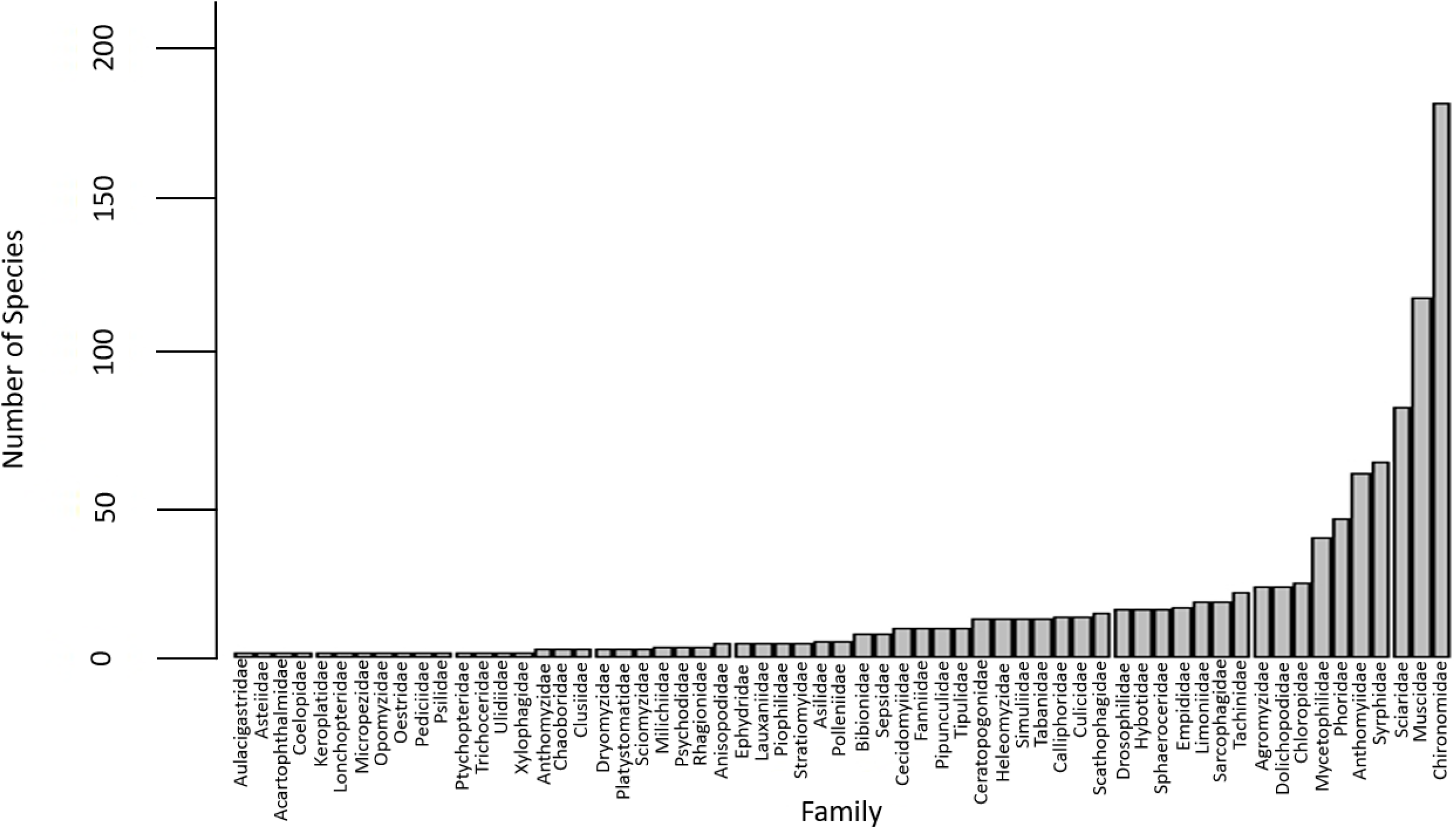
The number of Diptera species from Canada and Greenland of each family that is present in the dataset. The selected species included here also have multiple occurrence records and DNA barcode sequences publicly available on BOLD. For several small families, only one species is included. Chironomidae is the largest family within the dataset, with 182 species present.

### Trait Coverage

Description: The dataset contains information for three biological traits: habitat, adult diet, and larval diet. For habitat, 656 species were terrestrial, 277 were aquatic, and 58 were semi-aquatic (Fig 2a). For adult diet, 30 were saprophagous, 300 were nectar/pollen/honeydew feeding, 116 were predaceous, 28 were leaf/root/stem feeding, 5 were parasitic, 1 was polyphagous, 3 were detritus and algae feeding, 1 was kleptoparasitic, 18 were both nectar/pollen/honeydew feeding and predaceous, 4 were both nectar/pollen/honeydew feeding and mycophagous, 100 were both nectar/pollen/honeydew feeding and saprophagous, 56 were nectar/pollen/honeydew feeding and parasitic, 3 were nectar/pollen/honeydew feeding and kleptoparasitic, 1 was nectar/pollen/honeydew feeding, saprophagous and parasitic, 304 were non-feeding, and 21 were unclear. (Fig. 2b). For larval diet, 255 were saprophagous, 245 were predaceous, 59 were mycophagous, 5 were parasitic, 46 were parasitoids, 1 was kleptoparasitic, 98 were leaf/root/stem feeding, 175 were detritus and algae feeding, 9 were leaf/root/stem feeding and mycophagous, 9 were leaf/root/stem feeding and saprophagous, 18 were leaf/root/stem feeding and detritus and algae feeding, 1 was leaf/root/stem feeding or detritus and algae feeding, 1 was predaceous and saprophagous, 4 were detritus and algae feeding and predaceous, 10 were mycophagous and saprophagous, 10 were predaceous or saprophagous, 4 were predaceous or parasitoids, 24 were leaf/root/stem feeding or saprophagous, 15 were leaf/root/stem feeding, mycophagous and saprophagous, and 2 were unclear (Fig. 2c).

**Figure 2:**
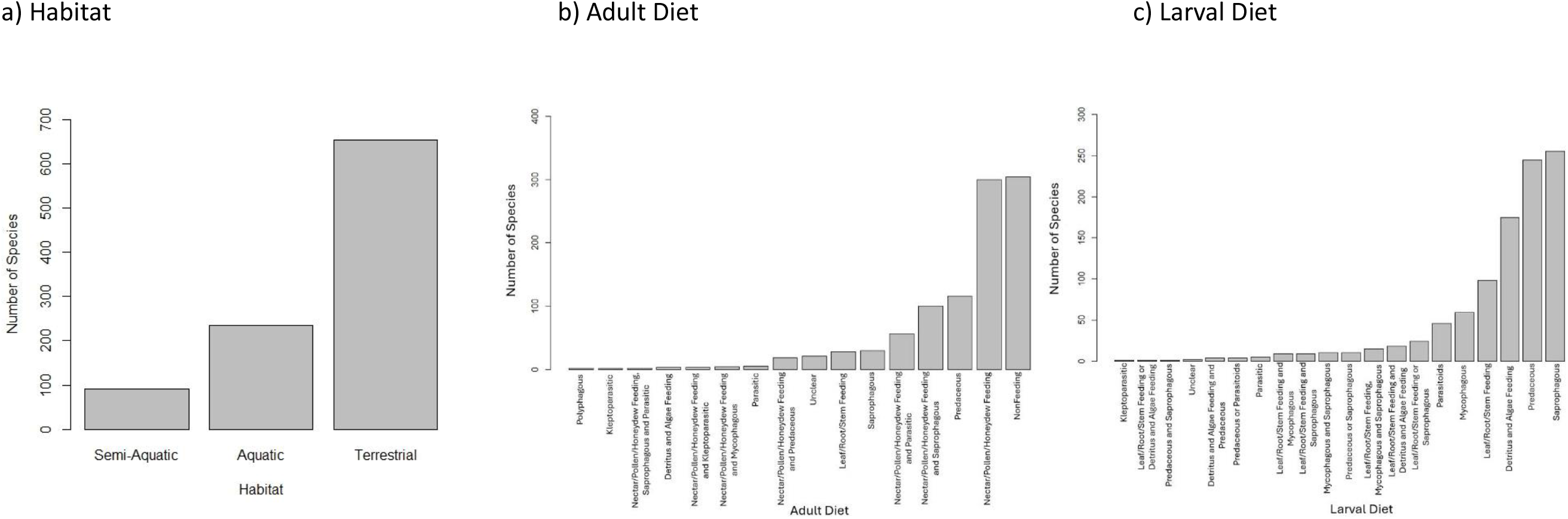
The number of Diptera species from Canada and Greenland in this dataset having each trait category for: a) habitat, b) adult diet, and c) larval diet. The majority of species included in the dataset are terrestrial. Most species are non-feeding at the adult life stage, and few are polyphagous or kleptoparasitic as adults. The majority of species included in the dataset are saprophagous as larva, and few are kleptoparasitic.

### Temporal Coverage

#### Notes

The literature searches were conducted from April 2021 to April 2024. The papers referenced were published from 1889 – 2023.

### Usage licence

Usage licence: Creative Commons Public Domain Wavier (CC-Zero)

#### Data Resources

Data package title: CanFlyet: Habitat Zone and Diet Trait Dataset for Diptera Species of Canada and Greenland

#### Number of data sets: 1

Data set name: CanFlyet: Habitat Zone and Diet Trait Dataset for Diptera Species of Canada and Greenland

Data format: csv, xlsx, tsv

Data format version: xlsx is version 2302. Csv and tsv are UTF-8.

#### Description

The dataset presents habitat, adult diet, and larval diet for 981 species across 380 genera, 25 subfamilies, and 61 families. A table showing the column labels and descriptions is provided in Table 2. The Diptera species included are from Canada and Greenland and have multiple occurrence records and DNA sequence data on BOLD. The traits were assigned to taxa using a series of literature searches conducted between April 2021 and April 2024. This dataset can be used by researchers to determine biological traits for Diptera species and conduct research that involves ecological trait data.

**Table 2:** The column labels and descriptions included within the dataset.

| Column Label | Description |
| --- | --- |
| id | The serial number of the record |
| order | The order level taxonomic classification |
| family | The family level taxonomic classification |
| subfamily | The subfamily level taxonomic classification |
| genus | The genus level taxonomic classification |
| specificEpithet | The species level taxonomic classification |
| scientificName | The scientific name of the species |
| habitat | The habitat in which the taxon is found. Taxa can be assigned terrestrial (taxa that live primarily in land habitats), aquatic (taxa that live primarily in water bodies or associated habitats) or semi-aquatic (taxa that primarily live in wet habitats and require high levels of moisture and some elements of both terrestrial and aquatic habitats). |
| traitName | The name of the biological trait. The traits included are adultDiet and larvalDiet. |
| traitValue | The trait category assigned to the species. Taxa can be assigned Predaceous (taxa who prey on insects or other animals), Mycophagous (those who feed on fungi), Saprophagous (those who feed on decaying organic matter), Nectar/Pollen/Honeydew Feeding (taxa that primarily feed on nectar, pollen and/or honeydew), Parasitic (those who live and feed in or on an organism of another species), Parasitoid (organism whose young develop within a host, eventually resulting in host death), Leaf/Root/Stem Feeding (those who feed on the leaves, roots and/or stems of plants), Detritus and Algae Feeding (those who feed on small particles of algae and detritus), Kleptoparasitic (those who take food and resources from another species), Polyphagous (those who engage in more than three of the other feeding modes), NonFeeding (those who do not feed at the specified life stage), and Unclear (taxa for which there is not enough information to make a trait assignment). The taxa can also be assigned a combination of these categories. In this case, the categories are separated by "and". For example, "Saprophagous and Predaceous". For cases where it is unclear which diet is more common, the categories are separated by "or". For example, "Saprophagous or Predaceous". |
| traitAssignmentLevelHabitat | The taxonomic level from which the habitat assignment was obtained and assigned. |
| traitAssignmentLevelAdultDiet | The taxonomic level from which the adult diet assignment was obtained and assigned. |
| traitAssignmentLevelLarvalDiet | The taxonomic level from which the larval diet assignment was obtained and assigned. |
| notes | Additional information on the habitat and diet of the taxa. Information is provided for the lowest taxonomic level used for the trait assignments if available. |
| basisOfRecord | Where the trait data was obtained from. |
| referenceNumber | The number of sources referenced when obtaining the trait information for each taxon. |
| references | The title and DOI of the reference used for the trait assignment. Full references are included in the manuscript. |

### Additional Information

For this study, we created a dataset containing habitat and adult and larval diet for 981 Diptera species across 61 families found in Canada and Greenland. This dataset allows for assignment of traits to a large variety of species. This is a valuable resource for bioinformatics work, as well as for ecological studies. This dataset was originally used to determine the relationship between biological traits and population genetic structure (Majoros et al., 2023) and can be applied to a wide range of other studies and research. These traits can also be linked to other types of data, such as species occurrence or community composition data.

The species included in this dataset possessed a wide variety of different habitat and dietary requirements. Most species were terrestrial, non-feeding at the adult life stage or saprophagous at the larval stage. Other diets were far less represented, such as those with a kleptoparasitic life stage.

Trait data has been used to investigate various ecological and evolutionary patterns and topics, such as community assembly (Kraft & Ackerly, 2010, Zhang et al., 2020), molecular evolution (May et al., 2020), phylogenetic assemblage structure (Barnagaud et al., 2014), and how populations change over time or in response to ecological change (Coulthard et al., 2019, De Palma et al., 2015). Similarly, this dataset can be used to answer a variety of questions and is a useful tool for future research. It can be used for various studies that relate to ecology, evolution, biogeography, and conservation of fly species and include the habitat and diet of flies. The dataset can be used to investigate the relationship between traits and population genetic structure or phylogenetic community structure. The dataset could also be used to find relationships between traits, such as in studies like Freire et al. (2021). A visualization is provided in Figure 3 of the phylogenetic relationships between species possessing different traits. This could be built upon to look at the relationship between traits and evolutionary history and to investigate the phylogenetic relationships, such as in studies like Rainford & Mayhew (2015). The trait dataset can also be used to investigate the diversity of different fly traits in an area of interest, to determine how these traits impact occurrence patterns, how traits may relate to responses to environmental change, or to determine which species may be most likely to colonize these areas in the future. Trait data could also prove useful for biomonitoring of insect diversity in agricultural areas, such as research into which farming practices are associated with a balance of ecological functions in associated insect communities.

**Figure 3:**
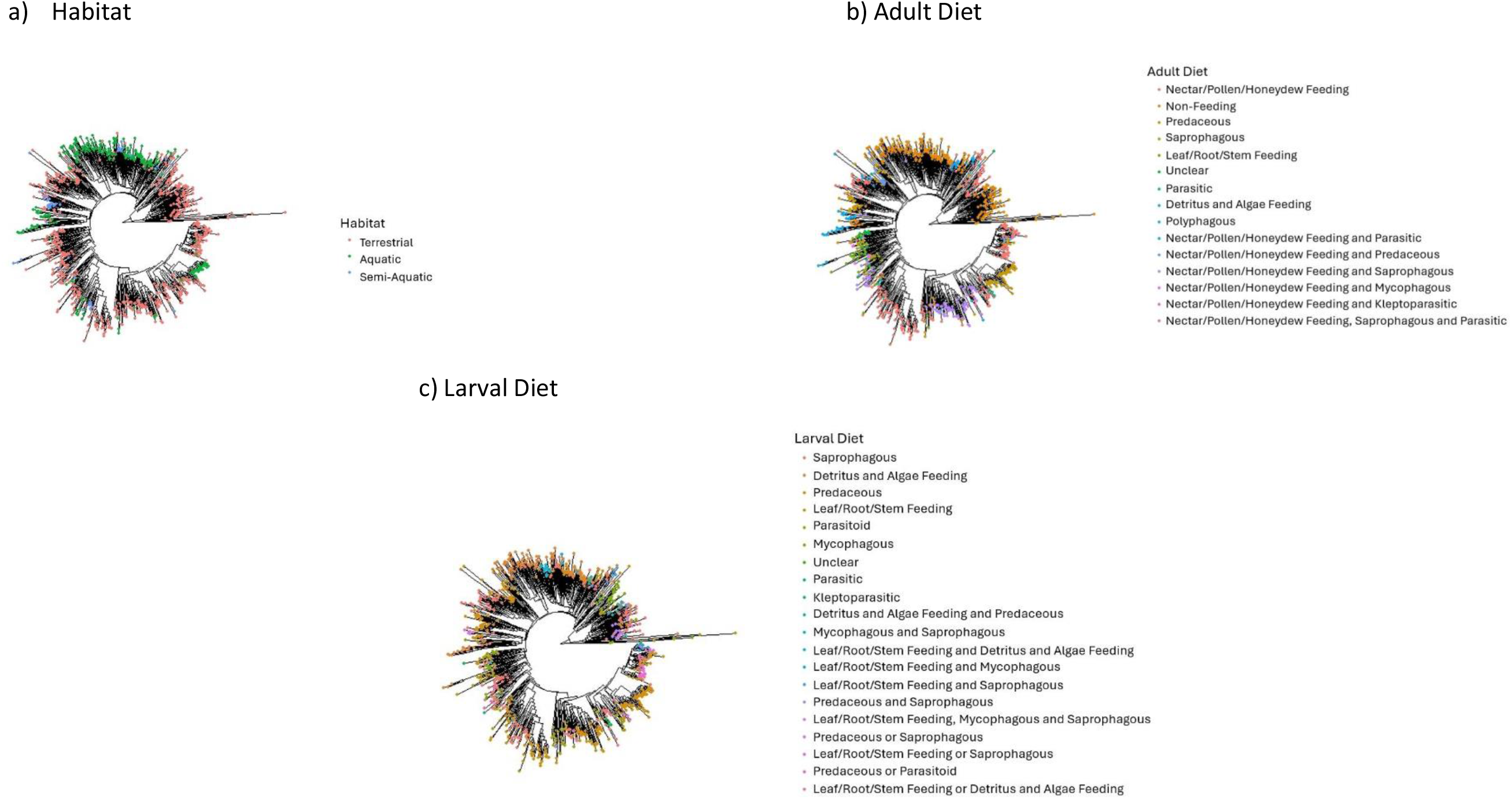
Phylogenetic trees of Diptera species from Canada and Greenland included in this dataset; these species also have multiple occurrence records and DNA barcode sequences publicly available on BOLD. The different colours of the tips represent species with different: a) habitat categories, b) adult diet categories, and c) larval diet categories. The traits provided in the dataset could be used to investigate the relationship between traits and evolutionary history. These maximum likelihood trees were created using cytochrome c oxidase subunit 1 (CO1) sequences from BOLD and functions from the package phangorn version 2.11.1(Schliep, 2011) in the R programming language (R Core Team, 2023). The tips were coloured using the package ggtree version 3.6.2 (Yu et al., 2017). The code for how to create this tree is provided at https://github.com/S-Majoros/Diptera_Dataset_Phylogenetic_Tree

In this paper we presented a trait dataset containing habitat and diet information for a large number of Diptera taxa from Canada and Greenland that also have occurrence records and DNA barcoding sequences available on BOLD. This dataset is publicly available on Data Dryad (https://datadryad.org/stash/dataset/doi:10.5061/dryad.fqz612jwx), and is also available at on GitHub (https://github.com/S-Majoros/Diptera_Dataset_Phylogenetic_Tree). The file is available in csv, xlsx, and tsv format in order to make it as accessible as possible for future users. This dataset is publicly available so that other researchers can use it to conduct further studies, answer more questions, and improve our knowledge of this ecologically important group.

## Acknowledgements

The authors acknowledge financial support from the Natural Sciences and Engineering Research Council of Canada (NSERC); the Government of Canada through Genome Canada and Ontario Genomics; the Ontario Ministry of Economic Development, Job Creation and Trade; and Food from Thought: Agricultural Systems for a Healthy Planet Initiative program funded by the Government of Canada through the Canada First Research Excellence Fund. We acknowledge and thank Karl Cottenie for additional funding provided through an NSERC Discovery Grant. The authors would like to thank Tyler Elliott for reviewing the manuscript, quality checking the dataset and providing helpful insight. The authors would also like to thank the researchers from whose work these trait data were gathered, as well as all of the researchers who contribute publicly available data to databases such as BOLD.

## Author Contributions

Samantha E. Majoros – project design, dataset construction, manuscript writing and editing. Sarah J. Adamowicz – project design, manuscript editing.

