## Appendix A for "CanFlyet: Habitat Zone and Diet Trait Dataset for Diptera Species of Canada and Greenland"

**APPENDICES**

Appendix A: References for Diptera habitat and diet traits

Acartophthalmidae

Roháček, J. (2012). The fauna of the opomyzoid families Clusiidae, Acartophthalmidae, Anthomyzidae, Opomyzidae, Stenomicridae, Periscelididae, Asteiidae (Diptera) in the Gemer area (Central Slovakia). *Časopis Slezského Zemského Muzea, 61*(2), 97-111. doi: 10.2478/v10210-012-0011-5

Roháček, J. (2013). The fauna of the Acalyptrate families Micropezidae, Psilidae, Clusiidae, Acartophthalmidae, Anthomyzidae, Aulacigastridae, Periscelididae and Asteiidae (Diptera) in the Gemer area (Central Slovakia): supplement 1. *Casopis Slezskeho Zemskeho Muzea, 62*(2), 125-136. doi: 10.2478/cszma-2013-0014

Agromyzidae

Ayabe, Y. (2010). Specific Mining Pattern as a Result of Selective Feeding Within a Leaf by the Dipteran Leafminer *Ophiomyia maura* (Diptera: Agromyzidae). *Annals of the Entomological Society of America, 103*(5), 806–812. https://doi.org/10.1603/AN10049

Ayabe, Y., Ueno, T. & Harvey, J.A. (2012). Complex feeding tracks of the sessile herbivorous insect *Ophiomyia maura* as a function of the defense against insect parasitoids. *PLoS one, 7*(2), p.e32594-e32594. doi: https://doi.org/10.1371/journal.pone.0032594

Bañuelos, M. & Kollmann, J. (2011). Effects of host-plant population size and plant sex on a specialist leaf-miner. *Acta Oecologica, 37*(2), 58-64. doi: https://doi.org/10.1016/j.actao.2010.11.007

Beri, S.K. (2007). Biology of a leaf miner *Liriomyza brassicae* (Riley) (Diptera : Agromyzidae). *Journal of Natural History, 8*(2), 143-151. doi: https://doi.org/10.1080/00222937400770101

Černý, M., Andrade, R., Gonçalves, A.R. & von Tschirnhaus, M. (2018). New records of Agromyzidae (Diptera) from Portugal, with an updated checklist. *Acta musei silesiae : scientiae naturales, 67*(1), 7-57. doi: https://doi.org/10.2478/cszma-2018-0002

Černý, M., Kocián, J. & Ševčík, J. (2021). First rearing record and DNA sequence of *Chromatomyia aizoon* (Hering, 1932) (Diptera: Agromyzidae) associated with *Saxifraga paniculata* Miller from the Czech Republic. *Acta Musei Silesiae, Scientiae Naturales, 69*(3), 283-288. doi: https://doi.org/10.2478/cszma-2020-0021

Cohen, M. (1936). The biology of the chrysanthemum leaf-miner, *Phytomyza atricornis* mg. (Diptera: Agromyzide). *Annals of Applied Biology, 23*(3), 612-632. doi: https://doi.org/10.1111/j.1744-7348.1936.tb06114.x

Day, M.D., Riding, N. & Chamberlain, A. (2009). Biology and host range of Ophiomyia camarae Spencer (Diptera: Agromyzidae), a potential biocontrol agent for Lantana spp. (Verbenaceae) in Australia. *Biocontrol Science and Technology, 19*(6), 627-637. https://doi.org/10.1080/09583150902968980

Eber, S. (2004). Bottom-up density regulation in the holly leaf-miner *Phytomyza ilicis*. *The Journal of Animal Ecology, 73*(5), 948-958. doi: https://doi.org/10.1111/j.0021-8790.2004.00867.x

Ellis, W.N. (2020). *Parasites*. Plant Parasites of Europe. https://bladmineerders.nl/parasites/

Gratton, C. & Welter, S.C. (1998). Oviposition preference and larval performance of *Liriomyza helianthi* (Diptera: Agromyzidae) on normal and novel host plants. *Environmental Entomology, 27*(4), 926-935. doi: https://doi.org/10.1093/ee/27.4.926

Griswold, G.H. (1928). A New Leaf Miner Injurious to Larkspur (*Phytomyza delphiniae* Frost). *Journal of Economic Entomology, 21*(6), 855–857. doi: https://doi.org/10.1093/jee/21.6.855

Hendrickson, Jr., R.M. & Barth, S.E. (1978). Biology of the Alfalfa Blotch Leafminer. *Annals of the Entomological Society of America, 71*(3), 295-298. doi: https://doi.org/10.1093/aesa/71.3.295

Hill, R.L., Wittenberg, R. & Gourlay, A.H. (2010). Biology and Host Range of *Phytomyza vitalbae* and its Establishment for the Biological Control of *Clematis vitalba* in New Zealand. *Biocontrol Science and Technology, 11*(4), 459-473. doi: https://doi.org/10.1080/09583150120067490

James, R. & Pritchard, I.M. (2007). Influence of the holly leaf miner, *Phytomyza ilicis* (Diptera Agromyzidae), on leaf abscission. *Journal of Natural History, 22*(2), 395-402. doi: https://doi.org/10.1080/00222938800770281

Kaurava, A.S., Dhamdhere, S.V. & Odak, S.C. (1970). Studies on the Biology of *Phytomyza Atricornis* Meigen (Agromyzidae: Diptera). *The Journal of the Bombay Natural History Society, 67*, 597-604.

Lambkin, C.L., Fayed, S.A., Manchester, C., La Salle, J., Scheffer, S.J. & Yeates, D.K. (2008). Plant hosts and parasitoid associations of leaf mining flies (Diptera: Agromyzidae) in the Canberra region of Australia. *Australian Journal of Entomology, 47*(1), 13-19. doi: https://doi.org/10.1111/j.1440-6055.2007.00622.x

Lonsdale, O. (2011). The Liriomyza (Agromyzidae: Schizophora: Diptera) of California. *Zootaxa, 2850*(1), 1-123. doi: https://doi.org/10.11646/zootaxa.2850.1.1

Lonsdale, O. (2021). Manual of North American Agromyzidae (Diptera, Schizophora), with revision of the fauna of the “Delmarva” states. *ZooKeys, 1051*, 1-481. doi: https://doi.org/10.3897/zookeys.1051.64603

Marino, P.C. & Cornell, H.V. (1992). Adult movement of the native holly leafminer, *Phytomyza ilicicola* Loew (Diptera: Agromyzidae): Consequences for host choice within and between habitats. *Oecologia, 92*(1), 76-82. doi: https://doi.org/10.1007/BF00317265

Nzama, S., Olckers, T. & Zachariades, C. (2014). Seasonal activity, habitat preferences and larval mortality of the leaf-mining fly *Calycomyza eupatorivora* (Agromyzidae), a biological control agent established on *Chromolaena odorata* (Asteraceae) in South Africa. *Biocontrol Science and Technology, 24*(11), 1297-1307. Doi: 10.1080/09583157.2014.935293

Nzama, S., Olckers, T. & Zachariades, C. (2014). Is oviposition and larval damage by the leaf-mining fly Calycomyza eupatorivora (Agromyzidae) on its target weed, Chromolaena odorata (Asteraceae), restricted by leaf-quality preferences? *Biocontrol Science and Technology, 24*(6), 680-689. https://doi.org/10.1080/09583157.2014.889658

Scheffer, S.J. & Lonsdale, O. (2011). *Phytomyza omlandi* spec. nov. - The First Species of Agromyzidae (Diptera: Schizophora) Reared from the Family Gelsemiaceae (Asteridae). *Proceedings of the Entomological Society of Washington, 113*(1), 42-49. https://doi.org/10.4289/0013-8797.113.1.42

Scheffer, S.J., Lewis, M.L., Hébert, J.B. & Jacobsen, F. (2020). Diversity and host plant-use North American *Phytomyza* holly leafminers (Diptera: Agromyzidae): colonization, divergence, and specificity in a host-associated radiation. *Annals of the Entomological Society of America, 114*(1), 56-69. doi: https://doi.org/10.1093/aesa/saaa03

Scheffer, S.J. & Lonsdale, O. (2011). *Phytomyza omlandi* spec. nov. – The first species of Agromyzidae (Diptera: Schizophora) reared from the family Gelsemiaceae (Asteridae). *Proceedings of the Entomological Society of Washington, 113*(1), 42-49. doi: 10.4289/0013-8797.113.1.42

Scheffer, S.J. & Wiegmann, B.M. (2000). Molecular phylogenetics of the holly leafminers (Diptera: Agromyzidae: *Phytomyza*): species limits, speciation, and dietary specialization. *Molecular Phylogenetics and Evolution, 17*(2), 244-255. doi: https://doi.org/10.1006/mpev.2000.0830

Scudder, G.G.E. & Cannings, R.A. (2006). The Diptera families of British Columbia. http://www.for.gov.bc.ca/hfd/library/FIA/2006/FSP_Y062001b.pdf

Sehgal, V.K. (1971). A Taxonomic Survey of the Agromyzidae (Diptera) of Alberta, Canada, with Observations on Host-Plant Relationships. *Quaestiones entomologicae, 7*, 291-405.

Simelane, D.O. (2002). Biology and host range of Ophiomyia camarae, a biological control agent for Lantana camara in South Africa. *BioControl, 47*(5), 575-585. https://doi.org/10.1023/A:1016541809545

Smith, P.H.D. & Jones, T.H. (2002). Effects of elevated CO2 on the chrysanthemum leaf-miner, Chromatomyia syngenesiae: a greenhouse study. *Global Change Biology, 4*(3), 287-291. https://doi.org/10.1046/j.1365-2486.1998.00149.x

Soltani, A., Ben Abda, M., Amri, M., Carapelli, A. & Mediouni Ben Jemâa, J. (2020). Seasonal incidence of the leaf miner *Liriomyza cicerina* Rond (Diptera: Agromyzidae) in chickpea fields and effects of climatic parameters, chickpea variety, and planting date on the leaf miner infestation rate. *Euro-Mediterranean journal for environmental integration, 5*(3). doi: https://doi.org/10.1007/s41207-020-00198-4

Spencer, K.A. (1987). Agromyzidae. In McAlpine, J.F., Peterson, B.V., Shewell, G.E., Teskey, H.J., Vockeroth, J.R. & Wood, D.M. (Eds), Manual of Nearctic Diptera. Volume 2. (pp. 869 – 880). Research Branch Agriculture Canada.

Talekar, N.S. & Lee, Y.H. (1988). Biology of *Ophiomyia centrosematis* (Diptera: Agromyzidae), a pest of soybean. *Annals of the Entomological Society of America, 81*(6), 938-942. https://doi.org/10.1093/aesa/81.6.938

Tauber, M.J. & Tauber, C.A. (2012). Biology and leaf-mining behaviour of *Phytomyza lanati* (Diptera: Agromyzidae). *The Canadian Entomologist, 100*(4), 341-349. doi: https://doi.org/10.4039/Ent100341-4

Williams, G.L. & Steck, G.J. (2015). *Ophiomyia kwansonis* Sasakawa (Diptera: Agromyzidae), the Daylily Leafminer, an Asian Species Recently Identified in the Continental United States. *Proceedings of the Entomological Society of Washington, 116*(4), 421-428. https://doi.org/10.4289/0013-8797.116.4.421

Anisopodidae

Peterson, B.V. (1981). Anisopodidae. In McAlpine, J.F., Peterson, B.V., Shewell, G.E., Teskey, H.J., Vockeroth, J.R. & Wood, D.M. (Eds), Manual of Nearctic Diptera. Volume 1. (pp. 305 – 312). Research Branch Agriculture Canada.

Scudder, G.G.E. & Cannings, R.A. (2006). The Diptera families of British Columbia. http://www.for.gov.bc.ca/hfd/library/FIA/2006/FSP_Y062001b.pdf

Anthomyiidae

Ackland, D.M. (2008). Revision of Afrotropical Delia Robineau-Desvoidy, 1830 (Diptera: Anthomyiidae), with Descriptions of six New Species. *African Invertebrates, 49*(1), 1-75. doi: https://doi.org/10.5733/afin.049.0101

Bažok, R., Ceranić-Sertić, M., Barčić, J.I., Borošić, J., Kozina, A., Kos, T., Lemić, D. & Čačija, M. (2012). Seasonal flight, optimal timing and efficacy of selected insecticides for cabbage maggot (*Delia radicum* L., Diptera: Anthomyiidae) control. *Insects, 3*(4), 1001-1027. doi: https://doi.org/10.3390/insects3041001

Bland, K.P. & Godfray, H.C.J. (1992). New foodplants for two species of leaf-mining *Pegomya* (Diptera: Anthomyiidae) in Britain. *British journal of entomology and natural history, 5*, 127-128.

Bosnyákné, H.E., Kerepesi, I. & Keszthelyi, S. (2016). New insight into the *Delia platura* Meigen caused alteration in nutrient content of soybean (*Glycine max* L. Merill). *Acta Biologica Hungarica, 67*(3), 261-268. doi: https://doi.org/10.1556/018.67.2016.3.4

Bultman, T.L. & Leuchtmann, A. (2008). Biology of the *Epichloë–Botanophila* interaction: An intriguing association between fungi and insects. *Fungal Biology Reviews, 22*(3-4), 131-138. doi: https://doi.org/10.1016/j.fbr.2009.04.003

Bultman, T.L., Leuchtmann, A., Sullivan, T.J. & Dreyer, A.P. (2011). Do *Botanophila* flies provide reproductive isolation between two species of *Epichloë* fungi? A field test. *New Phytologist, 190*(1), 206-212. doi: https://doi.org/10.1111/j.1469-8137.2010.03612.x

Córdova-García, G., Navarro-de-la-Fuente, L., Pérez-Staples, D., Williams, T. & Lasa, R. (2023). Biology and Ecology of *Delia planipalpis* (Stein) (Diptera: Anthomyiidae), an Emerging Pest of Broccoli in Mexico. *Insects, 14*(7), 659. doi: https://doi.org/10.3390/insects14070659

Ellis, W.N. (2020). *Parasites.* Plant Parasites of Europe. https://bladmineerders.nl/parasites/

Gomes, L.R.P. & de Carvalho, C.J.B. (2021). Three new species of Botanophila Lioy (Diptera: Anthomyiidae) from the Mexican Transition Zone. *Zootaxa, 5005*(3), 317-328. doi: https://doi.org/10.11646/zootaxa.5005.3.6

Gomes, L.R.P. & de Carvalho, C.J.B. (2022). Taxonomy of the Neotropical species of *Calythea* (Anthomyiidae: Diptera), with description of two new species from South America. *Revista Brasileira de Entomologia, 66*(1), 1. doi: https://doi.org/10.1590/1806-9665-RBENT-2021-0102

Gomes, L.R.P., Couri, M.S. & de Carvalho, C.J.B. (2018). Anthomyiidae, Fanniidae and Muscidae (Diptera) from the Juan Fernández Archipelago (Chile): 60 years after Willi Hennig's contributions. *Zootaxa, 4402*(2), 373-389. doi: https://doi.org/10.11646/zootaxa.4402.2.9

Griffiths, K. & Stewart, D. (2004). New Record and Phylogenetic Analysis of *Anthomyia pluvialus* in Nova Scotia. *Northeastern Naturalist, 11*(2), 189-19. doi: https://doi.org/10.1656/1092-6194(2004)011[0189:NRAPAO]2.0.CO;2

Guerra, P.C., Keil, C.B., Stevenson, P.C., Mina, D., Samaniego, S., Peralta, E., Mazon, N. & Chancellor, T.C.B. (2016). Larval performance and adult attraction of *Delia platura* (Diptera: Anthomyiidae) in a native and an introduced crop. *Journal of Economic Entomology, 110*(1), 186-191. doi: https://doi.org/10.1093/jee/tow237

Huckett, H.C. (1987). Anthomyiidae. In McAlpine, J.F., Peterson, B.V., Shewell, G.E., Teskey, H.J., Vockeroth, J.R. & Wood, D.M. (Eds), Manual of Nearctic Diptera. Volume 2. (pp. 1099 – 1114). Research Branch Agriculture Canada.

Johansen, T.J. (1990). On the biology of *Delia fabricii* Holm. (Dipt., Anthomyiidae) in Northern Norway. *Journal of Applied Entomology, 110*(1-5), 454-461. doi: https://doi.org/10.1111/j.1439-0418.1990.tb00145.x

Kutty, S.N., Bernasconi, M.V., Šifner, F. & Meier, R. (2007). Sensitivity analysis, molecular systematics and natural history evolution of Scathophagidae (Diptera: Cyclorrhapha: Calyptratae). *Cladistics, 23*(1), 64-83. doi: https://doi.org/10.1111/j.1096-0031.2006.00131.x

Lamb, R.J. & Boivin, G. (2018). Population variability of three Delia species (Diptera: Anthomyiidae) from the same agricultural habitat in Quebec, Canada. *Canadian Entomologist, 150*(1), 80-96. doi: https://doi.org/10.4039/tce.2017.52

Leuchtmann, A. (2007). Botanophila flies on Epichloë host species in Europe and North America: no evidence for co-evolution. *Entomologia Experimentalis et Applicata, 123*(1), 13-23. doi: https://doi.org/10.1111/j.1570-7458.2006.00518.x

Mesmin, X., Vincent, M., Tricault, Y., Estorgues, V., Daniel, L., Cortesero, A., Faloya, V. & Le Ralec, A. (2019). Assessing the relationship between pest density and plant damage: a case study with the belowground herbivore *Delia radicum* (Diptera: Anthomyiidae) on broccoli. *Applied Entomology and Zoology, 54*(2), 155-165. doi: https://doi.org/10.1007/s13355-019-00607-3

Michelsen, V. (2007). Taxonomic review of Eurasian *Paradelia* Ringdahl (Diptera: Anthomyiidae) with descriptions of two new species. *Zootaxa, 1592*, 1-44.

Michelsen, V. (2009). Taxonomic revision of the *Pegomya meridiana* species group (Diptera: Anthomyiidae) including natural enemies of invasive *Hypericum* spp. (Clusiaceae). *Zootaxa, 2299*(1), 29-43. doi: https://doi.org/10.11646/zootaxa.2299.1.3

Michelsen, V. (2015). Taxonomic review of the major larval pests of bolete fungi (Boletaceae) in Europe: The *Pegomya fulgens*, furva and *tabida* species groups (Diptera: Anthomyiidae). *Zootaxa, 4020*(1), 51-80. doi: http://dx.doi.org/10.11646/zootaxa.4020.1.2

Michelsen, V. & Palmer, M.W. (2020). *Pegomya disticha* Griffiths and *P. cedrica* Huckett (Diptera: Anthomyiidae)—first documented case of insects trespassing the silica barrier of Common scouring-rush, *Equisetum hyemale* L. *Zootaxa, 4718*(3), 355-370. doi: https://doi.org/10.11646/zootaxa.4718.3.4

Mlynarek, J.J., MacDonald, M., Sim, K., Hiltz, K., McDonald, M.R. & Blatt, S. (2020). Oviposition, feeding preferences and distribution of *Delia* species (Diptera: Anthomyiidae) in Eastern Canadian onions. *Insects, 11*(11), 780. doi: https://doi.org/10.3390/insects11110780

Nilsson, U., Rännbäck, L.M., Anderson, P. & Rämert, B. (1986). Herbivore response to habitat manipulation with floral resources: a study of the cabbage root fly. *Journal of Applied* Entomology, 136(7), 481-489. doi: https://doi.org/10.1111/j.1439-0418.2011.01685.x

Rao, S. & Baumann, D. (2004). The interaction of a Botanophila fly species with an exotic Epichloë fungus in a cultivated grass: fungivore or mutualist? *Entomologia Experimentalis et Applicata, 112*(2), 99-105. doi: https://doi.org/10.1111/j.0013-8703.2004.00189.x

Rao, S., Alderman, S.C., Takeyasu, J. & Matson, B. (2005). Botanophila-Epichloe association in cultivated Festuca in Oregon: evidence of simple fungivory. *Entomologia Experimentalis et Applicata, 115*(3), 427-433. doi: https://doi.org/10.1111/j.1570-7458.2005.00309.x

Rotheray, G. & Lyszkowski, R. (2015). Diverse mechanisms of feeding and movement in Cyclorrhaphan larvae (Diptera). *Journal of Natural History, 49*(35-36), 2139-2211. doi: https://doi.org/10.1080/00222933.2015.1010314

Scudder, G.G.E. & Cannings, R.A. (2006). The Diptera families of British Columbia. http://www.for.gov.bc.ca/hfd/library/FIA/2006/FSP_Y062001b.pdf

Vitou, J., Briese, D.T., Sheppard, A.W. & Thomann, T. (2008). Comparative biology of two rosette crown-feeding flies of the genus *Botanophila* (Dipt., Anthomyiidae) with potential for biological control of their thistle hosts. *Journal of Applied Entomology, 125*(1-2), 89-95. doi: https://doi.org/10.1111/j.1439-0418.2001.00495.x

Anthomyzidae

Ellis, W.N. (2020). *Parasites.* Plant Parasites of Europe. https://bladmineerders.nl/parasites/

Roháček, J. (2012). The fauna of the opomyzoid families Clusiidae, Acartophthalmidae, Anthomyzidae, Opomyzidae, Stenomicridae, Periscelididae, Asteiidae (Diptera) in the Gemer area (Central Slovakia). *Časopis Slezského Zemského Muzea, 61*(2), 97-111. doi: https://doi.org/10.2478/v10210-012-0011-5

Roháček, J. (2013). The fauna of the Acalyptrate families Micropezidae, Psilidae, Clusiidae, Acartophthalmidae, Anthomyzidae, Aulacigastridae, Periscelididae and Asteiidae (Diptera) in the Gemer area (Central Slovakia): supplement 1. *Acta musei silesiae: scientiae naturales, 62*(2), 125. doi: https://doi.org/10.2478/cszma-2013-0014

Roháček, J. (2021). New species and records of Anthomyzidae (Diptera) from the East Palaearctic, with a checklist of taxa occurring in the area. *Acta Entomologica Musei Nationalis Pragae, 61*(1), 261-288. doi: https://doi.org/10.37520/aemnp.2021.016

Roháček, J. & Barber, K.N. (2023). Nearctic Anthomyzidae: Genera *Mumetopia* Melander and *Xerocomyza* gen. n. (Diptera). *European Journal of Entomology, 120*(1), 254-292. doi: https://doi.org/10.14411/eje.2023.028

Vockeroth, J.R. (1987). Anthomyzidae. In McAlpine, J.F., Peterson, B.V., Shewell, G.E., Teskey, H.J., Vockeroth, J.R. & Wood, D.M. (Eds), Manual of Nearctic Diptera. Volume 2. (pp. 887 – 890). Research Branch Agriculture Canada.

Asilidae

Baker, N.T. & Fischer, R.L. (1975). A Taxonomic and Ecological Study of the Asilidae of Michigan. *The Great Lakes Entomologist, 8*(2). doi: https://doi.org/10.22543/0090-0222.1248

Dennis, D.S., Barnes, J.K. & Knutson, L. (2013). Review and analysis of information on the biology and morphology of immature stages of robber flies (Diptera: Asilidae). *Zootaxa, 3673*(1), 1-64. doi: http://dx.doi.org/10.11646/zootaxa.3673.1.1

Dennis, D.S., Lavigne, R.J. & Dennis, J.G. (2012). Spiders (Araneae) as prey of robber flies (Diptera: Asilidae). *Journal of Entomological Research Society, 14*(1), 65-76.

Londt, J.G.H. (2010). A Taxonomic Analysis of Gambian Asilidae (Diptera) Based Chiefly on a Collection Assembled by W.F. Snow between 1974 and 1977. *African Entomology, 18*(2), 328-353. doi: https://doi.org/10.4001/003.018.0210

Scudder, G.G.E. & Cannings, R.A. (2006). The Diptera families of British Columbia. http://www.for.gov.bc.ca/hfd/library/FIA/2006/FSP_Y062001b.pdf

Wood, G.C. (1981). Asilidae. In McAlpine, J.F., Peterson, B.V., Shewell, G.E., Teskey, H.J., Vockeroth, J.R. & Wood, D.M. (Eds), Manual of Nearctic Diptera. Volume 1. (pp. 549 – 574). Research Branch Agriculture Canada.

### Asteiidae

Roháček, J. (2012). The fauna of the opomyzoid families Clusiidae, Acartophthalmidae, Anthomyzidae, Opomyzidae, Stenomicridae, Periscelididae, Asteiidae (Diptera) in the Gemer area (Central Slovakia). *Časopis Slezského Zemského Muzea, 61*(2), 97-111. doi: https://doi.org/10.2478/v10210-012-0011-5

Roháček, J. (2013). The fauna of the Acalyptrate families Micropezidae, Psilidae, Clusiidae, Acartophthalmidae, Anthomyzidae, Aulacigastridae, Periscelididae and Asteiidae (Diptera) in the Gemer area (Central Slovakia): supplement 1. *Acta musei silesiae: scientiae naturales, 62*(2), 125. doi: https://doi.org/10.2478/cszma-2013-0014

Sabrosky, C.W. (1987). Asteiidae. In McAlpine, J.F., Peterson, B.V., Shewell, G.E., Teskey, H.J., Vockeroth, J.R. & Wood, D.M. (Eds), Manual of Nearctic Diptera. Volume 2. (pp. 899 – 902). Research Branch Agriculture Canada.

### Aulacigastridae

Roháček, J. (2013). The fauna of the Acalyptrate families Micropezidae, Psilidae, Clusiidae, Acartophthalmidae, Anthomyzidae, Aulacigastridae, Periscelididae and Asteiidae (Diptera) in the Gemer area (Central Slovakia): supplement 1. C*asopis Slezskeho Zemskeho Muzea, 62*(2), 125-136. doi: https://doi.org/10.2478/cszma-2013-0014

Rung, A. & Mathis, W.N. (2011). A revision of the genus *Aulacigaster* Macquart (Diptera:Aulacigastridae). *Smithsonian Contributions to Zoology, 633*, 1-129. doi: https://doi.org/10.5479/si.00810282.633

Winkler, I.S., Rung, A. & Scheffer, S.J. (2010). Hennig's orphans revisited: Testing morphological hypotheses in the ''Opomyzoidea" (Diptera: Schizophora). *Molecular phylogenetics and evolution, 54*(3), 746-762. doi: http://doi.org/10.1016/j.ympev.2009.12.016

### Bibionidae

Fitzgerald, S.J. (2004). *Evolution and Classification of Bibionidae (Diptera: Bibionomorpha).* [Doctoral dissertation, Oregon State University].

Fitzgerald, S.J. (2021). *Penthetria* Meigen (Diptera: Bibionidae): Revision of the New World species and world catalog. *Zootaxa, 4926*(4), 451-500. doi: https://doi.org/10.11646/zootaxa.4926.4.1

Frouz, J., Lin, Q., Li, X., Abakumov, E., Brune, A. & Šustr, V. (2020). Utilization of dietary protein in the litter-dwelling larva of *Bibio marci* (Diptera: Bibionidae). *Eurasian Soil Science, 52*(12), 1583-1587. doi: https://doi.org/10.1134/S1064229319120032

Morris, H.M. (1917). On the larval and pupal stages of *Bibio johannis* L. *Annals of Applied Biology, 4*(3), 91-114. doi: https://doi.org/10.1111/j.1744-7348.1917.tb05907.x

Needham, J.G. (1902). A remarkable occurrence of the fly, *Bibio fraternus* Loew. The American Naturalist, 36(423), 181-185. doi: https://doi.org/10.1086/278097

Calliphoridae

Bennett, G.F. & Whitworth, T.L. (1991). Studies on the life history of some species of *Protocalliphora* (Diptera: Calliphoridae). *Canadian Journal of Zoology, 69*(8), 2048-2058. doi: https://doi.org/10.1139/z91-286

Biavati, G.M., De Assis Santana, F.H. & Pujol-Luz, J.R. (2010). A Checklist of Calliphoridae Blowflies (Insecta, Diptera) Associated with a Pig Carrion in Central Brazil. *Journal of Forensic Sciences, 55*(6), 1603-1606. doi: https://doi.org/10.1111/j.1556-4029.2010.01502.x

Coupland, J.B. & Barker, G.M. (2004). Diptera as predators and parasitoids of terrestrial gastropods, with emphasis on Phoridae, Calliphoridae, Sarcophagidae, Muscidae and Fanniidae. *Natural Enemies of Terrestrial Molluscs*, 85-158. doi: https://doi.org/10.1079/9780851993195.0085

Daoust, S.P., Savage, J., Whitworth, T.L., Bélisle, M. & Brodeur, J. (2012). Diversity and abundance of ectoparasitic blow flies *Protocalliphora* (Diptera: Calliphoridae) and their *Nasonia* (Hymenoptera: Pteromalidae) parasitoids in tree swallow nests within agricultural lands of southern Québec, Canada. *Annals of the Entomological Society of America, 105*(3), 471–478. doi: https://doi.org/10.1603/AN11155

De Jong, G.D., Meyer, F. & Goddard, J. (2021). An annotated list of the blow flies and cluster flies (Diptera: Calliphoridae, Polleniidae) of Mississippi. *Transactions of the American Entomological Society, 147*(3), 827-843. doi: https://doi.org/10.3157/061.147.0304

Dykstra, C.R., Hays, J.L., Simon, M.M. & Wegman, A.R. (2012). *Protocalliphora* (Diptera: Calliphoridae) infestations of nestling red-shouldered hawks in Southern Ohio. *The Wilson Journal of Ornithology, 124*(4), 783-787. doi: https://doi.org/10.1676/1559-4491-124.4.783

Erzinçlioğlu, Y.Z. (2007). Immature stages of British *Calliphora* and *Cynomya,* with a re-evaluation of the taxonomic characters of larval Calliphoridae (Diptera). *Journal of Natural History, 19*(1), 69-96. doi: https://doi.org/10.1080/00222938500770041

Giordani, G., Tuccia, F., Floris, I. & Vanin, S. (2018). First record of *Phormia regina* (Meigen, 1826) (Diptera: Calliphoridae) from mummies at the Sant'Antonio Abate Cathedral of Castelsardo, Sardinia, Italy. *PeerJ, 6*. doi: https://doi.org/10.7717/peerj.4176

Gold, C.S. & Dahlsten, D.L. (1989). Prevalence, habitat selection, and biology of *Protocalliphora* (Diptera: Calliphoridae) found in nests of mountain and chestnut-backed chickadees in California. *Hilgardia, 57*(2), 1-19. doi: https://doi.org/10.3733/hilg.v57n02p019

Maeda, T., Tamotsu, M., Yamaoka, R. & Ozaki, M. (2015). Effects of floral scents and their dietary experiences on the feeding preference in the blowfly, *Phormia regina*. *Frontiers in Integrative Neuroscience, 9*, 59. doi: https://doi.org/10.3389/fnint.2015.00059

Rotheray, G. & Lyszkowski, R. (2015). Diverse mechanisms of feeding and movement in Cyclorrhaphan larvae (Diptera). *Journal of Natural History, 49*(35-36), 2139-2211. doi: https://doi.org/10.1080/00222933.2015.1010314

Scudder, G.G.E. & Cannings, R.A. (2006). The Diptera families of British Columbia. http://www.for.gov.bc.ca/hfd/library/FIA/2006/FSP_Y062001b.pdf

Shewell, G.E. (1987). Calliphoridae. In McAlpine, J.F., Peterson, B.V., Shewell, G.E., Teskey, H.J., Vockeroth, J.R. & Wood, D.M. (Eds), Manual of Nearctic Diptera. Volume 2. (pp. 1133 – 1146). Research Branch Agriculture Canada.

Wesolowski, T. (1987). Host–parasite interactions in natural holes: marsh tits (*Parus palustris*) and blow flies (*Protocalliphora falcozi*). *Journal of Zoology, 255*(4), 495-503. doi: https://doi.org/10.1017/S0952836901001571

Cecidomyiidae

Andersson, M.N., Corcoran, J.A., Zhang, D.-D., Hillbur, Y., Newcomb, R.D. & Löfstedt, C. (2016). A sex pheromone receptor in the Hessian fly *Mayetiola destructor* (Diptera, Cecidomyiidae). *Frontiers in cellular neuroscience, 10,* 212. doi: https://doi.org/10.3389/fncel.2016.00212

Gagné, R.J. (1981). Cecidomyiidae. In McAlpine, J.F., Peterson, B.V., Shewell, G.E., Teskey, H.J., Vockeroth, J.R. & Wood, D.M. (Eds), Manual of Nearctic Diptera. Volume 1. (pp. 257 – 292). Research Branch Agriculture Canada.

Heath, J.J. & Stireman, J.O. III (2010). Dissecting the association between a gall midge, *Asteromyia carbonifera*, and its symbiotic fungus, *Botryosphaeria dothidea, 137*(1), 36-49. doi: https://doi.org/10.1111/j.1570-7458.2010.01040.x

Jaschhof, M. & Bae, Y.J. (2019). Twenty new records of mycophagous gall midges (Diptera: Cecidomyiidae) from Korea. *Journal of Species Research, 8*(2), 238-246. doi: https://doi.org/10.12651/JSR.2019.8.2.238

Lo, P.L., Walker, J.T.S. & Suckling, D.M. (2014). Prospects for the control of apple leaf midge *Dasineura mali* (Diptera: Cecidomyiidae) by mass trapping with pheromone lures. *Pest Management Science, 71*(7), 907-913. doi: https://doi.org/10.1002/ps.3857

Sadeghi, R., Odubiyi, S., Nikoukar, A., Schroeder, K.L. & Rashed, A. (2021). *Mayetiola destructor* (Diptera: Cecidmyiidae) host preference and survival on small grains with respect to leaf reflectance and phytohormone concentrations. *Scientific Reports, 11*(1), 4761. doi: https://doi.org/10.1038/s41598-021-84212-x

Schmid, R.B., Knutson, A., Giles, K.L. & McCornack, B.P. (2018). Hessian fly (Diptera: Cecidomyiidae) biology and management in wheat. *Journal of Integrated Pest Management, 9*(1). doi: https://doi.org/10.1093/jipm/pmy008

Scudder, G.G.E. & Cannings, R.A. (2006). The Diptera families of British Columbia. http://www.for.gov.bc.ca/hfd/library/FIA/2006/FSP_Y062001b.pdf

Stireman III, J.O., Devlin, H., Carr, T.G. & Abbot, P. (2010). Evolutionary diversification of the gall midge genus *Asteromyia* (Cecidomyiidae) in a multitrophic ecological context. *Molecular Phylogenetics and Evolution, 54*(1), 194-210. doi: https://doi.org/10.1016/j.ympev.2009.09.010

Stireman III, J.O., Janson, E.M., Carr, T.G., Devlin, H. & Abbot, P. (2008). Evolutionary radiation of *Asteromyia carbonifera* (Diptera: Cecidomyiidae) gall morphotypes on the goldenrod *Solidago altissima* (Asteraceae). *Biological Journal of the Linnean Society, 95*(4), 840-858. doi: https://doi.org/10.1111/j.1095-8312.2008.01101.x

Wearing, C. H., Marshall, R. R., Attfield, B. & Colhoun, C. (2013). Phenology and distribution of the apple leafcurling midge (*Dasineura mali* (Kieffer)) (Diptera: Cecidomyiidae) and its natural enemies on apples under biological and integrated pest management in Central Otago, New Zealand. *New Zealand Entomologist, 36*(2), 87-106. doi: https://doi.org/10.1080/00779962.2012.712887

Ceratopogonidae

Bakhoum, M.T., Fall, A.G., Fall, M., Bassene, C.K., Baldet, T., Seck, M.T., Bouyer, J. Garros, C. & Gimonneau, G. (2016). Insight on the larval habitat of Afrotropical *Culicoides Latreille* (Diptera: Ceratopogonidae) in the Niayes area of Senegal, West Africa. *Parasites & Vectors, 9*(1), 462. doi: https://doi.org/10.1186/s13071-016-1749-1

Bartsch, S., Bauer, B., Wiemann, A., Clausen, P.-H. & Steuber, S. (2009). Feeding patterns of biting midges of the *Culicoides obsoletus* and *Culicoides pulicaris* groups on selected farms in Brandenburg, Germany. *Parasitology Research, 105*(2), 373-380. doi: https://10.1007/s00436-009-1408-y

Büsse, S., Wildermuth, H. & Gorb, S.N. (2022). Morphological adaptations of the mouthparts to the ectoparasitic lifestyle of the biting midge *Forcipomyia paludis* (Diptera: Ceratopogonidae), specialized in Odonata. *Zoomorphology, 141*(3-4), 307-314. doi: https://doi.org/10.1007/s00435-022-00564-6

Díaz, F., Mangudo C., Gleiser, R.M. & Ronderos, M.M. (2019). Redescription of immatures of *Dasyhelea flavifrons* Guérin-Méneville (Culicomorpha: Ceratopogonidae) and new contribution to the knowledge of its larval habitats. *Anais da Academia Brasileira de Ciências, 91*(1). doi: https://doi.org/10.1590/0001-3765201920180047

Dominiak, P. & Borkent, A. (2023). A new species of *Dasyhelea* (Diptera: Ceratopogonidae), mining the leaves of the floating fern *Salvinia minima* Baker. *Journal of Natural History, 57*(9-12). doi: https://doi.org/10.1080/00222933.2023.2203336

Downes, J.A. & Wirth, W.W. (1981). In McAlpine, J.F., Peterson, B.V., Shewell, G.E., Teskey, H.J., Vockeroth, J.R. & Wood, D.M. (Eds), Manual of Nearctic Diptera. Volume 1. (pp. 393 – 422). Research Branch Agriculture Canada.

El-Hawagry, M.S., El-Azab, S.E.-D.A., Abdel-Dayem, M.S. & Al Dhafer, H.M. (2020). Biting midges of Egypt (Diptera: Ceratopogonidae). *Biodiversity Data Journal, 8*(4). doi: https://doi.org/10.3897/BDJ.8.e52357

Erram, D., Black, T.V. & Burkett-Cadena, N. (2021). Host bloodmeal source has no significant effect on the fecundity and subsequent larval development traits of the progeny in *Culicoides furens* Poey (Diptera: Ceratopogonidae). *Journal of Medical Entomology, 58*(6), 2439–2445. doi: https://doi.org/10.1093/jme/tjab085

Erram, D., Blosser, E.M. & Burkett-Cadena, N. (2019). Habitat associations of *Culicoides* species (Diptera: Ceratopogonidae) abundant on a commercial cervid farm in Florida, USA. *Parasites & Vectors, 12*(1), 367. doi: https://doi.org/10.1186/s13071-019-3626-1

Erram, D. & Zurek, L. (2017). Larval development of *Culicoides sonorensis* (Diptera: Ceratopogonidae) in mud supplemented with manure of various farm animals. *Journal of Medical Entomology, 55*(1), 43-50. doi: https://doi.org/10.1093/jme/tjx197

Huerta, H., Romero, D.I. & Díaz, F. (2023). Full redescription and first record of *Dasyhelea bifida* Zilahi-Sebess (Diptera: Ceratopogonidae) from Mexico. *Journal of Natural History, 57*(29-32), 1464-1471. doi: https://doi.org/10.1080/00222933.2023.2257386

Kasičová, Z., Schreiberová, A., Kimáková, A. & Kočišová, A. (2021). Blood meal analysis: host-feeding patterns of biting midges (Diptera, Ceratopogonidae, *Culicoides* Latreille) in Slovakia. *Parasite, 28*, 58. doi: https://doi.org/10.1051/parasite/2021058

Kawahara, A.Y., Winkler, I.S. & Hsu, W.W. (2006). New host records of the ectoparasitic biting midge *Forcipomyia (Trichohelea) pectinunguis* (Diptera: Ceratopogonidae) on adult geometrid moths (Lepidoptera: Geometridae). *Journal of the Kansas Entomological Society, 79*(3), 297-300. doi: https://doi.org/10.2317/0511.15.1

Kitching, R.L. (1972). The immature stages of *Dasyhelea dufouri* Laboulbene (Diptera: Ceratopogonidae) in water-filled tree-holes. *Journal of Entomology Series A, General Entomology, 47*(1), 109-114. doi: https://doi.org/10.1111/j.1365-3032.1972.tb00014.x

Kriska, G. (2013). *Freshwater Invertebrates in Central Europe*. Springer. doi: https://doi.org/10.1007/978-3-7091-1547-3_22

Lane, R.P. & Cotman, H.E. (1985). A new species of *Forcipomyia* (Diptera: Ceratopogonidae) ectoparasitic on butterflies in New Guinea. *Journal of Natural History, 20*(3), 617-620. doi: https://doi.org/10.1080/00222938600770411

Lee, D.J. (1948). Australasian Ceratopogonidae (Diptera, Nematocera). Part I. Relation to disease, biology, general characters and generic classification of the family, with a note on the genus *Ceratopogon*. *Proceedings of the Linnean Society of New South Wales, 72*, 313-331.

Long Jr., W.H. (1902). New species of *Ceratopogon*. *The Biological Bulletin, 3*(1-2), 3-14. doi: https://doi.org/10.2307/1535522

Pfannenstiel, R.S., Mullens, B.A., Ruder, M.G., Zurek, L., Cohnstaedt, L.W. & Nayduch, D. (2015). Management of North American *Culicoides* biting midges: Current knowledge and research needs. *Vector-Borne and Zoonotic Diseases, 15*(6), 374-384. doi: https://doi.org/10.1089/vbz.2014.1705

Purse, B.V., Carpenter, S., Venter, G.J., Bellis, G. & Mullens, B.A. (2015). Bionomics of temperate and tropical *Culicoides* midges: Knowledge gaps and consequences for transmission of *Culicoides*-borne viruses. *Annual Review of Entomology, 60*(1), 373-392. doi: https://doi.org/10.1146/annurev-ento-010814-020614

Riddin, M. A., Venter, G. J., Labuschagne, K. & Villet, M. H. (2019). Bloodmeal analysis in *Culicoides* midges collected near horses, donkeys and zebras in the Eastern Cape, South Africa. *Medical and Veterinary Entomology, 33*(4), 467-475. doi: https://doi.org/10.1111/mve.12381

Salvato, M.H., Salvato, H.L. & Grogan, W.L. (2012). *Forcipomyia (Microhelea) Eriophora* (Williston) (Diptera: Ceratopogonidae) an Ectoparasite of Larval *Anaea troglodyta floridalis* (Nymphalidae). The *Journal of the Lepidopterists' Society, 66*(4), 232-233. doi: https://doi.org/10.18473/lepi.v66i4.a9

Scudder, G.G.E. & Cannings, R.A. (2006). The Diptera families of British Columbia. http://www.for.gov.bc.ca/hfd/library/FIA/2006/FSP_Y062001b.pdf

Steinke, S., Lühken, R., Balczun, C. & Kiel, E. (2016). Emergence of *Culicoides obsoletus* group species from farm-associated habitats in Germany: Emergence of *Culicoides obsoletus* group species. *Medical and Veterinary Entomology, 30*(2), 174-184. doi: https://doi.org/10.1111/mve.12159

Swanson, D.A. & McGregor, B.L. (2022). Life history metrics for *Culex tarsalis* (Diptera: Culicidae) and *Culicoides sonorensis* (Diptera: Ceratopogonidae) are not impacted by artificial feeding on defibrinated versus EDTA-treated blood. *Journal of Medical Entomology, 60*(1), 224–227. doi: https://doi.org/10.1093/jme/tjac171

Urbanek, A. & Rost-Roszkowska, M.M. (2015). Ultrastructural studies on the midgut of biting midge *Forcipomyia nigra* (Winnertz) (Diptera: Ceratopogonidae). *Micron, 69,* 25-34. doi: https://doi.org/10.1016/j.micron.2014.11.003

Van den Eynde, C., Sohier, C., Matthijs, S. & De Regge, N. (2021). Temperature and food sources influence subadult development and blood-feeding response *of Culicoides obsoletus* (sensu lato) under laboratory conditions. *Parasites & Vectors, 14*(1), 1-300. doi: https://doi.org/10.1186/s13071-021-04781-8

Veiga, J., Martínez-de la Puente, J., Václav, R., Figuerola, J. & Valera, F. (2018*). Culicoides paolae* and *C. circumscriptus* as potential vectors of avian haemosporidians in an arid ecosystem. *Parasites & Vectors, 11*(1), 524. doi: https://doi.org/10.1186/s13071-018-3098-8

Werner, D., Groschupp, S., Bauer, C. & Kampen, H. (2020). Breeding habitat preferences of major Culicoides species (Diptera: Ceratopogonidae) in Germany. *International Journal of Environmental Research and Public Health, 17*(14), 5000. doi: https://doi.org/10.3390/ijerph17145000

Zimmer, J.-Y., Brostaux, Y., Haubruge, E. & Francis, F. (2014). Larval development sites of the main *Culicoides* species (Diptera: Ceratopogonidae) in northern Europe and distribution of coprophilic species larvae in Belgian pastures. *Veterinary Parasitology, 205*(3-4), 676-686. doi: https://doi.org/10.1016/j.vetpar.2014.08.029

Chaoboridae

Barth, L.E., Sprules, W.G., Wells, M., Coman,M. & Prairie, Y. (2014). Seasonal changes in the diel vertical migration of *Chaoborus punctipennis* larval instars. *Canadian Journal of Fisheries and Aquatic Sciences, 71*(5), 665-674. doi: https://doi.org/10.1139/cjfas-2013-0440

Härkönen, L., Pekcan-Hekim, Z., Hellén, N. & Horppila, J. (2013). Feeding efficiency of *Chaoborus flavicans* (Insecta, Diptera) under turbulent conditions. *Hydrobiologia, 722*(1), 9-17. doi: https://doi.org/10.1007/s10750-013-1670-y

Jäger, I.S., Hölker, F., Flöder, S. & Walz, N. (2011). Impact of *Chaoborus flavicans* -predation on the zooplankton in a mesotrophic lake - a three year study. *International Review of Hydrobiology, 96*(1), 191-208. doi: https://doi.org/10.1002/iroh.201011253

Moore, M.V. (1988). Differential use of food resources by the instars of *Chaoborus punctipennis*. *Freshwater Biology, 19*(2), 249-268. doi: https://doi.org/10.1111/j.1365-2427.1988.tb00346.x

Salmela, J., Härmä, O. & Taylor, D.J. (2021). *Chaoborus flavicans* Meigen (Diptera, Chaoboridae) is a complex of lake and pond dwelling species: a revision. *Zootaxa, 4927*(2), 151-196. doi: https://doi.org/10.11646/zootaxa.4927.2.1

Schröder, A. & Dam, H.G. (2013). Density- and size-dependent winter mortality and growth of late *Chaoborus flavicans* larvae. *PLoS One, 8*(10), p.e75839. doi: https://doi.org/10.1371/journal.pone.0075839

Chironomidae

Anderson, A.M., Stur, E. & Ekrem, T. (2013). Molecular and morphological methods reveal cryptic diversity and three new species of Nearctic *Micropsectra* (Diptera:Chironomidae). *Freshwater Science, 32*(3), 892-921. doi: https://doi.org/10.1899/12-026.1

Anderson, J.F. & Hitchcock, S.W. (1968). Biology of *Chironomus atrella* in a Tidal Cove. *Annals of the Entomological Society of America, 61*(6), 1597–1603. https://doi.org/10.1093/aesa/61.6.1597

Baniszewski, J., Miller, N., Kariuki, E.M., Cuda, J.P. & Weeks, E.N.I. (2020). *Cricotopus lebetis* intraspecific competition and damage to *Hydrilla*. *The Florida Entomologist, 103*(1), 32-37. doi: https://doi.org/10.1653/024.103.0405

Belle, S., Millet, L., Gillet, F., Verneaux, V. & Magny, M. (2015). Assemblages and paleo-diet variability of subfossil Chironomidae (Diptera) from a deep lake (Lake Grand Maclu, France). *Hydrobiologia, 755*(1), 145-160. doi: https://doi.org/10.1007/s10750-015-2222-4

Boggero, A., Füreder, L., Lencioni, V., Simcic, T., Thaler, B., Ferrarese, U., Lotter, A. F. & Ettinger, R. (2006). Littoral chironomid communities of Alpine lakes in relation to environmental factors. *Hydrobiologia, 562*(1), 145-165. doi: https://doi.org/10.1007/s10750-005-1809-6

Boonsoong, B. (2016). Phoretic associations between *Nanocladius asiaticus* (Diptera, Chironomidae) and its hosts *Gestroiella* (Heteroptera, Naucoridae) and *Euphaea masoni* (Odonata, Euphaeidae) in streams in Western Thailand. *Annales de Limnologie, 52*, 163-169. doi: https://doi.org/10.1051/limn/2015025

Brown, N.E., Mitchell, S.C. & Garbary, D.J. (2013). Host selection by larvae of a marine insect *Halocladius variabilis*: nutritional dependency or escape from predation? *Journal of the Marine Biological Association of the United Kingdom, 93*(5), 1373-1379. doi: https://doi.org/10.1017/S0025315412001634

Cranston, P.S. (1984). The taxonomy and ecology of *Orthocladius* (Eudactylocladius) *fuscimanus* (Kieffer), a hygropetric chironomid (Diptera). *Journal of Natural History, 18*(6), 873-895. doi: https://doi.org/10.1080/00222938400770771

Cranston, P.S. & Saether, O.A. (1984). *Rheosmittia* (Diptera: Chironomidae): a generic validation and revision of the western Palaearctic species. *Journal of Natural History, 20*(1), 31-51. doi: https://doi.org/10.1080/00222938600770041

Cuda, J.P., Coon, B.R., Dao, Y.M. & Center, T.D. (2002). Biology and Laboratory Rearing of *Cricotopus lebetis* (Diptera: Chironomidae), a Natural Enemy of the Aquatic Weed *Hydrilla* (Hydrocharitaceae). *Annals of the Entomological Society of America, 95*(5), 587–596. doi: https://doi.org/10.1603/0013-8746(2002)095[0587:BALROC]2.0.CO;2

Delettre, Y.R. (2005). Short-range spatial patterning of terrestrial Chironomidae (Insecta: Diptera) and farmland heterogeneity. *Pedobiologia, 49*(1), 15-27. doi: https://doi.org/10.1016/j.pedobi.2004.06.010

Dosdall, L.M. & Parker, D.W. (1998). First report of a symphoretic association between *Nanocladius branchicolus* Saether (Diptera: Chironomidae) and *Argia moesta* (Hagen) (Odonata: Coenagrionidae). *The American Midland Naturalist, 139*(1), 181-185. Doi: https://doi.org/10.1674/0003-0031(1998)139[0181:FROASA]2.0.CO2

Doucett, R.R., Giberson, D.J. & Power, G. (1999). Parasitic association of *Nanocladius* (Diptera: Chironomidae) and *Pteronarcys biloba* (Plecoptera: Pteronarcyidae): insights from stable-isotope analysis. *Journal of the North American Benthological Society, 18*(4), 514-523. doi: https://doi.org/10.2307/1468383

Earle, W., Mangan, R., O'Brien, M. & Baars, J. (2013). Biology of *Polypedilum* n. sp. (Diptera: Chironomidae), a promising candidate agent for the biological control of the aquatic weed *Lagarosiphon* major (Hydrocharitaceae) in Ireland. *Biocontrol Science and Technology, 23*(11), 1267-1283. doi: https://doi.org/10.1080/09583157.2013.826344

Egan, A.T. & Ferrington, L.C. (2015). Chironomidae (Diptera) in freshwater coastal rock pools at Isle Royale, Michigan. *Transactions of the American Entomological Society, 141*(1), 1-25. doi: https://doi.org/10.3157/061.141.0102

Einarsson, Á., Gardarsson, A., Gíslason, G.M. & Ives, A.R. (2002). Consumer–resource interactions and cyclic population dynamics of *Tanytarsus gracilentus* (Diptera: Chironomidae). *Journal of Animal Ecology, 71*(5), 832-845. doi: https://doi.org/10.1046/j.1365-2656.2002.00648.x

Ekrem, T. (2007). A taxonomic revision of the genus *Stempellinella* (Diptera: Chironomidae). *Journal of Natural History, 41*(21-24), 1367-1465. doi: https://doi.org/10.1080/00222930701437360

Enushchenko, I.V. (2018). Most interesting findings from remnants of Chironomid (Insecta: Diptera: Chironomidae) larvae in layers of a 200-year-old stratum of lake Oron bottom sediments. *Russian Journal of Ecology, 49*(1), 69-74. doi: https://doi.org/10.1134/S1067413618010058
Giłka, W. (2011). Six unusual *Cladotanytarsus* Kieffer: towards a systematics of the genus and resurrection of *Lenziella* Kieffer (Diptera: Chironomidae: Tanytarsini). *Zootaxa, 3100*(1), 1-34. doi: https://doi.org/10.11646/zootaxa.3100.1.1

Goffová, K., Bitušik, P., Čiamporová-Zaťovičová, Z., Bukvová, D. & Hamerlí, L. (2015). Seasonal dynamics and life cycle of *Heterotrissocladius marcidus* (Diptera: Chironomidae) in high altitude lakes (High Tatra Mts, Slovakia). *Biologia, 70*(7), 943-947. doi: https://doi.org/10.1515/biolog-2015-0103

Hamerlik, L. & da Silva, F.L. (2020). *Chironomidae of Central America*. CRC Press. https://doi.org/10.1201/9780429021572

Hänel, C. & Chown, S.L. (1997). The impact of a small, alien invertebrate on a sub-Antarctic terrestrial ecosystem: *Limnophyes minimus* (Diptera, Chironomidae) at Marion Island. *Polar Biology, 20*(2), 99-106. doi: https://doi.org/10.1007/s003000050282

Hayford, B. (2012). Parochlus Kiefferi (Garrett, 1925) in Nebraska (Diptera: Chrionomidae). *Great Plains Research, 22*(1), 27-33.

Hazra, N., Saha, G.K. & Chaudhuri, P.K. (2002). Records of Orthoclad species from the Darjeeling-Sikkim Himalayas of India (Diptera: Chironomidae), with notes on their ecology. *Hydrobiologia, 474*(1), 41-55. doi: https://doi.org/10.1023/A:1016511702944

Hazra, N., Som, D.K. and Chaudhuri, P.K. (1998). Immatures of *Rheocricotopus* (*Psilocricotopus*) valgus Chaudhuri & Sinharay of Darjeeling Himalaya with notes on ecology (Diptera : Chironomidae). *International Journal of Limnology, 34*(1), 75-82. doi: https://doi.org/10.1051/limn/1998008

Hershey, A.E. (1986). Selective predation by *Procladius* in an Arctic Alaskan lake. *Canadian Journal of Fisheries and Aquatic Sciences, 43*(12), 2523-2528. doi: https://doi.org/10.1139/f86-312

Ingvason, H.R., Ólafsson, J.S. & Gardarsson, A. (2004). Food selection of *Tanytarsus gracilentus* larvae (Diptera: Chironomidae): An analysis of instars and cohorts. *Aquatic Ecology, 38*(2), 231-237. doi: https://doi.org/10.1023/B:AECO.0000032053.67992.03

Inoue, E., Kawai, K. & Imabayashi, H. (2004). A new species of the genus *Stempellinella* (Diptera: Chironomidae) from Hiroshima, Japan. *Limnology, 5*(3), 141-147. doi: https://doi.org/10.1007/s10201-004-0131-8

Jackson, G.A. (1977). Nearctic and Palaearctic *Paracladopelma* Harnisch and *Saetheria* n.gen. (Diptera: Chironomidae). *Journal of the Fisheries Board of Canada, 34*(9), 1321-1359. doi: https://doi.org/10.1139/f77-194

Johnson, R.K. (1987). Seasonal variation in diet of *Chironomus plumosus* (L.) and *C. anthracinus* Zett. (Diptera: Chironomidae) in mesotrophic Lake Erken. *Freshwater Biology, 17*(3), 525-532. doi: https://doi.org/10.1111/j.1365-2427.1987.tb01073.x

Langton, P.H. (1994). A redescription of *Parakiefferiella* sp. D. Wulker, the pupa of *Parakiefferiella wuelkeri* Moubayed (Diptera: Chironomidae), a species new to Britain. *British Journal of Entomology and Natural History, 7*, 11-13.

Langton, P.H. & Cobo, F. (1992). *Hydrobaenus cranstoni* n. sp. (Diptera: Chironomidae) from north-west Spain. *British Journal of Entomology and Natural History, 5,* 139-141.

Li, Z. & Tang, H. (2021). Two new species of *Paratanytarsus* Thienemann & Bause (Diptera: Chironomidae) from Oriental China. *Zootaxa, 4903*(3), 430-438. doi: https://doi.org/10.11646/zootaxa.4903.3.8

Lilley, T.M., Ruokolainen, L., Pikkarainen, A., Laine, V.N., Kilpimaa, J., Rantala, M.J., Nikinmaa, M. (2012). Impact of Tributyltin on immune response and life history traits of *Chironomus riparius*: single and multigeneration effects and recovery from pollution. *Environmental Science & Technology, 46*(13), 7382-7389. doi: https://doi.org/10.1021/es300536t

Kariuki, E.M., Cuda, J.P., Hight, S.D., Hix, R.L., Gettys, L.A. & Gillett-Kaufman, J.L. (2019). Foraging depth of *Cricotopus lebetis* larvae. *Journal of Aquatic Plant Management, 57*(2), 69-78.

Kesler, D.H. (1981). Grazing rate determination of *Corynoneura scutellata* Winnertz (Chironomidae: Diptera). *Hydrobiologia, 80*(1), 63-66. doi: https://doi.org/10.1007/BF00130681

Kriska, G. (2013). *Freshwater Invertebrates in Central Europe*. Springer. doi: https://doi.org/10.1007/978-3-7091-1547-3_22

Martin, J., Andreeva, E.N., Kiknadze, I.I. & Wülker, W.F. (2007). Polytene chromosomes and phylogenetic relationships of Chironomus atrella (Diptera: Chironomidae) in North America. *Genome, 49*(11), 1384-1392. doi: https://doi.org/10.1139/g06-095

Medeiros, A.S. & Quinlan, R. (2011). The distribution of the Chironomidae (Insecta: Diptera) along multiple environmental gradients in lakes and ponds of the eastern Canadian Arctic. *Canadian Journal of Fisheries and Aquatic Sciences, 68*(9), 1511-1527. doi: https://doi.org/10.1139/F2011-076

Moore, J.W. (1979). Factors influencing algal consumption and in *Heterotrissocladius changi* Saether and *Polypedilum nebeculosum* (Meigen) (Chironomidae: Diptera). *Oecologia, 40*(2), 219-227. doi: https://doi.org/10.1007/BF00347939

Moubayed, J. (1994). *Parakiefferiella wuelkeri* n. sp. (Diptera: Chironomidae) from western Europe and North Africa. *British Journal of Entomology and Natural History, 7*, 7-10.

Moubayed, J. & Langton, P.H. (1996). *Krenopsectra nohedensis* n. sp. and the pupal exuviae of *Micropsectra auvergnensis* Reiss (Diptera: Chironomidae) from the eastern Pyrenee. *British Journal of Entomology and Natural History, 9*, 77-86.

Moubayed-Breil, J. & Bitušík, P. (2019). Taxonomic notes on the genus *Chaetocladius* (laminatus-group) (Diptera: Chironomidae, Orthocladiinae) II. Descriptions of *C. bitusiki* sp. n. and *C. mantetensis* sp. n., two relict species inhabiting cold alpine springs and streams. *Biologia, 74*(11), 1489-1500. doi: https://doi.org/10.2478/s11756-019-00253-8

Namayandeh, A. & Beresford, D.V. (2019). Chironomidae of Fosheim Peninsula, Ellesmere Island: first female description of *Chaetocladius* (Chaetocladius) *glacialis* (Lundström, 1915) (Diptera: Chironomidae: Orthocladiinae) and new faunistic records from the Nunavut (Canada) and the Nearctic Region. *Aquatic Insects, 40*(1), 1-18. doi: https://doi.org/10.1080/01650424.2018.1523435

Oliver, D.R. (1981). Chironomidae. In McAlpine, J.F., Peterson, B.V., Shewell, G.E., Teskey, H.J., Vockeroth, J.R. & Wood, D.M. (Eds), Manual of Nearctic Diptera. Volume 1. (pp. 423 – 458). Research Branch Agriculture Canada.

Paggi, A.C. (1993). Redescription of *Pseudosmittia bilobulata* (Edw.) comb. n ( = Spaniotoma (*Smittia*) bilobulata Edwards 1931) and description of *P. neobilobulata* sp.n. (Diptera : Chironomidae) from Argentina. *International Journal of Limnology, 29*(2), 171-174. doi: https://doi.org/10.1051/limn/1993015

Pett, L.A., Linde, S. & Gotelli, N.J. (2022). Midge larvae *Metriocnemus knabi* can emigrate to new pitchers within *Sarracenia purpurea* after pitcher drainage. *Northeastern Naturalist, 29*(3), 335-341. doi: https://doi.org/10.1656/045.029.0304

Puchalski, M., Zimny, F. & Giłka, W. (2016). *Cladotanytarsus molestus* Hirvenoja, 1962 in Poland: toward the identification of bioindicative Tanytarsini (Diptera: Chironomidae). *Oceanological and Hydrobiological Studies, 45*(3), 316-323. doi: https://doi.org/10.1515/ohs-2016-0030

Rempel, J.G. (1936). The Life-History and Morphology of *Chironomus hyperboreus*. *Journal of the Biological Board of Canada*. doi: https://doi.org/10.1139/f36-007

Ricciardi, A. (1994). Occurrence of chironomid larvae (*Paratanytarsus* sp.) as commensals of dreissenid mussels (*Dreissena polymorpha* and *D. bugensis*). *Canadian Journal of Zoology, 72*(6), 1159-1162. doi: https://doi.org/10.1139/z94-155

Roback, S.S. (1986). The Immature Chironomids of the Eastern United States VIII. Pentaneurini: Genus *Nilotanypus*, with the Description of a New Species from Kansas. *Proceedings of the Academy of Natural Sciences of Philadelphia, 138*(2), 443-465.

Schütz, S.A., Brittain, J.E. & Füreder, L. (2022). Diverging life cycle patterns of two *Diamesa* species (Diptera, Chironomidae) in High Arctic streams, Svalbard. *Polar Biology, 45*(2), 285-296. doi: https://doi.org/10.1007/s00300-021-02987-1

Scudder, G.G.E. & Cannings, R.A. (2006). The Diptera families of British Columbia. http://www.for.gov.bc.ca/hfd/library/FIA/2006/FSP_Y062001b.pdf

Silva, F.L.D., Dantas, G.P.S. & Hamada, N. (2019). Description of immature stages of *Ablabesmyia cordeiroi* Neubern, 2013 (Diptera: Chironomidae: Tanypodinae). *Acta Amazonica, 49*(2), 118-121. doi: https://doi.org/10.1590/1809-4392201801151

Silva, C.J.M., Silva, A.L.P., Gravato, C. & Pestana, J.L.T. (2019). Ingestion of small-sized and irregularly shaped polyethylene microplastics affect *Chironomus riparius* life-history traits. *The Science of the Total Environment, 672*, 862-868. doi: https://doi.org/10.1016/j.scitotenv.2019.04.017

Skuse, F.A.A. (1889). Diptera of Australia. Part VI. The Chironomidae. *Proceedings of the Linnean Society of New South Wales, 4*, 215-311.

Som, D.K., Das, N. & Hazra, N. (2013). Systematics and biology of Metriocnemus clarivirgulus sp. n. (Diptera: Chrionomidae) from Darjeeling, India with revised keys to male and female adults of Metriocnemus van der Wulp. *Deutsche Entomologische Zeitschrift*, *60*(1), 111-121. doi: https://doi.org/10.1002/mmnd.201300014

Syrovátka, V. (2018). The predatory behaviour of *Monopelopia tenuicalcar* (Kieffer, 1918) larvae in a laboratory experiment. *Journal of Limnology, 77.* doi: https://doi.org/10.4081/jlimnol.2018.1792

Tarakhovskaya, E.R. & Garbary, D.J. (2009). *Halocladius variabilis* (Diptera: Chironomidae): a marine insect symbiotic with seaweeds from the White Sea, Russia. *Journal of the Marine Biological Association of the United Kingdom, 89*(7), 1381-1385. doi: https://doi.org/10.1017/S0025315409000071

Tarkowska-Kukuryk, M. (2014). Spatial distribution of epiphytic chironomid larvae in a shallow macrophyte-dominated lake: effect of macrophyte species and food resources. *Limnology, 15*(2), 141-153. doi: https://doi.org/10.1007/s10201-014-0425-4

Vodopich, D.S. & Cowell, B.C. (1984). Interaction of Factors Governing the Distribution of a Predatory Aquatic Insect. *Ecology, 65*(1), 39-52. doi: https://doi.org/10.2307/1939456

Wiedenbrug, S. & da Silva, F.L. (2013). New species of *Nanocladius* Kieffer, 1913 (Diptera: Chironomidae: Orthocladiinae) from Neotropical region. *International Journal of Limnology, 49*(4), 255-264. doi: http://dx.doi.org/10.1051/limn/2013057

Winnell, M.H. & White, D.S. (1986). The distribution of *Heterotrissocludius oliveri* Saether (Diptera: Chironomidae) in Lake Michigan. *Hydrobiologia, 131*(3), 205-214. doi: https://doi.org/10.1007/BF00008856

Young, R.M. (1969). Field observations on a midwinter breeding flight of *Diamesa arctica* (Diptera: Chironomidae). *Annals of the Entomological Society of America, 62*(5), 1204. doi: https://doi.org/10.1093/aesa/62.5.1204

Chloropidae

Allen, W.A. & Pienkowski, R.L. (1974). The biology and seasonal abundance of the fruit fly, *Oscinella* in reed canarygrass in Virginia. *Annals of the Entomological Society of America, 67*(4), 539-544. doi: https://doi.org/10.1093/aesa/67.4.539

Ellis, W.N. (2020). *Parasites*. Plant Parasites of Europe. https://bladmineerders.nl/parasites/

Foster, G.A. (2022). Notes on the Genus *Apallates* Sabrosky (Diptera: Chloropidae). *Proceedings of the Entomological Society of Washington, 124*(1), 46-56. doi: https://doi.org/10.4289/0013-8797.124.1.46

Klepzig, K.D., Hartshorn, J.A., Tsalickis, A. & Sheehan, T.N. (2022). Eye gnat (*Liohippelates*, Diptera: Chloropidae) biology, ecology, and management: Past, present, and future. *Journal of Integrated Pest Management, 13*(1), 19. doi: https://doi.org/10.1093/jipm/pmac015

Mlynarek, J.J. & Wheeler, T.A. (2018). Chloropid flies (Diptera, Chloropidae) associated with pitcher plants in North America. *PeerJ, 6*. doi: https://doi.org/10.7717/peerj.4491

Nartshuk, E.P. (2014). Grass-fly larvae (Diptera, Chloropidae): Diversity, habitats, and feeding specializations. *Entomological Review, 94*(4), 514-525. doi: https://doi.org/10.1134/S001387381404006X

Nartshuk, E. & Hugo Andersson, H. (2013). The Frit Flies (Chloropidae, Diptera) of Fennoscandia and Denmark. *Fauna Entomologica Scandinavica, 43.* doi: http://doi.org/10.1163/9789004190665

Nartshuk, E.P. & Khruleva, O.A. (2011). Plant-feeding dipterans (Diptera, Chloropidae, Agromyzidae) from Wrangel Island (the Chukchi Sea). *Entomological Review, 91*(7), 849-854. doi: https://doi.org/10.1134/S0013873811070062

Nielsen, L.B. & Nielsen, B.O. (1984). *Oscinella frit* (L.) and *O. pusilla* (Mg.) (Diptera, Chloropidae) in agricultural grass in Denmark. *Zeitschrift für Angewandte Entomologie, 98*(1-5), 264-275. doi: https://doi.org/10.1111/j.1439-0418.1984.tb02711.x

Riccardi, P. (2016). Family Chloropidae. *Zootaxa, 4122*(1), 696-707. doi: http://dx.doi.org/10.11646/zootaxa.4122.1.60

Rogers, T.P., Foote, B.A. and Todd, J.L. (1991). Biology and immature stages of *Chlorops certimus* and *Epichlorops exilis* (Diptera: Chloropidae), stem-borers of wetland sedges. *Journal of the New York Entomological Society, 99*(4), 664-683. doi: https://www.jstor.org/stable/25009932

Sabrosky, C.W. (1987). Chloropidae. In McAlpine, J.F., Peterson, B.V., Shewell, G.E., Teskey, H.J., Vockeroth, J.R. & Wood, D.M. (Eds), Manual of Nearctic Diptera. Volume 2. (pp. 1049 – 1068). Research Branch Agriculture Canada.

Scudder, G.G.E. & Cannings, R.A. (2006). The Diptera families of British Columbia. http://www.for.gov.bc.ca/hfd/library/FIA/2006/FSP_Y062001b.pdf

Vickerman, G.P. (1978). Host plant preferences of *Oscinella* spp. (Diptera: Chloropidae) in the laboratory. *Annals of Applied Biology, 89*(3), 379-386. doi: https://doi.org/10.1111/j.1744-7348.1978.tb05963.x

Yarkulov, F.Y. (2019). Biology of *Thaumatomyia* Zenker, 1833 (Diptera, Chloropidae) frit flies, predators of root aphids in Middle Asia. *Entomological review, 99*(8), 1069-1082. doi: https://doi.org/0.1134/S0013873819080013

Clusiidae

Roháček, J. (2012). The fauna of the opomyzoid families Clusiidae, Acartophthalmidae, Anthomyzidae, Opomyzidae, Stenomicridae, Periscelididae, Asteiidae (Diptera) in the Gemer area (Central Slovakia). Časopis Slezského Zemského Muzea, 61(2), 97-111. doi: https://doi.org/10.2478/v10210-012-0011-5

Roháček, J. (2013). The fauna of the Acalyptrate families Micropezidae, Psilidae, Clusiidae, Acartophthalmidae, Anthomyzidae, Aulacigastridae, Periscelididae and Asteiidae (Diptera) in the Gemer area (Central Slovakia): supplement 1. *Acta Musei Silesiae: Scientiae Naturales, 62*(2), 125. doi: https://doi.org/10.2478/cszma-2013-0014

Roháček, J., Andrade, R., Gonçalves, A.R. & Almeida, J.M. (2016). New records of Micropezidae, Clusiidae and Periscelididae (Diptera: Acalyptrata) from Portugal. *Acta Musei Silesiae, Scientiae Naturales, 65*(2), 153-166. doi: https://doi.org/10.1515/cszma-2016-0020

Scudder, G.G.E. & Cannings, R.A. (2006). The Diptera families of British Columbia. http://www.for.gov.bc.ca/hfd/library/FIA/2006/FSP_Y062001b.pdf

Soós, À. (1987). Clusiidae. In McAlpine, J.F., Peterson, B.V., Shewell, G.E., Teskey, H.J., Vockeroth, J.R. & Wood, D.M. (Eds), Manual of Nearctic Diptera. Volume 2. (pp. 853 – 858). Research Branch Agriculture Canada.

Coelopidae

Edward, D.A., Blyth, J.E., Mckee, R. & Gilburn, A.S. (2007). Change in the distribution of a member of the strand line community: the seaweed fly (Diptera: Coelopidae). *Ecological Entomology, 32*(6), 741-746. doi: https://doi.org/10.1111/j.1365-2311.2007.00919.x

Edward, D.A., Newton, J. & Gilburn, A.S. (2008). Investigating dietary preferences in two competing dipterans, *Coelopa frigida* and *Coelopa pilipes*, using stable isotope ratios of carbon and nitrogen. *Entomologia Experimentalis et Applicata, 127*(3), 169-175. doi: https://doi.org/10.1111/j.1570-7458.2008.00692.x

Rotheray, G. & Lyszkowski, R. (2015). Diverse mechanisms of feeding and movement in Cyclorrhaphan larvae (Diptera). *Journal of Natural History, 49*(35-36), 2139-2211. doi: https://doi.org/10.1080/00222933.2015.1010314

Scudder, G.G.E. & Cannings, R.A. (2006). The Diptera families of British Columbia. http://www.for.gov.bc.ca/hfd/library/FIA/2006/FSP_Y062001b.pdf

Vockeroth, J.R. (1987). Coelopidae. In McAlpine, J.F., Peterson, B.V., Shewell, G.E., Teskey, H.J., Vockeroth, J.R. & Wood, D.M. (Eds), Manual of Nearctic Diptera. Volume 2. (pp. 919 – 922). Research Branch Agriculture Canada.

Culicidae

Ahebwa, A., Hii, J., Neoh, K-B. & Chareonviriyaphap, T. (2023). *Aedes aegypti* and *Aedes albopictus* (Diptera: Culicidae) ecology, biology, behaviour, and implications on arbovirus transmission in Thailand: Review. *One Health, 16*. doi: https://doi.org/10.1016/j.onehlt.2023.100555

Al-Jaran, T.K.H. & Katbeh-Bader, A.M. (2010). Laboratory Studies on the Biology of Culiseta longiareolata(Macquart) (Diptera: Culicidae). *International Journal of Freshwater Entomology, 23*(1), 11-22. doi: https://doi.org/10.1076/aqin.23.1.11.4928

Anderson, I.H. (1992). The effect of sugar meals and body size on fecundity and longevity of female *Aedes communis* (Diptera: Culicidae). *Physiological Entomology, 17*(3), 203-207. doi: https://doi.org/10.1111/j.1365-3032.1992.tb01011.x

Bartlett, K. (2009). *Factors contributing to the host specificity of the frog-feeding mosquito Culex territans Walker (Diptera: Culicidae).* [Doctoral dissertation, Rutgers University]. RUcore: Rutgers University Community Repository.

Friend, W.G., Smith, J.J.B., Schmidt, J.M. & Tanner, R.J. (1989). Ingestion and diet destination in *Culiseta inornata*: responses to water, sucrose and cellobiose. *Physiological Entomology, 14*(2), 137-146. doi: https://doi.org/10.1111/j.1365-3032.1989.tb00945.x

Frohne, W.C. (1953). Natural history of *Culiseta impatiens* (Wlk.), (Diptera, Culicidae), in Alaska. *Transactions of the American Microscopical Society, 72*(2), 103-118. doi: https://www.jstor.org/stable/3223507

Gadawski, R.M. & Smith, S.M. (1992). Nectar sources and age structure in a population of *Aedes provocans* (Ditpera: Culicidae). *Journal of Medical Entomology, 29*(5), 879-886. doi: https://doi.org/10.1093/jmedent/29.5.879

Garrett, M. & Bradley, T.J. (1984). The pattern of osmotic regulation in larvae of the mosquito *Culiseta Inornata*. *Journal of Experimental Biology, 113*(1), 133-141. doi: https://doi.org/10.1242/jeb.113.1.133

Greenberg, J.A., Lujan, D.A., DiMenna, M.A., Wearing, H.J. & Hofkin, B.V. (2013). Identification of blood meal sources in *Aedes vexans* and *Culex quinquefasciatus* in Bernalillo County, New Mexico. *Journal of Insect Science, 13*(75), 1-12. doi: https://doi.org/10.1673/031.013.7501

Guarido, M.M., Riddin, M.A., Johnson, T., Braack, L.E.O., Schrama, M., Gorsich, E.E., Brooke, B.D., Almeida, A.P.G. & Venter, M. (2021). *Aedes* species (Diptera: Culicidae) ecological and host feeding patterns in the north-eastern parts of South Africa, 2014–2018. *Parasites & Vectors, 14*. doi: https://doi.org/10.1186/s13071-021-04845-9

Haufe, W.O. (2012). Physical environment and behaviour of immature stages of *Aedes communis* (Deg.) (Diptera: Culicidae) in Subarctic Canada. *The Canadian Entomologist, 89*(3), 120 – 139. doi: https://doi.org/10.4039/Ent89120-3

Jensen, T. & Washino, R.K. (1991). An assessment of the biological capacity of a Sacramento Valley population of *Aedes Melanimon* to vector arboviruses. *The American Journal of Tropical Medicine and Hygiene, 44*(4), 355-363. doi: https://doi.org/10.4269/ajtmh.1991.44.355

Kaufman, M.G. & and Fonseca, D.M. (2014). Invasion Biology of *Aedes japonicus japonicus* (Diptera: Culicidae). *Annual Review of Entomology, 59*(1), 31-49. doi: https://doi.org/10.1146/annurev-ento-011613-162012

Kriska, G. (2013). *Freshwater Invertebrates in Central Europe*. Springer. doi: https://doi.org/10.1007/978-3-7091-1547-3_22

Lindström, A., Eklöf, D. & Lilja, T. (2021). Different hatching rates of floodwater mosquitoes *Aedes sticticus, Aedes rossicus* and *Aedes cinereus* from different flooded environments. *Insects, 12*(4), 279. doi: https://doi.org/10.3390/insects12040279

Organization for Economic Co-operation and Development. (2018). Safety Assessment of Transgenic Organisms in the Environment, Volume 8 : OECD Consensus Document of the Biology of Mosquito *Aedes aegypti.* Organization for Economic Co-operation and Development.

Rogers, R.E. & Yee, D.A. (2019). Response of *Aedes aegypti* and *Aedes albopictus* (Diptera: Culicidae) survival, life history, and population growth to oak leaf and acorn detritus. *Journal of Medical Entomology, 56*(2), 303–310. doi: https://doi.org/10.1093/jme/tjy172

Schäfer, M.L. & Lundstroem, J.O. (2006). Different responses of two floodwater mosquito species, *Aedes vexans* and *Ochlerotatus sticticus* (Diptera: Culicidae), to larval habitat drying. *Journal of Vector Ecology, 31*(1), 123-128.

Scudder, G.G.E. & Cannings, R.A. (2006). The Diptera families of British Columbia. http://www.for.gov.bc.ca/hfd/library/FIA/2006/FSP_Y062001b.pdf

Sherwood, J. A., Stehman, S., Howard, J. J. & Oliver, J. (2020). Cases of Eastern equine encephalitis in humans associated with *Aedes canadensis*, *Coquillettidia perturbans* and *Culiseta melanura* mosquitoes with the virus in New York State from 1971 to 2012 by analysis of aggregated published data. *Epidemiology and Infection, 148*, p.e72. doi: https://doi.org/10.1017/S0950268820000308

Slaff, M. & Haefner, J.D. (1985). Seasonal and spatial distribution of *Mansonia Dyari*, *Mansonia Titillans,* and *Coquillettidia Perturbans* (Diptera: Culicidae) in the Central Florida, USA, Phosphate Region. *Journal of Medical Entomology, 22*(6), 624–629. doi: https://doi.org/10.1093/jmedent/22.6.624

Smith, S.M. & Gadawski, R.M. (1994). Nectar feeding by the early-spring mosquito *Aedes provocans*. *Medical and Veterinary Entomology, 8*(3), 201-213. doi: https://doi.org/10.1111/j.1365-2915.1994.tb00499.x

Stone, A. (1981). Culicidae. In McAlpine, J.F., Peterson, B.V., Shewell, G.E., Teskey, H.J., Vockeroth, J.R. & Wood, D.M. (Eds), Manual of Nearctic Diptera. Volume 1. (pp. 341 – 350). Research Branch Agriculture Canada.

Turell, M.J., Lundström, J.O. & Niklasson, B. (1990). Transmission of Ockelbo virus by *Aedes cinereus*, *Ae. communis*, and *Ae. Excrucians* (Diptera: Culicidae) collected in an enzootic area in Central Sweden. *Journal of Medical Entomology, 27*(3), 266–268. doi: https://doi.org/10.1093/jmedent/27.3.266

Yee, W.L., Foster, W.A., Howe, M.J. & Hancock, R.G. (1992). Simultaneous field comparison of evening temporal distributions of nectar and blood feeding by *Aedes vexans and Ae. trivittatus* (Diptera: Culicidae) in Ohio. *Journal of Medical Entomology, 29*(2), 356-360. doi: https://doi.org/10.1093/jmedent/29.2.356

Dolichopodidae

Aukema, B.H. & Raffa, K.F. (2004). Behavior of adult and larval *Platysoma cylindrica* (Coleoptera: Histeridae) and larval *Medetera bistriata* (Diptera: Dolichopodidae) during subcortical predation of *Ips pini* (Coleoptera: Scolytidae). *Journal of Insect Behavior, 17*, 115–128. https://doi.org/10.1023/B:JOIR.0000025136.73623.f5

Gelbič, I. & Olejníček, J. (2011). Ecology of Dolichopodidae (Diptera) in a wetland habitat and their potential role as bioindicators. *Central European Journal of Biology, 6*(1), 118-129. doi: https://doi.org/10.2478/s11535-010-0098-x

Grichanov, I.Y. (2020). New species of *Hercostomus* Loew, 1857 from Afrotropics (Diptera: Dolichopodidae) and key to Afrotropical fauna. *European Journal of Taxonomy, 722*. doi: https://doi.org/10.5852/ejt.2020.722.1131

Kechev, M., Belilov, S. & Georgiev, G. (2022). Two predatory *Medetera* (Diptera: Dolichopodidae) species associated with *Ips acuminatus* (Gyllenhal) (Coleoptera: Curculionidae) in Bulgaria. *Historia Naturalis Bulgarica, 44*(8), 63-67. doi: https://doi.org/10.48027/hnb.44.081

Pollet, M. (1993). Morphological and ecological characterization of *Hercostomus* (*Hercostomus*) *plagiatus* and a sibling species, *H. verbekei* sp.n. (Diptera: Dolichopodidae). *Zoologica Scripta, 22*(1), 101-109. doi: https://doi.org /10.1111/j.1463-6409.1993.tb00344.x

Pollet, M.A.A., Brooks, S.E. & Cumming, J.M. (2004). Catalog of the Dolichopodidae (Diptera) of America north of Mexico. *Bulletin of the American Museum of Natural History, 2004*(283), 1-114. doi: https://doi.org/10.1206/0003-0090(2004)283%3C0001:COTDDO%3E2.0.CO;2

Scudder, G.G.E. & Cannings, R.A. (2006). The Diptera families of British Columbia. http://www.for.gov.bc.ca/hfd/library/FIA/2006/FSP_Y062001b.pdf

Vockeroth, J.R. & Robinson, H. (1981). Dolichopodidae. In McAlpine, J.F., Peterson, B.V., Shewell, G.E., Teskey, H.J., Vockeroth, J.R. & Wood, D.M. (Eds), Manual of Nearctic Diptera. Volume 1. (pp. 625 – 640). Research Branch Agriculture Canada.

Drosophilidae

Band, H.T., Bachli, G. & Band, R.N. (2005). Behavioral constancy for interspecies dependency enables Nearctic *Chymomyza amoena* (Loew) (Diptera: Drosophilidae) to spread in orchards and forests in Central and Southern Europe. *Biological Invasions, 7*(3), 509-530. doi: https://doi.org/10.1007/s10530-004-6352-2

Church, S.H., Extavour, C.G. & Rosenberg, M. (2022). Phylotranscriptomics reveals discordance in the phylogeny of Hawaiian *Drosophila* and *Scaptomyza* (Diptera: Drosophilidae). *Molecular Biology and Evolution, 39*(3). doi: https://doi.org/10.1093/molbev/msac012

Collinge, S.K. and Louda, S.M. (1989). *Scaptomyza nigrita* Wheeler (Diptera: Drosophilidae), a leaf miner of the native crucifer, *Cardamine cordifolia* A. Gray (Bittercress). *Journal of the Kansas Entomological Society, 62*(1), 1-10.

Debban, C.L. & Dyer, K.A. (2013). No evidence for behavioural adaptations to nematode parasitism by the fly *Drosophila putrida*. *Journal of Evolutionary Biology, 26*(8), 1646–1654. doi: https://doi.org/10.1111/jeb.12158

Dyer, K.A. (2012). Local selection underlies the geographic distribution of sex-ratio drive in *Drosophila neotestacea*. *Evolution, 66*(4), 973-984. doi: https://doi.org/10.1111/j.1558-5646.2011.01497.x

Dyer, KA., White, B.E., Sztepanacz, J.L., Bewick, E.R. & Rundle, H.D. (2014). Reproductive character displacement of epicuticular compounds and their contribution to mate choice in *Drosophila subquinaria* and *Drosophila recens*. *Evolution, 68*(4), 1163-1175. doi: https://doi.org/10.1111/evo.12335

Eberhard, W.G. (2002). Natural history and behavior of *Chymomyza mycopelates* and *C. exophthalma* (Diptera: Drosophilidae), and allometry of structures used as signals, weapons, and spore collectors. *Canadian entomologist, 134*(5), 667-687. doi: https://doi.org/10.4039/Ent134667-5

Ellis, W.N. (2020). *Parasites*. Plant Parasites of Europe. https://bladmineerders.nl/parasites/

Jaenike, J. (1985). Genetic and environmental determinants of food preference in *Drosophila tripunctata*. *Evolution, 39*(2), 362-369. doi: https://doi.org/10.1111/j.1558-5646.1985.tb05673.x

James, A.C., Jakubczak, J., Riley, M.P. & Jaenike, J. (1988). On the causes of monophagy in *Drosophila quinaria*. *Evolution, 42*(3), 626-630. doi: https://doi.org/10.1111/j.1558-5646.1988.tb04166.x

Jaramillo, S.L., Mehlferber, E. & Moore, P.J. (2015). Life-history trade-offs under different larval diets in *Drosophila suzukii* (Diptera: Drosophilidae). *Physiological Entomology, 40*(1), 2-9. doi: https://doi.org/10.1111/phen.12082

Lapoint, R.T., O'Grady, P.M. & Whiteman, N.K. (2013). Diversification and dispersal of the Hawaiian Drosophilidae: The evolution of *Scaptomyza*. *Molecular Phylogenetics and Evolution, 69*(1), 95-108. doi: https://doi.org/10.1016/j.ympev.2013.04.032

Muona, O. & Lumme, J. (1981). Geographical variation in the reproductive cycle and photoperiodic diapause of *Drosophila phalerata* and *D. transversa* (Drosophilidae: Diptera). *Evolution, 35*(1), 158-167. doi: https://doi.org/10.2307/2407949

Rotheray, G. & Lyszkowski, R. (2015). Diverse mechanisms of feeding and movement in Cyclorrhaphan larvae (Diptera). *Journal of Natural History, 49*(35-36), 2139-2211. doi: https://doi.org/10.1080/00222933.2015.1010314

Sampson, B.J., Mallette, T., Addesso, K.M., K.M., Liburd, O.E., Iglesias, L.E., J. Stringer, S.J., Werle, C.T., Shaw, D.A., Larsen, D. & Adamczyk, J.J. (2016). Novel aspects of *Drosophila suzukii* (Diptera: Drosophilidae) biology and an improved method for culturing this invasive species with a modified *D. melanogaster* diet. *Florida Entomologist, 99*(4), 774-780. doi: https://doi.org/10.1653/024.099.0433

Scudder, G.G.E. & Cannings, R.A. (2006). The Diptera families of British Columbia. http://www.for.gov.bc.ca/hfd/library/FIA/2006/FSP_Y062001b.pdf

Spicer, G.S. & Jaenike, J. (1996). Phylogenetic analysis of breeding site use and a- amanitin tolerance within the *Drosophila quinaria* species group. *Evolution, 50*(6), 2328–2337. doi: https://doi.org/10.1111/j.1558-5646.1996.tb03620.x

Stalker, H.D. (1945). On the biology and genetics of *Scaptomyza graminum* fallen (Diptera, Drosophilidae). *Genetics, 30*(3), 266–279. doi: https://doi.org/10.1093/genetics/30.3.266

Tidon, R. & de Almeida, J.M. (2016). Family Drosophilidae. *Zootaxa, 4122*(1), 719-751. doi: http://dx.doi.org/10.11646/zootaxa.4122.1.63

Trajković, J., Vujić, V., Miličić, D., Gojgić-Cvijović, G., Pavković-Lučić, S. & Savić, T. (2017). Fitness traits of *Drosophila melanogaster* (Diptera: Drosophilidae) after long-term laboratory rearing on different diets. *European Journal of Entomology, 144*(1), 222-229. doi: https://doi.org/10.14411/eje.2017.027

Peñafiel-Vinueza, A.D. & Rafael, V. (2018). Five new species of *Drosophila guarani* group from the Andes of southern Ecuador (Diptera, Drosophilidae). *ZooKeys, 781,* 141-163. doi: https://doi.org/10.3897/zookeys.781.22841

Wheeler, M.R. (1987). Drosophilidae. In McAlpine, J.F., Peterson, B.V., Shewell, G.E., Teskey, H.J., Vockeroth, J.R. & Wood, D.M. (Eds), Manual of Nearctic Diptera. Volume 2. (pp. 1011 – 1018). Research Branch Agriculture Canada.

Worthen, W.B., Jones, M.T. & Jetton, R.M. (1998). Community structure and environmental stress: desiccation promotes nestedness in mycophagous fly communities [*Drosophila tripunctata, Drosophila putrida*, nested-subset pattern, depauperate assemblage]. *Oikos, 81*(1), 45-54. doi: https://doi.org/10.2307/3546466

Živković, I.P., Barić, B., Šubić, M., Seljak, G. & Mešić, A. (2017). First record of alien species *Chymomyza amoena* [Diptera: Drosophilidae] in Croatia. *Šumarski list, 141*(9-10). doi: https://doi.org/10.31298/sl.141.9-10.6

Dryomyzidae

Gibson, J.F. & Choong, H.H.C. (2020). New range records and life history observations of insects (Diptera: Dryomyzidae, Chironomidae; Coleoptera: Staphylinidae) associated with barnacles (Balanomorpha: Balanidae, Chthamalidae) on the Pacific coasts of North America and Japan. *Canadian Entomologist, 153*(2), 196-210. doi: https://doi.org/10.4039/tce.2020.69

Scudder, G.G.E. & Cannings, R.A. (2006). The Diptera families of British Columbia. http://www.for.gov.bc.ca/hfd/library/FIA/2006/FSP_Y062001b.pdf

Steyskal, G.C. (1987). Dryomyzidae. In McAlpine, J.F., Peterson, B.V., Shewell, G.E., Teskey, H.J., Vockeroth, J.R. & Wood, D.M. (Eds), Manual of Nearctic Diptera. Volume 2. (pp. 923 – 926). Research Branch Agriculture Canada.

Wachkoo, A.A., Kurahashi, H., Khurshid, N. & Akbar, S.A. (2017). First record of *Dryomyza pakistana* Kurahashi, 1989 (Diptera, Dryomyzidae) from India. *Oriental Insects, 52*(1), 96-100. doi: https://doi.org/10.1080/00305316.2017.1377644

Empididae

Akbar, S.A., Kanturski, M., Barták, M., Wachkoo, A.A. & Maqbool, A. (2022). SEM studies and discovery of an intriguing new *Rhamphomyia* (*Pararhamphomyia*) (Diptera, Empididae, Empidinae)

species from the Kashmir Himalayas. *The European Zoological Journal, 89*(1), 1325-1350. doi: https://doi.org/10.1080/24750263.2022.2139864

Evans, H.E. (1988). Observations on swarms of *Rhamphomyia sociabilis* (Williston) (Diptera: Empididae). *Journal of the New York Entomological Society, 96*(3), 316-322.

Ivkovic, M., Perovic, M., Grootaert, P. & Pollet, M. (2021). High endemicity in aquatic dance flies of Corsica, France (Diptera, Empididae, Clinocerinae and Hemerodromiinae), with the description of a new species of *Chelipoda*. *ZooKeys, 1039*(1039), 177-197. doi: https://doi.org/10.3897/zookeys.1039.66493

Khruleva, O. A., Shamshev, I. V. & Sinclair, B. J. (2021). The Empidoid Flies (Diptera: Brachystomatidae, Empididae, Hybotidae) of Wrangel Island (Chukotka Autonomous Okrug): composition and distribution of the fauna. *Entomological Review, 101*(6), 792-819. doi: https://doi/org/10.1134/S0013873821060063

Kriska, G. (2013). *Freshwater Invertebrates in Central Europe*. Springer. doi: https://doi.org/10.1007/978-3-7091-1547-3_22

Newkirk, M.R. (1970). Biology of the longtailed dance fly, *Rhamphomyia longicauda* (Diptera: Enipididae); a new look at swarming. *Annals of the Entomological Society of America, 63*(5), 1407–1412. doi: https://doi.org/10.1093/aesa/63.5.1407

Rafael, J.A. & Câmara, J.T. (2016). Family Empididae. *Zootaxa, 4122*(1), 393-396. doi: http://dx.doi.org/10.11646/zootaxa.4122.1.34

Scudder, G.G.E. & Cannings, R.A. (2006). The Diptera families of British Columbia. http://www.for.gov.bc.ca/hfd/library/FIA/2006/FSP_Y062001b.pdf

Shamshev, I.V. & Selitskaya, O.G. (2016). Methyl salicylate as an attractant for the dance fly *Rhamphomyia gibba* (Fallen) (Diptera, Empididae). *Entomological Review, 96*(8), 1003-1007. doi: https://doi.org/10.1134/S0013873816080054

Shamshev, I.V. & Sinclair, B.J. (2009). Revision of the *Iteaphila setosa* group (Diptera: Empididae). *European Journal of Entomology, 106*(3), 441-450. doi: https://doi.org/10.14411/eje.2009.055

Steyskal, G.C. & Knutson, L.V. (1981). Empididae. In McAlpine, J.F., Peterson, B.V., Shewell, G.E., Teskey, H.J., Vockeroth, J.R. & Wood, D.M. (Eds), Manual of Nearctic Diptera. Volume 1. (pp. 607 – 624). Research Branch Agriculture Canada.

Ephydridae

Bownes, A. (2018). Effects of *Hydrilla verticillata* (L.f.) Royle (Hydrocharitaceae) growing conditions and nutrient content on the performance of a leaf-mining fly, *Hydrellia purcelli* Deeming (Diptera: Ephydridae). *Biocontrol Science and Technology, 28*(3), 278-292. doi: https://doi.org/10.1080/09583157.2018.1441370

Bownes, A. & Deeming, J. (2016). A new species of *Hydrellia* (Diptera: Ephydridae) mining *Hydrilla verticillata* (Hydrocharitaceae) leaves in Singapore. *Austral Entomology, 55*(4), 353-359. doi: https://doi-org/10.1111/aen.12193

Buckingham, G.R., Okrah, E.A. & Christian-Meier, M. (1991). Laboratory biology and host range of *Hydrellia balciunasi* [Diptera: Ephydridae]. *Entomophaga, 36*, 575-586. doi: https://doi.org/10.1007/BF02374440

Eiseman, C.S. & Zatwarnicki, T. (2019). First nearctic eecord of *Hydrellia albilabris* (Meigen) (Diptera: Ephydridae), a leafminer of duckweed (Araceae: Lemnoideae), with comments on related species. *Proceedings of the Entomological Society of Washington, 121*(2), 160-167. doi: https://doi.org/10.4289/0013-8797.121.2.160

Ellis, W.N. (2020). *Parasites.* Plant Parasites of Europe. https://bladmineerders.nl/parasites/

Purcell, M.F., Brown, B.T., Harms, N.E., Sun-Hee, H. & McCulloch, G.A. (2021). Biology and preliminary host range of a Korean leaf-mining *Hydrellia* sp. (Diptera: Ephydridae) rejected as a potential biological control agent for monoecious *Hydrilla verticillata* in the United States. *Biocontrol Science and Technology, 31*(4), 343-356. doi: https://doi.org/10.1080/09583157.2020.1853051

Kahanpää, J. & Zatwarnicki, T. (2015). Notes on Shore Flies (Diptera: Ephydridae) from Finland and north-western Russia. *Biodiversity Data Journal, 3.* doi: https://doi.org/10.3897/BDJ.3.e4701

Krishnaswamy, S. & Chacko, M.J. (1990). *Hydrellia* spp. [Diptera: Ephydridae] attacking *Hydrilla verticillata* in south India. *Entomophaga, 35*, 211-216. doi: https://doi.org/10.1007/BF02374795

Kriska, G. (2013). *Freshwater Invertebrates in Central Europe*. Springer. doi: https://doi.org/10.1007/978-3-7091-1547-3_22

Martin, G.D., Coetzee, J.A. & Baars, J-R. (2013). *Hydrellia lagarosiphon* Deeming (Diptera: Ephydridae), a potential biological control agent for the submerged aquatic weed, *Lagarosiphon major* (Ridl.) Moss ex Wager (Hydrocharitaceae). *African Entomology, 21*(1), 151-160. doi: https://doi.org/10.4001/003.021.0118

Mangan, R. & Baars, J.-R. (2023). Risk assessment of the host range of *Hydrellia lagarosiphon* for the biological control of *Lagarosiphon* major in Ireland. *Biocontrol Science and Technology, 33*(7), 681-700. doi: https://doi.org/10.1080/09583157.2023.2215993

Mangan, R., Dirilgen, T. & Baars, J-R. (2015). Responses of adult *Hydrellia lagarosiphon* to a revised diet: implications for life cycle studies and laboratory culturing techniques. *Entomologia Experimentalis et Applicata, 157*(2), 164-169. doi: https://doi.org/10.1111/eea.12350

Morton, A. & García del Pino, F. (2007). Susceptibility of shore fly *Scatella stagnalis* to five entomopathogenic nematode strains in bioassays. *BioControl, 52*(4), 533-545. doi: https://doi.org/10.1007/s10526-006-9047-z

Scheiring, J.F. (1974). Diversity of shore flies (Diptera: Ephydridae) in inland freshwater habitat. *Journal of the Kansas Entomological Society, 47*(4), 485-491.

Scheiring, J.F. (1975). A multivariate analysis of similarity in shore fly habitats (Diptera: Ephydridae). *Journal of the Kansas Entomological Society, 48*(2), 232-243.

Scheiring, J.F. (1977). Geographic variation in *Scatella stagnalis* (Diptera: Ephydridae) [North America, insects]. *Annals of the Entomological Society of America, 70*(4), 511-523. doi: https://doi.org/10.1093/aesa/70.4.511

Scudder, G.G.E. & Cannings, R.A. (2006). The Diptera families of British Columbia. http://www.for.gov.bc.ca/hfd/library/FIA/2006/FSP_Y062001b.pdf

Smith, R., Mangan, R. & Coetzee, J.A. (2019). Risk assessment to interpret the physiological host range of *Hydrellia egeriae*, a biocontrol agent for *Egeria densa*. *BioControl, 64*(4), 447-456. doi: https://doi.org/10.1007/s10526-019-09942-4

Walsh, G.C., Dalto, Y.M., Mattioli, F.M., Carruthers, R.I. & Anderson, L.W. (2013). Biology and ecology of Brazilian elodea *(Egeria densa*) and its specific herbivore*, Hydrellia* sp., in Argentina. *BioControl, 58*(1), 133-147. doi: https://doi.org/10.1007/s10526-012-9475-x

Wirth, W.W., Mathis, W.N. & Vockeroth, J.R. (1987). Ephydridae. In McAlpine, J.F., Peterson, B.V., Shewell, G.E., Teskey, H.J., Vockeroth, J.R. & Wood, D.M. (Eds), Manual of Nearctic Diptera. Volume 2. (pp. 1027 – 1048). Research Branch Agriculture Canada.

Fanniidae

Coupland, J.B. & Barker, G.M. (2004). Diptera as predators and parasitoids of terrestrial gastropods, with emphasis on Phoridae, Calliphoridae, Sarcophagidae, Muscidae and Fanniidae. *Natural Enemies of Terrestrial Molluscs,* 85-158. doi: https://doi.org/10.1079/9780851993195.0085

Kang, S., Eun, C.U., Kim, D. & Suh, S.J. (2023). A taxonomic revision of the genus *Fannia* Robineau-Desvoidy (Diptera: Fanniidae) from Korea. *Journal of Asia-Pacific Biodiversity, 16*(4), 516-524. doi: https://doi.org/10.1016/j.japb.2023.03.005

Kriska, G. (2013). *Freshwater Invertebrates in Central Europe*. Springer. doi: https://doi.org/10.1007/978-3-7091-1547-3_22

Kutty, S.N., Bernasconi, M.V., Šifner, F. & Meier, R. (2007). Sensitivity analysis, molecular systematics and natural history evolution of Scathophagidae (Diptera: Cyclorrhapha: Calyptratae). *Cladistics, 23*(1), 64-83. doi: https://doi.org/10.1111/j.1096-0031.2006.00131.x

Michalski, M., Gadawski, P., Klemm, J. & Szpila, K. (2021). New species of soldier fly—*Sargus bipunctatus (*Scopoli, 1763) (Diptera: Stratiomyidae), recorded from a human corpse in Europe—A case report. *Insects, 12*(4), 302. doi: https://doi.org/10.3390/insects12040302

Mohr, R.M., Mullens, B.A. & Gerry, A.C. (2011). Evaluation of ammonia, human sweat, and bovine blood as attractants for the female canyon fly, *Fannia conspicua* (Diptera: Muscidae), in southern California. *Journal of Vector Ecology, 36*(1), 55-58. doi: https://doi.org/10.1111/j.1948-7134.2011.00140.x

Mullens, B.A. & Gerry, A.C. (2006). Life history and seasonal abundance of *Fannia benjamini* complex (Diptera: Muscidae) in Southern California. *Journal of Medical Entomology, 43*(2), 192-199. doi: https://doi.org/10.1603/0022-2585(2006)043[0192:LHASAO]2.0.CO;2

Velásquez, Y., Martínez-Sánchez, A. & Rojo, S. (2013). First record of *Fannia leucosticta* (Meigen) (Diptera: Fanniidae) breeding in human corpses. *Forensic Science International, 229*(1), p.e13-e15. doi: https://doi.org/10.1016/j.forsciint.2013.03.022

Zhang, C. & Gerry, A.C. (2015). Laboratory colonization, life history observations, and desiccation tolerance of the canyon fly *Fannia conspicua* (Diptera: Fanniidae). *Journal of Medical Entomology, 52*(4), 532–538. doi: https://doi.org/10.1093/jme/tjv042

Heleomyzidae

Ellis, W.N. (2020). *Parasites.* Plant Parasites of Europe. https://bladmineerders.nl/parasites/

Gill, G.D. & Peterson, B.V. (1987). Heleomyzidae. In McAlpine, J.F., Peterson, B.V., Shewell, G.E., Teskey, H.J., Vockeroth, J.R. & Wood, D.M. (Eds), Manual of Nearctic Diptera. Volume 2. (pp. 973 – 980). Research Branch Agriculture Canada.

Mun, S.Y. & Suh, S.J. (2019). Taxonomic revision of the genus *Suillia* Robineau-Desvoidy (Diptera: Heleomyzidae) from Korea. *Journal of Asia-Pacific Biodiversity, 12*(3), 400-406. doi: https://doi.org/10.1016/j.japb.2019.04.004

Preisler, J., Roháček, J. & Tkoč, M. (2022). The fauna of Heleomyzidae (Diptera) in the Gemer area (Central Slovakia). *Acta Musei Silesiae, Scientiae Naturales, 71*(2), 131-181. doi: https://doi.org/10.2478/cszma-2022-0007

Rotheray, G.E. (2012). Morphology of the puparium and breeding sites of eight species of Heleomyzidae (Diptera). *Journal of Natural History, 46*(33-34), 2075-2102. doi: https://doi.org/10.1080/00222933.2012.707241

Scudder, G.G.E. & Cannings, R.A. (2006). The Diptera families of British Columbia. http://www.for.gov.bc.ca/hfd/library/FIA/2006/FSP_Y062001b.pdf

Soszynska-Maj, A. & Woznica, A.J. (2016). A case study of Heleomyzidae (Diptera) recorded on snow in Poland with a review of their winter activity in Europe. *European Journal of Entomology, 113*(1), 279-294. doi: https://doi.org/10.14411/eje.2016.035

Tuno, N., Sagara, N. & Okadome, T. (2003). Territorial behaviour of *Suillia* males on basidiocarps of *Hebeloma radicosum* in central Japan. *Mycologist, 17*(3), 122-125. doi: https://doi.org/10.1017/S0269915X03002143

Hybotidae

Barták, M. & Kubík, S. (2013). Species of *Bicellaria* Macquart (Diptera: Hybotidae) of Europe, with descriptions of four new species. *Zootaxa, 3647*(2), 251-278. doi: http://dx.doi.org/10.11646/zootaxa.3647.2.2

Barták, M. & Kubík, S. (2015). Three new species of European *Platypalpus* (Diptera, Hybotidae). *ZooKeys, 470*(470), 145-155. doi: https://doi.org/10.3897/zookeys.470.8967

Cumming, J.M. & Cooper, B.E. (2012). The identity of *Micrempis anatolica* Chillcott and *M. bomboxynon* Chillcott (Diptera: Empididae), with remarks on allied nearctic species. *Canadian Entomologist, 121*(7), 565-568. doi: https://doi.org/10.4039/Ent121565-7

Gonzàlez, C.R., Elgueta, M. & Ale-Rocha, R. (2021). A catalog of the Hybotidae of Chile (Diptera: Empidoidea). *Zootaxa, 5005*(2), 161-174. doi: https://doi.org/10.11646/zootaxa.5005.2.3

Grootaert, P. & Shamshev, I.V. (2012). The fast-running flies (Diptera, Hybotidae, Tachydromiinae) of Singapore and adjacent regions. *European Journal of Taxonomy, 5*. doi: http://dx.doi.org/10.5852/ejt.2012.5

Khruleva, O. A., Shamshev, I. V. & Sinclair, B. J. (2021). The Empidoid Flies (Diptera: Brachystomatidae, Empididae, Hybotidae) of Wrangel Island (Chukotka Autonomous Okrug): composition and distribution of the fauna. *Entomological Review, 101*(6), 792-819. doi: https://doi.org/10.1134/S0013873821060063

Nagy, Z.T., Sonet, G., Mortelmans, J., Vandewynkel, C. & Grootaert, P. (2013). Using DNA barcodes for assessing diversity in the family Hybotidae (Diptera, Empidoidea). *ZooKeys, 365*, 263-278. doi: https://doi.org/10.3897/zookeys.365.6070

Keroplatidae

Falaschi, R.L. (2016). Family Keroplatidae. *Zootaxa, 4122*(1), 56-61. doi: http://dx.doi.org/10.11646/zootaxa.4122.1.11

Jakovlev, J., Salmela, J., Polevoi, A., Penttinen, J. & Vartija, N-A. (2014). Recent noteworthy findings of fungus gnats from Finland and northwestern Russia (Diptera: Ditomyiidae, Keroplatidae, Bolitophilidae and Mycetophilidae). *Biodiversity Data Journal, 2*. doi: https://doi.org/10.3897/BDJ.2.e1068

Salmela, J, & Suuronen, A. (2014). A new *Neoplatyura* Malloch from Finland (Diptera, Keroplatidae). *Biodiversity Data Journal, 2*. doi: https://doi.org/10.3897/BDJ.2.e1323

Sivinski, J. (1982). Prey attraction by luminous larvae of the fungus gnat *Orfelia fultoni*. *Ecological Entomology, 7*(4), 443-446. doi: https://doi.org/10.1111/j.1365-2311.1982.tb00686.x

Lauxaniidae

Broadhead, E.C. (1989). The species of *Poecilominettia*, *Homoeominettia* and *Floriminettia* (Diptera: Lauxaniidae) in Panama. *Bulletin of the British Museum (Natural History) Entomology, 58*, (185-226).

Hesler, L.S. & Fernandes, G.W. (2016). Capture of nontarget flies (Diptera: Lauxaniidae, Chloropidae, and Anthomyiidae) on traps baited with volatile chemicals in field-crop habitats. *Psyche, 2016,* 1-8. doi: https://doi.org/10.1155/2016/6938368

Merz, B. (2001). Two new species of *Lauxania* Latreille s. str. (Diptera, Lauxaniidae) from Southern Europe. *Revue suisse de zoologie, 108*, 441-453. doi: https://doi.org/10.5962/bhl.part.80154

Merz, B. (2004). Revision of the *Minettia fasciata* species-group (Diptera, Lauxaniidae). *Revue suisse de zoologie, 111*, 183-211. doi: https://doi.org/10.5962/bhl.part.80234

Pérusse, J.R. & Wheeler, T.A. (2012). Revision of the nearctic species of *Lauxanza* (Diptera: Lauxaniidae). *The Canadian Entomologist, 132*(4), 411-427. doi: https://doi.org/10.4039/Ent132411-4

Scudder, G.G.E. & Cannings, R.A. (2006). The Diptera families of British Columbia. http://www.for.gov.bc.ca/hfd/library/FIA/2006/FSP_Y062001b.pdf

Shewell, G.E. (1987). Lauxaniidae. In McAlpine, J.F., Peterson, B.V., Shewell, G.E., Teskey, H.J., Vockeroth, J.R. & Wood, D.M. (Eds), Manual of Nearctic Diptera. Volume 2. (pp. 951 – 964). Research Branch Agriculture Canada.

Limoniidae

Bayfield, N. (1979). Some effects of trampling on *Molophilus ater* (Meigen) (Diptera, Tipulidae). *Biological Conservation, 16*(3), 219-232. doi: https://doi.org/10.1016/0006-3207(79)90023-5

Bilalli, A., Ibrahimi, H., Musliu, M., Grapci-Kotori, L., Geci, D., Slavevska-Stamenkovic, V., Hinic, J., Mitic-Kopanja, D. & Keresztes, L. (2021). New records of the craneflies (Diptera: Limoniidae, Tipulidae) from the Western Balkans. *Journal of Entomological Research Society, 23*(2), 141-152. doi: https://doi.org/10.51963/jers.v23i2.1929

Daichi, K., Takeyuki, N. & Takuji, T. (2020). Taxonomic study of the genus *Epiphragma* of Japan (Diptera: Limoniidae). *Acta Entomologica Musei Nationalis Pragae, 60*(2), 449-461. doi: https://doi.org/10.37520/aemnp.2020.29

Hinton, H.E. (1954). On the structure and function of the respiratory horns of the pupae of the genus *Pseudolimnophila* (Diptera: Tipulidae). *Proceedings of the Royal Entomological Society of London. Series A, General Entomology, 29*(10-12), 135-140. doi: https://doi.org/10.1111/j.1365-3032.1954.tb01186.x

Kania, I., Wang, B. & Szwedo, J. (2015). *Dicranoptycha* Osten Sacken, 1860 (Diptera, Limoniidae) from the earliest Cenomanian Burmese amber. *Cretaceous Research, 52*, 522-530. doi: https://doi.org/10.1016/j.cretres.2014.03.002

Kolcsár, L-P., Ivković, M. & Ternjej, I. (2015). New records of Limoniidae and Pediciidae (Diptera) from Croatia. *ZooKeys, 513*(513), 23-37. doi: https://doi.org/10.3897/zookeys.513.10066

Kolcsár, L-P., Török, E. & Keresztes, L. (2015). A new species and new records of *Molophilus* Curtis, 1833 (Diptera: Limoniidae) from the Western Palaearctic Region. *Biodiversity Data Journal, 3.* doi: https://doi.org/10.3897/BDJ.3.e5466

Kopeć, K., Perkovsky, E., Skibińska, K. (2019). A New species of a Genus *Cheilotrichia* (Diptera: Limoniidae) from Baltic and Ukrainian Amber. *Annales Zoologici, 69*(2), 423-426. doi: https://doi.org/10.3161/00034541ANZ2019.69.2.009

Krivosheina, M.G. & Krivosheina, N.P. (2010). New data on morphology and ecology of Limoniid fly larvae of the genus *Metalimnobia* (Diptera, Limoniidae) developed in fungal substrates. *Entomological Review, 90*(6), 764-782. doi: https://doi.org/10.1134/S0013873810060138

Krivosheina, N.P. (2010). New data on the ecology and morphology of xylobiont larvae of the genus *Elephantomyia* Ost.-Sack. (Diptera, Limoniidae). *Entomological Review, 90*(5), 603-614. doi: https://doi.org/10.1134/S0013873810050076

Mabrouki, Y., Terec, A.B., Taybi, F.A., Dénes, A. & Keresztes, L. (2023). Taxonomic notes and key to the West Palearctic *Antocha* (Antocha) Osten Sacken, 1860 (Diptera, Limoniidae) with description of a new species from Morocco. *Biodiversity Data Journal, 11*. doi: https://doi.org/10.3897/BDJ.11.e103849

Özgül, O. & Koç, H. (2014). Four new species of Limoniidae (Diptera, Nematocera) from the Inner-West Anatolian Subregion of Turkey. *The Florida Entomologist, 97*(2), 620-625. doi: https://doi.org/10.1653/024.097.0238

Podenas, S. & Gelhaus, J. (2011). Three new species of Chioneinae crane flies (Diptera: Limoniidae) from North-Central Mongolia. *Proceedings of the Academy of Natural Sciences of Philadelphia, 161*(1), 73-86. doi: https://doi.org/10.1635/053.161.0105

Podenas, S., Podeniene, V., Kim, T-W., Kim, A-Y., Park, S-J. & Aukštikalnienė, R. (2020). A new species of *Elephantomyia* crane fly (Diptera, Limoniidae) from Jeju Island, South Korea. *ZooKeys, 966*, 41-55. doi: https://doi.org/10.3897/zookeys.966.48590

Podeniene, V., Podenas, S. & Gelhaus, J.K. (2004). First record of a crane fly larva (Diptera, Limoniidae: Chioneinae) from Baltic amber. *Annals of the Entomological Society of America, 97*(6), 1126-1128. doi: https://doi.org/10.1603/0013-8746(2004)097[1126:FROACF]2.0.CO2

Ribeiro, G.C. & Santos, D. (2016). Families Tipulidae and Limoniidae. *Zootaxa, 4122*(1), 73-97. doi: http://dx.doi.org/10.11646/zootaxa.4122.1.14

Salmela, J. & Starý, J. (2008). Description of *Metalimnobia* (*Metalimnobia*) *charlesi* sp. n. from Europe (Diptera, Limoniidae). *Entomologica Fennica, 19*(4), 268–272. doi: https://doi.org/10.33338/ef.84444

Starý, J. & Salmela, J. (2004). Redescription and biology of *Limonia badia* (Walker) (Diptera: Limoniidae). *Entomologica Fennica, 15*(1), 41–47. doi: https://doi.org/10.33338/ef.84205

Lonchopteridae

Coulson, J.C. & Butterfield, J. (1982). The distribution and biology of Lonchopteridae (Diptera) in upland regions of northern England. *Ecological Entomology, 7*(1), 31-38. doi: https://doi.org/10.1111/j.1365-2311.1982.tb00641.x

Rotheray, G. & Lyszkowski, R. (2015). Diverse mechanisms of feeding and movement in Cyclorrhaphan larvae (Diptera). *Journal of Natural History, 49*(35-36), 2139-2211. doi: https://doi.org/10.1080/00222933.2015.1010314

Stalker, H.D. (1956). On the evolution of parthenogenesis in *Lonchoptera* (Diptera). *Evolution, 10*(4), 345-359. doi: https://doi.org/10.1111/j.1558-5646.1956.tb02862.x

Micropezidae

Barnes, J.K. (2015). Biology and immature stages of *Compsobata univitta* (Walker, 1849) (Diptera: Micropezidae: Calobatinae). *Proceedings of the Entomological Society of Washington, 117*(4), 421-434. doi: https://doi.org/10.4289/0013-8797.117.4.421

Rohácek, J. (2013). The fauna of the Acalyptrate families Micropezidae, Psilidae, Clusiidae, Acartophthalmidae, Anthomyzidae, Aulacigastridae, Periscelididae and Asteiidae (Diptera) in the Gemer area (Central Slovakia): supplement 1. *Acta musei silesiae: scientiae naturales, 62*(2),125. doi: https://doi.org/10.2478/cszma-2013-0014

Scudder, G.G.E. & Cannings, R.A. (2006). The Diptera families of British Columbia. http://www.for.gov.bc.ca/hfd/library/FIA/2006/FSP_Y062001b.pdf

Steyskal, G.C. (1987). Micropezidae. In McAlpine, J.F., Peterson, B.V., Shewell, G.E., Teskey, H.J., Vockeroth, J.R. & Wood, D.M. (Eds), Manual of Nearctic Diptera. Volume 2. (pp. 761 – 768). Research Branch Agriculture Canada.

Milichiidae

Brochu, K. & Wheeler, T.A. (2009). Systematics and ecology of the Nearctic species of *Neophyllomyza* (Diptera: Milichiidae). *The Canadian Entomologist, 141*(2), 103 – 111. doi: https://doi.org/10.4039/n09-001

Levesque-Beaudin, V. & Mlynarek, J.J. (2020). Revision of Nearctic *Paramyia* Williston (Diptera: Milichiidae). *Zootaxa, 4732*(1), 1-56. doi: https://doi.org/10.11646/zootaxa.4732.1.1

Rohácek, J. (1996). Fourth supplement to the acalyptrate Diptera fauna of the Czech Republic and Slovakia. *Casopis Slezského musea v Opava, 45*, 17-28.

Rotheray, G. & Lyszkowski, R. (2015). Diverse mechanisms of feeding and movement in Cyclorrhaphan larvae (Diptera). *Journal of Natural History, 49*(35-36), 2139-2211. doi: https://doi.org/10.1080/00222933.2015.1010314

Sabrosky, C.W. (1987). Milichiidae. In McAlpine, J.F., Peterson, B.V., Shewell, G.E., Teskey, H.J., Vockeroth, J.R. & Wood, D.M. (Eds), Manual of Nearctic Diptera. Volume 2. (pp. 903 – 908). Research Branch Agriculture Canada.

Scudder, G.G.E. & Cannings, R.A. (2006). The Diptera families of British Columbia. http://www.for.gov.bc.ca/hfd/library/FIA/2006/FSP_Y062001b.pdf

Muscidae

Arntfield, P.W. (1975). A revision of *Graphomya* Robineau-Desvoidy (Diptera: Muscidae) from North America. *The Canadian Entomologist, 107*(3), 257 – 302. doi: https://doi.org/10.4039/Ent107257-3

Baleba, S.B.S., Torto, B., Masiga, D., Weldon, C.W. & Getahun, M.N. (2019). Egg-laying decisions based on olfactory cues enhance offspring fitness in *Stomoxys calcitrans* L. (Diptera: Muscidae). *Scientific Reports, 9*(1), 3850. doi: https://doi.org/10.1038/s41598-019-40479-9

Bautista-Martínez, N., Illescas-Riquelme, C.P. & García-Ávila, C., de J. (2017). First report of “hunter-fly” *Coenosia attenuata* (Diptera: Muscidae) in Mexico. *The Florida Entomologist, 100*(1), 174-175. doi: https://doi.org/10.1653/024.100.0126

Butterworth, N.J. & Wallman, J.F. (2022). Flies getting filthy: The precopulatory mating behaviours of three mud-dwelling species of Australian *Lispe* (Diptera: Muscidae). *Ethology, 128*(4), 369-377. doi: https://doi.org/10.1111/eth.13236

Chen, W., Shang, Y., Ren, L., Zhang, X. & Guo, Y. (2018). The complete mitochondrial genome of *Graphomya rufitibia* (Diptera: Muscidae). *Mitochondrial DNA Part B Resources, 3*(1), 403-404. doi: https://doi.org/10.1080/23802359.2018.1456375

Coupland, J.B. & Barker, G.M. (2004). Diptera as predators and parasitoids of terrestrial gastropods, with emphasis on Phoridae, Calliphoridae, Sarcophagidae, Muscidae and Fanniidae. *Natural Enemies of Terrestrial Molluscs,* 85-158. doi: https://doi.org/10.1079/9780851993195.0085

de Carvalho, C.J.B., Couri, M.S., Pont, A.C., Pamplona, D. & Lopes, S.M. (2005). A catalogue of the Muscidae (Diptera) of the neotropical region. *Zootaxa, 860*(1), 1-282. doi: https://doi.org/10.11646/zootaxa.860.1.1

Drummond, F.A., Groden, E., Haynes, D.L. & Edens, T.C. (1989). Some Aspects of the Biology of a Predaceous Anthomyiid Fly, *Coenosia Tigrina*. *The Great Lakes Entomologist, 22*(1). doi: https://doi.org/10.22543/0090-0222.1659

Duarte, J.L.P., Krüger, R.F. & Ribeiro, P.B. (2013). Interaction between *Musca domestica* L. and its predator *Muscina stabulans* (Fallén) (Diptera, Muscidae): effects of prey density and food source abundance. *Revista Brasileira de Entomologia, 57*(1), 55-58. doi: https://doi.org/10.1590/S0085-56262013000100009

Fowler, F.E. & Mullens, B.A. (2016). Dividing the pie: differential dung pat size utilization by sympatric *Haematobia irritans* and *Musca autumnalis*: Dung pat size use by horn flies and face flies. *Medical and Veterinary Entomology, 30*(2), 185-192. doi: https://doi.org/10.1111/mve.12166

Fryxell, R.T.T., Moon, R.D., Boxler, D.J. & Watson, D.W. (2021). Face fly (Diptera: Muscidae)—Biology, pest status, current management prospects, and research needs. *Journal of Integrated Pest Management, 12*(1), 5. doi: https://doi.org/10.1093/jipm/pmaa020

Giordani, G., Grzywacz, A. & Vanin, S. (2019). Characterization and identification of puparia of *Hydrotaea* Robineau-Desvoidy, 1830 (Diptera: Muscidae) from forensic and archaeological contexts. *Journal of Medical Entomology, 56*(1), 45–54. doi: https://doi.org/10.1093/jme/tjy142

Giordani, G., Tuccia, F., Floris, I. & Vanin, S. (2018). First record of *Phormia regina* (Meigen, 1826) (Diptera: Calliphoridae) from mummies at the Sant'Antonio Abate Cathedral of Castelsardo, Sardinia, Italy. *PeerJ, 6*. doi: https://doi.org/10.7717/peerj.4176

Gomes, L.R.P., Couri, M.S. & de Carvalho, C.J.B. (2018). Anthomyiidae, Fanniidae and Muscidae (Diptera) from the Juan Fernández Archipelago (Chile): 60 years after Willi Hennig's contributions. *Zootaxa, 4402*(2), 373-389. doi: https://doi.org/10.11646/zootaxa.4402.2.9

Grzywacz, A., Hall, M.J.R., Pape, T. & Szpila, K. (2017). Muscidae (Diptera) of forensic importance—an identification key to third instar larvae of the western Palaearctic region and a catalogue of the muscid carrion community. *International Journal of Legal Medicine, 131*(3), 855-866. doi: https://doi.org/10.1007/s00414-016-1495-0

Huckett, H.C. & Vockeroth, J.R. (1987). Muscidae. In McAlpine, J.F., Peterson, B.V., Shewell, G.E., Teskey, H.J., Vockeroth, J.R. & Wood, D.M. (Eds), Manual of Nearctic Diptera. Volume 2. (pp. 1115 – 1132). Research Branch Agriculture Canada.

Ivković, M. & Pont, A.C. (2016). Long-time emergence patterns of *Limnophora* species (Diptera, Muscidae) in specific karst habitats: tufa barriers. *Limnologica, 61*, 29-35. doi: https://doi.org/10.1016/j.limno.2016.09.003

Jarzen, D.M. & Hogsette, J.A. (2008). Pollen from the exoskeletons of stable flies, *Stomoxys calcitrans* (linnaeus 1758), in Gainesville, Florida, U.S.A. *Palynology, 32*(1), 77–81. doi: https://doi.org/10.2113/gspalynol.32.1.77

Krafsur, E.S. & Moon, R.D. (1997). Bionomics of the face fly, *Musca autumnalis*. *Annual Review of Entomology, 42*(1), 503-523. doi: https://doi.org/10.1146/annurev.ento.42.1.503

Kriska, G. (2013). *Freshwater Invertebrates in Central Europe*. Springer. doi: https://doi.org/10.1007/978-3-7091-1547-3_22

Krivosheina, N.P. (2013). On the ecology of *Phaonia* larvae (Diptera, Muscidae). *Entomological Review, 93*(3), 324-333. doi: https://doi.org/10.1134/S0013873813030068

Krivosheina, N.P. (2013). Morpho-ecological characteristics of the xylobiont larvae of the genus *Phaonia* Robineau-Desvoidy, 1830 (Diptera, Muscidae) with a description of the larva of *Ph. canescens* Stein, 1916. *Entomological Review, 93*(2), 249-257. doi: https://doi.org/10.1134/S0013873813020140

Krivosheina, N.P. (2013). New data on the morphology and ecology of *Phaonia* larvae with a description of the larva of *Phaonia wahlbergi* (Diptera, Muscidae). *Entomological Review, 93*(7), 925-934. doi: https://doi.org/10.1134/S0013873813070166

Krivosheina, N.P. (2014). Morpho-ecological characteristic of immature stages of the genus *Mydaea* (Diptera, Muscidae). *Entomological Review, 94*(5), 675-686. doi: https://doi.org/10.1134/S0013873814050042

Krivosheina, N.P. (2019). Morpho-ecological characteristics of immature stages of Xytobiont species of the genus *Phaonia* Robineau-Desvoidy, 1830 (Muscidae, Diptera) inhabiting tree holes and sap accumulations. Entomological Review, 99(2), 250-261. doi: https://doi.org/10.1134/S0013873819020118

Kutty, S.N., Bernasconi, M.V., Šifner, F. & Meier, R. (2007). Sensitivity analysis, molecular systematics and natural history evolution of Scathophagidae (Diptera: Cyclorrhapha: Calyptratae). *Cladistics, 23*(1), 64-83. doi: https://doi.org/10.1111/j.1096-0031.2006.00131.x

Kutty, S.N., Pont, A.C., Meier, R. & Pape, T. (2014). Complete tribal sampling reveals basal split in Muscidae (Diptera), confirms saprophagy as ancestral feeding mode, and reveals an evolutionary correlation between instar numbers and carnivory. *Molecular Phylogenetics and Evolution, 78*, 349-364. doi: https://doi.org/10.1016/j.ympev.2014.05.027

LeRoux, E.J. & Perron, J.P. (1960). Descriptions of immature stages of *Coenosia tigrina* (F.) (Diptera: Anthomyiidae), with notes on hibernation of larvae and predation by adults. *The Canadian Entomologist, 92*(4), 284 – 296. doi: https://doi.org/10.4039/Ent92284-4

Ma, T. & Huang, J. (2018). A new species of the genus *Morellia* Robineau-Desvoidy (Diptera: Muscidae) from Yunnan, China, with analysis of available DNA barcoding sequences. *Biologia,* 73(12), 1205-1213. doi: https://doi.org/10.2478/s11756-018-0134-2

Merritt, R.W. & Wotton, R.S. (1988). The life history and behavior of *Limnophora riparia* (Diptera: Muscidae), a predator of larval black flies. *Journal of the North American Benthological Society, 7*(1), 1-12. doi: https://doi.org/10.2307/1467826

Michalski, M., Gadawski, P., Klemm, J. & Szpila, K. (2021). New species of soldier fly—*Sargus bipunctatus (*Scopoli, 1763) (Diptera: Stratiomyidae), recorded from a human corpse in Europe—A case report. *Insects, 12*(4), 302. doi: https://doi.org/10.3390/insects12040302

Morris, D.E. & Cloutier, C. (1987). Biology of the predatory fly *Coenosza tigrina* (fab.) (Diptera: Anthomyiidae): reproduction, development, and larval feeding on earthworms in the laboratory. *The Canadian Entomologist, 119*(4), 381 – 393. doi: https://doi.org/10.4039/Ent119381-4

Patitucci, L.D., Migale, S. & Mulieri, P.R. (2020). The killer flies *Coenosia* Meigen (Diptera: Muscidae) of southern South America: Resolving the taxonomic puzzle of *Coenosia inaequalis* Malloch, 1934. *Zoologischer Anzeiger, 288*, 66-73. doi: https://doi-org/10.1016/j.jcz.2020.06.006

Perron, J.P., LeRoux, E.J. & Lafrance, J. (1956). Notes on *Coenosia tigrina* (F.) (Dipera: Anthomyiidae), mainly on habits and rearing. *The Canadian Entomologist, 88*(10), 608 – 611. doi: https://doi.org/10.4039/Ent88608-10

Pont, A.C. & Ivković, M. (2013). The Hunter-flies of Croatia (Diptera: Muscidae: genus *Limnophora* Robineau-Desvoidy). *Journal of Natural History, 47*(15-16), 1069-1082. doi: https://doi.org/10.1080/00222933.2012.750775

Rochon, K., Hogsette, J.A., Kaufman, P.E., Olafson, P.U., Swiger, S.L. & Taylor, D.B. (2021). Stable fly (Diptera: Muscidae)—Biology, management, and research needs. *Journal of Integrated Pest Management, 12*(1), 38. doi: https://doi.org/10.1093/jipm/pmab029

Rotheray, G.E. & Wilkinson, G. (2015). Trophic structure and function in the larvae of predatory muscid flies (Diptera, Muscidae). Zoomorphology, 134(4), 553-563. doi: https://doi.org/10.1007/s00435-015-0284-5

Savage, J. & Sorokina, V.S. (2021). Review of the North American fauna of *Drymeia* Meigen (Diptera, Muscidae) and evaluation of DNA barcodes for species-level identification in the genus. *ZooKeys, 1024*(5), 31-89. doi: https://doi.org/10.3897/zookeys.1024.60393

Savage, J., Wheeler, T.A. & Wiegmann, B.M. (2004). Phylogenetic analysis of the genus *Thricops rondani* (Diptera: Muscidae) based on molecular and morphological characters. *Systematic Entomology, 29*(3), 395-414. doi: https://doi.org/10.1111/j.0307-6970.2004.00252.x

Scudder, G.G.E. & Cannings, R.A. (2006). The Diptera families of British Columbia. http://www.for.gov.bc.ca/hfd/library/FIA/2006/FSP_Y062001b.pdf

Seabra, S.G., Martins, J., Bras, P., Tavares, A.M., Freitas, I., Barata, A., Rebelo, M.T., Mateus, C., Paulo, O.S. & Figueiredo, E. (2021). PCR-based detection of prey DNA in the gut contents of the tiger-fly, *Coenosia attenuata* (Diptera: Muscidae), a biological control agent in Mediterranean greenhouses. *European Journal of Entomology, 118*(1), 335-343. doi: https://doi.org/10.14411/eje.2021.035

Sommer, C., Jensen, K.-M.V. & Jespersen, J.B. (2001). Topical treatment of calves with synthetic pyrethroids: effects on the non-target dung fly *Neomyia cornicina* (Diptera: Muscidae). *Bulletin of Entomological Research, 91*(2), 131-137. doi: https://doi.org/10.1079/BER200079

Sorokina, V.S. (2022). New taxonomic notes on the genus *Coenosia* Meigen (Diptera: Muscidae), with the description of four new species from North-East Russia and the Altai Mountains. *International Journal of Entomology, 58*(1), 43-62. doi: https://doi.org/10.1080/00379271.2022.2027270

Teskey, H.J. (1960). A review of the life-history and habits of *Musca autumnalis* DeGeer (Diptera: Muscidae). *The Canadian Entomologist, 92*(5), 360 – 367. doi: https://doi.org/10.4039/Ent92360-5

Vikhrev, N.E. (2011). Review of the Palaearctic members of the *Lispe tentaculata* species-group (Diptera, Muscidae): revised key, synonymy and notes on ecology. *ZooKeys, 84*(84), 59-70. doi: https://doi.org/10.3897/zookeys.84.819

Vikhrev, N.E. (2012). Revision of the *Lispe longicollis*-group (Diptera, Muscidae). *ZooKeys, 235*(235), 23-39. doi: https://doi.org/10.3897/zookeys.235.3306

Xue, W-Q., Zhang, L. & Wang, M-F. (2009). Study of the genus *Spilogona* Schnabl (Diptera: Muscidae) from China, with descriptions of four new species. *Proceedings of the Entomological Society of Washington, 111*(2), 530-538. doi: https://doi.org/10.4289/0013-8797-111.2.530

Zou, D., Coudron, T.A., Xu, W., Xu, J. & Wu, H. (2020). Performance of the tiger-fly *Coenosia attenuata* Stein reared on the alternative prey, *Chironomus plumosus* (L.) larvae in coir substrate. *Phytoparasitica, 49*(1), 83-92. doi: https://doi.org/10.1007/s12600-020-00866-9

Zou, D., Coudron, T.A., Zhang, L., Xu, W., Xu, J., Wang, M., Xiao, X. & Wu, H. (2021). Effect of prey species and prey densities on the performance of adult *Coenosia attenuata*. *Insects, 12*(8), 669. doi: https://doi.org/10.3390/insects12080669

Mycetophilidae

Deady, R.J., Delaney, M.A., Jones, E. & Chandler, P.J. (2022). Further interceptions of the Neotropical fungus gnat *Sciophila fractinervis* Edwards, 1940 (Diptera, Mycetophilidae) in Britain with comments and observations on its biology and spread. *Biodiversity Data Journal, 10*. doi: https://doi.org/10.3897/BDJ.10.e94812

Jakovlev, J. & Penttinen, J. (2007). *Boletina dispectoides* sp.n. and six other species of fungus gnats (Diptera: Mycetophilidae) new to Finland. *Entomologica Fennica, 18*(4), 211–217. doi: https://doi.org/10.33338/ef.84401

Jürgenstein, S., Kurina, O. & Põldmaa, K. (2015). The *Mycetophila ruficollis* Meigen (Diptera, Mycetophilidae) group in Europe: elucidating species delimitation with COI and ITS2 sequence data. *ZooKeys, 508*(508), 15-51. doi: https://doi.org/10.3897/zookeys.508.9814

Kurina, O., Kjærandsen, J., Kirik, H., Hadbavná, D., Dénes, A., Oboňa, J. & Manko, P. (2023). On the identity and distribution of the rare *Rymosia tolleti* Burghele-Balacesco, 1965 (Diptera, Mycetophilidae) encountered in European caves. *Check List, 19*(3), 381-389. doi: https://doi.org/10.15560/19.3.381

Lindemann, J.P., Søli, G. & Kjærandsen, J. (2021). Revision of the *Exechia* parva group (Diptera: Mycetophilidae). *Biodiversity Data Journal, 9*. doi: https://doi.org/10.3897/BDJ.9.e67134

Pessacq, P., Omad, G., Kerr, P.H. & Pardo, C. (2017). First studies on Patagonian immature Mycetophilidae: description of the larva and pupa, redescription and comments on the biology of *Mycomya chilensis*. *Revista Mexicana de Biodiversidad, 88*(4), 815-819. doi: https://doi.org/10.1016/j.rmb.2017.10.022

Salmela, J., Suuronen, A. & Kaunisto, K.M. (2016). New and poorly known Holarctic species of *Boletina* Staeger, 1840 (Diptera, Mycetophilidae). *Biodiversity Data Journal, 4*. doi: https://doi.org/10.3897/BDJ.4.e7218

Scudder, G.G.E. & Cannings, R.A. (2006). The Diptera families of British Columbia. http://www.for.gov.bc.ca/hfd/library/FIA/2006/FSP_Y062001b.pdf

Vockeroth, J.R. (1981). Mycetophilidae. In McAlpine, J.F., Peterson, B.V., Shewell, G.E., Teskey, H.J., Vockeroth, J.R. & Wood, D.M. (Eds), Manual of Nearctic Diptera. Volume 1. (pp. 223 – 246). Research Branch Agriculture Canada.

Opomyzidae

Ellis, W.N. (2020). *Parasites*. Plant Parasites of Europe. https://bladmineerders.nl/parasites/

Roháček, J. (2012). The fauna of the opomyzoid families Clusiidae, Acartophthalmidae, Anthomyzidae, Opomyzidae, Stenomicridae, Periscelididae, Asteiidae (Diptera) in the Gemer area (Central Slovakia). *Časopis Slezského Zemského Muzea, 61*(2), 97-111. doi: https://doi.org/10.2478/v10210-012-0011-5

Scudder, G.G.E. & Cannings, R.A. (2006). The Diptera families of British Columbia. http://www.for.gov.bc.ca/hfd/library/FIA/2006/FSP_Y062001b.pdf

Thomas, I. (1938). On the bionomics and structure of some Dipterous larvae infesting cereals and grasses. *Annals of Applied Biology, 25*(1), 181-196. doi: https://doi.org/10.1111/j.1744-7348.1938.tb04356.x

Vockeroth, J.R. (1961). The North American Species of the Family Opomyzidae (Diptera : Acalypterae). *The Canadian Entomologist, 93*(7), 503 – 522. doi: https://doi.org/10.4039/Ent93503-7

Vockeroth, J.R. (1987). Opomyzidae. In McAlpine, J.F., Peterson, B.V., Shewell, G.E., Teskey, H.J., Vockeroth, J.R. & Wood, D.M. (Eds), Manual of Nearctic Diptera. Volume 2. (pp. 881 – 886). Research Branch Agriculture Canada.

Oestridae

Colwell, D.D., Hall, M.J.R. & Scholl, P.J. (2006). *The oestrid flies: biology, host-parasite relationships, impact and management*. CABI Publishing. doi: https://doi.org/10.1079/9780851996844.0000

El-Hawagry, M.S.A., Abdel-Dayem, M.S. & Al Dhafer, H.M. (2020). The family Oestridae in Egypt and Saudi Arabia (Diptera, Oestroidea). *ZooKeys, 947*(3), 113-142. doi: https://doi.org/10.3897/zookeys.947.52317

Rukke, B., Cholidis, S., Johnsen, A. & Ottesen, P. (2014). Confirming *Hypoderma tarandi* (Diptera: Oestridae) human ophthalmomyiasis by larval DNA barcoding. *Acta Parasitologica, 59*(2), 301-304. doi: http://doi.org/10.2478/s11686-014-0242-2

Scudder, G.G.E. & Cannings, R.A. (2006). The Diptera families of British Columbia. http://www.for.gov.bc.ca/hfd/library/FIA/2006/FSP_Y062001b.pdf

Wood, D.M. (1987). Oestridae. In McAlpine, J.F., Peterson, B.V., Shewell, G.E., Teskey, H.J., Vockeroth, J.R. & Wood, D.M. (Eds), Manual of Nearctic Diptera. Volume 2. (pp. 1147 – 1158). Research Branch Agriculture Canada.

Pediciidae

Ferreira, S., Oosterbroek, P., Stary, J., Sousa, P., Mata, V.A., da Silva, L.P., Pauperio, J. & Beja, P. (2021). The InBIO Barcoding Initiative Database: DNA barcodes of Portuguese Diptera 02-Limoniidae, Pediciidae and Tipulidae. *Biodiversity Data Journal*, *9*(4), p.e69841. doi: https://doi.org/10.3897/BDJ.9.e69841

Kolcsár, L-P., Oosterbroek, P., Olsen, K.M., Paramonov, N.M., Gavryushin, D.I., Pilipenko, V.E., Polevoi, A.V., Eiroa, E., Andersson, M., Dufour, C., Syratt, M., Kurina, O., Lindström, M., Starý, J., Lantsov, V.I., Wiedeńska, J., Pape, T., Friman, M., Peeters, K., Gritsch, W., Salmela, J., Viitanen, E., Aristophanous, M., Janević, D. & Watanabe, K. (2023). Contribution to the Knowledge of Cylindrotomidae, Pediciidae and Tipulidae (Diptera: Tipuloidea): First Records of 86 Species from Various European Countries. *Diversity, 15*(3), 336. doi: https://doi.org/10.3390/d15030336

Krivosheina, N.P. (2011). New data on the larval morphology of limoniid flies of the genus *Ula* (Diptera, Pediciidae). *Entomological Review, 91*(4), 432-443. doi: https://doi.org/10.1134/S001387381104004X

Phoridae

Ament, D.C. & Brown, B.V. (2016). Family Phoridae. *Zootaxa, 4122*(1), 414-451. doi: http://dx.doi.org/10.11646/zootaxa.4122.1.37

Ament, D.C. & dos Santos, T.G. (2017). Taxonomy and first records of two *Megaselia Rondani* species (Diptera: Phoridae) preying upon eggs of *Phyllomedusa iheringii* Boulenger (Anura: Phyllomedusidae). *Neotropical Entomology, 46*(3)289-294. doi: http://doi.org/10.1007/s13744-016-0462-2

Brown, B.V. & Emily A. Hartop, E.A. (2017). Mystery mushroom malingerers: *Megaselia marquezi* Hartop et al. 2015 (Diptera: Phoridae). *Biodiversity Data Journal, 5*. doi: https://doi.org/10.3897/BDJ.5.e15052

Brian V. Brown, B.V. & Philpott, S.M. (2012). *Pseudacteon* parasitoids of *Azteca instabilis* ants in Southern Mexico (Diptera: Phoridae; Hymenoptera: Formicidae). *Psyche*, 1-6. doi: https://doi.org/10.1155/2012/351232

Brown, B.V., Wong, M.A. & Hartop, E. (2019). A new white-spotted *Megaselia Rondani* (Diptera: Phoridae) from western North America. *Biodiversity Data Journal, 7*. doi: https://doi.org/10.3897/BDJ.7.e34310

Chen, L. & Fadamiro, H.Y. (2018). *Pseudacteon* phorid flies: Host specificity and impacts on *Solenopsis* fire ants. *Annual Review of Entomology, 63*, 47-67. doi: https://doi.org/10.1146/annurev-ento-020117-043049

Chen, L. & Porter, S.D. (2020). Biology of *Pseudacteon* decapitating flies (Diptera: Phoridae) that parasitize ants of the *Solenopsis saevissima* complex (Hymenoptera: Formicidae) in South America. *Insects, 11*(2), 107. doi: https://doi.org/10.3390/insects11020107

Cônsoli, F.L, Wuellner, C.T., Vinson, S.B. & Gilbert, L.E. (2001). Immature development of *Pseudacteon tricuspis* (Diptera: Phoridae), an endoparasitoid of the red imported fire ant (Hymenoptera: Formicidae). *Annals of the Entomological Society of America, 94*(1), 97–109. doi: https://doi.org/10.1603/0013-8746(2001)094[0097:IDOPTD]2.0.CO;2

Coupland, J.B. & Barker, G.M. (2004). Diptera as predators and parasitoids of terrestrial gastropods, with emphasis on Phoridae, Calliphoridae, Sarcophagidae, Muscidae and Fanniidae. *Natural Enemies of Terrestrial Molluscs,* 85-158. doi: https://doi.org/10.1079/9780851993195.0085

Disney, R.H.L. (2007). Natural history of the scuttle fly, *Megaselia scalaris*. *Annual Review of Entomology, 53*(1), 39-60. doi: https://doi.org/10.1146/annurev.ento.53.103106.093415

Disney, R.H.L. (2013). An unusually rich scuttle fly fauna (Diptera, Phoridae) from north of the Arctic Circle in the Kola Peninsula, N. W. Russia. *ZooKeys, 342*, 45-74. doi: https://doi.org/10.3897/zookeys.342.5772

Ellis, W.N. (2020). *Parasites*. Plant Parasites of Europe. https://bladmineerders.nl/parasites/

Folgarait, P.J., Chirino, M.G., Patrock, R.J.W. & Gilbert, L.E. (2005). Development of *Pseudacteon obtusus* (Diptera: Phoridae) on *Solenopsis invicta* and *Solenopsis richteri* fire ants (Hymenoptera: Formicidae). *Environmental Entomology, 34*(2), 308–31. doi: https://doi.org/10.1603/0046-225X-34.2.308

Folgarait, P.J., Plowes, R.M., Gomila, C. & Gilbert, L.E. (2020). A small parasitoid of fire ants, *Pseudacteon obtusitus* (Diptera: Phoridae): native range ecology and laboratory rearing. *Florida Entomologist, 103*(1), 9-1. doi: https://doi.org/10.1653/024.103.0402

Morrison, L.W. & Gilbert, L.E. (2004). Parasitoid–host relationships when host size varies: the case of *Pseudacteon* flies and *Solenopsis* fire ants. *Ecological Entomology, 23*(4), 409-416. doi: https://doi.org/10.1046/j.1365-2311.1998.00159.x

Pesquero, M.A., Vaz, A.P.de A., & de Arruda, F.V. (2013). Laboratory rearing and niche resources of *Pseudacteon* spp. Coquillett (Diptera: Phoridae) parasitoids of *Solenopsis saevissima* (Smith)

(Hymenoptera: Formicidae). *Sociobiology*, *60*(4), 484–486. doi: https://doi.org/10.13102/sociobiology.v60i4.484-486

Peterson, B.V. (1987). Phoridae. In McAlpine, J.F., Peterson, B.V., Shewell, G.E., Teskey, H.J., Vockeroth, J.R. & Wood, D.M. (Eds), Manual of Nearctic Diptera. Volume 2. (pp. 689 – 712). Research Branch Agriculture Canada.

Polidori, C., Disney, R.H.L. & Andrietti, F. (2004). Some observations on the reproductive biology of the scuttle fly *Megaselia andrenae* (Diptera: Phoridae) at the nesting site of its host *Andrena agilissima* (Hymenoptera: Andrenidae). *European Journal of Entomology, 101*(2), 337-340. doi: https://doi.org/10.14411/eje.2004.045

Porter, S.D. & Plowes, R.M. (2018). Rearing and biology of the decapitating fly *Pseudacteon bifidus* (Diptera: Phoridae): A parasitoid of tropical fire ants. *Florida Entomologist, 101*(2), 265-272. doi: https://doi.org/10.1653/024.101.0218

Porter, S.D., Plowes, R.M. & Causton, C.E. (2018). The fire ant decapitating fly, *Pseudacteon bifidus* (Diptera: Phoridae): Host specificity and attraction to potential food items. *Florida Entomologist, 101*(1), 55-60. doi: https://doi.org/10.1653/024.101.0111

Sánchez-Restrepo, A.F., Chifflet, L., Confalonieri, V.A., Tsutsui, N.D., Pesquero, M.A. & Calcaterra, L.A. (2020). A species delimitation approach to uncover cryptic species in the South American fire ant decapitating flies (Diptera: Phoridae: *Pseudacteon*). *PLOS ONE, 15*(11). doi: https://doi.org/10.1371/journal.pone.0236086

Scudder, G.G.E. & Cannings, R.A. (2006). The Diptera families of British Columbia. http://www.for.gov.bc.ca/hfd/library/FIA/2006/FSP_Y062001b.pdf

Shikano, I., Woolcott, J., Cloonan, K., Andreadis, S. & Jenkins, N.E. (2021). Biology of mushroom phorid flies, *Megaselia halterata* (Diptera: Phoridae): Effects of temperature, humidity, crowding, and compost stage. *Environmental Entomology, 50*(1), 149–153. doi: https://doi.org/10.1093/ee/nvaa142

Piophilidae

López-García, J., Angell, C. & Martín-Vega, D. (2020). Wing morphometrics for the identification of Nearctic and Palaearctic Piophilidae (Diptera) of forensic relevance. *Forensic Science International, 309*. doi: https://doi.org/10.1016/j.forsciint.2020.110192

Martín-Vega, D. (2011). Skipping clues: Forensic importance of the family Piophilidae (Diptera). *Forensic Science International, 212*(1–3), 1-5. doi: https://doi.org/10.1016/j.forsciint.2011.06.016

McAlpine, J.F. (1987). Piophilidae. In McAlpine, J.F., Peterson, B.V., Shewell, G.E., Teskey, H.J., Vockeroth, J.R. & Wood, D.M. (Eds), Manual of Nearctic Diptera. Volume 2. (pp. 844 – 852). Research Branch Agriculture Canada.

Michalski, M., Gadawski, P., Klemm, J. & Szpila, K. (2021). New species of soldier fly—*Sargus bipunctatus* (Scopoli, 1763) (Diptera: Stratiomyidae), recorded from a human corpse in Europe—A case report. *Insects, 12*(4), 302. doi: https://doi.org/10.3390/insects12040302

Prado e Castro, C., Cunha, E., Serrano, A. & García, M.D. (2012). *Piophila megastigmata* (Diptera: Piophilidae): First records on human corpses. *Forensic Science International, 214*(1–3), 23-26. doi: https://doi.org/10.1016/j.forsciint.2011.07.009

Scudder, G.G.E. & Cannings, R.A. (2006). The Diptera families of British Columbia. http://www.for.gov.bc.ca/hfd/library/FIA/2006/FSP_Y062001b.pdf

Taleb, M., Tail, G. & Açıkgöz, H.N. (2019). DNA barcoding of *Stearibia nigriceps* (Meigen) and *Piophila casei* (Linnaeus) (Diptera: Piophilidae) from Algeria and the first African report of *Stearibia nigriceps*. *International Journal of Legal Medicine, 134*(3), 895-902. doi: https://doi.org/10.1007/s00414-019-02223-w

Pipunculidae

Hardy, D.E. (1987). Pipunculidae. In McAlpine, J.F., Peterson, B.V., Shewell, G.E., Teskey, H.J., Vockeroth, J.R. & Wood, D.M. (Eds), Manual of Nearctic Diptera. Volume 2. (pp. 745 – 748). Research Branch Agriculture Canada.

El-Hawagry, M.S., El-Azab, S.A. & Gilbert, F. (2019). Catalogue of the family Pipunculidae in Egypt (Diptera: Cyclorrhapha). *African Entomology, 27*(1), 238-244. doi: https://doi.org/10.4001/003.027.0238

Jervis, M.A. (1992). A taxonomic revision of the pipunculid fly genus *Chalarus* Walker, with particular reference to the European fauna. *Zoological Journal of the Linnean Society, 105*(3), 243-352. doi: https://doi.org/10.1111/j.1096-3642.1992.tb01232.x

Kehlmaier, C. (2005). Taxonomic studies on Palaearctic and Oriental Eudorylini (Diptera: Pipunculidae), with the description of three new species. *Zootaxa, 1030*(1), 1-48. doi: https://doi.org/10.11646/zootaxa.1030.1.1

May, Y.Y. (1979). The biology of *Cephalops curtifrons* (Diptera: Pipunculidae), an endoparasite of *Stenocranus minutus* (Hemiptera: Delphacidae). *Zoological Journal of the Linnean Society, 66*(1), 15–29. https://doi.org/10.1111/j.1096-3642.1979.tb01899.x

Parker, H.L. (1967). Notes on the biology of *Tomosvaryella frontata* (Diptera: Pipunculidae), a parasite of the leafhopper *Opsius stactogalus* on *Tamarix*. *Annals of the Entomological Society of America, 60*(2), 292–295. doi: https://doi.org/10.1093/aesa/60.2.292

Ramos-Pastrana, Y. & Rafael, J.A. (2021). *Tomosvaryella* Aczél (Diptera: Pipunculidae) of Colombia, with description of two new species. *Zootaxa, 4985*(1), 37-68. doi: https://doi.org/10.11646/zootaxa.4985.1.2

Scudder, G.G.E. & Cannings, R.A. (2006). The Diptera families of British Columbia. http://www.for.gov.bc.ca/hfd/library/FIA/2006/FSP_Y062001b.pdf

Skevington, J. & Marshall, S.A. (1997). First record of a big-headed fly, *Eudorylas alternatus* (cresson) (Diptera: Pipunculidae), reared from the subfamily Cicadellinae (Homoptera: Cicadellidae), with an overview of pipunculid-host associations in the nearctic region. *The Canadian Entomologist, 129*(3), 387 – 398. doi: https://doi.org/10.4039/Ent129387-3

Suh, S-J. & Kwon, Y-J. (2010). Three species of the genus *Pipunculus* Latreille (Insecta: Diptera: Pipunculidae) new to Korea. *Animal Systematics, Evolution and Diversity, 26*(3), 187-190. doi: https://doi.org/10.5635/KJSZ.2010.26.3.187

Platystomatidae

Han, H-Y. (2013). A checklist of the families Lonchaeidae, Pallopteridae, Platystomatidae, and Ulidiidae (Insecta: Diptera: Tephritoidea) in Korea with notes on 12 species new to Korea. *Animal Systematics, Evolution and Diversity, 29*(1), 56-69. doi: http://dx.doi.org/10.5635/ASED.2013.29.1.56

McMichael, B.A., Foote, B.A. & Bowker, B.D. (1990). Biology of *Rivellia melliginis* (Diptera: Platystomatidae), a consumer of the nitrogen-fixing root nodules of black locust (Leguminosae). *Annals of the Entomological Society of America, 83*(5), 967-974. doi: https://doi.org/10.1093/aesa/83.5.967

Polleniidae

Cerretti, P., Stireman III, J.O., Badano, D., Gisondi, S., Rognes, K., Giudice, G.L. & Pape, T. (2019). Reclustering the cluster flies (Diptera: Oestroidea, Polleniidae). *Systematic Entomology, 44*(4), 957-972. doi: https://doi.org/10.1111/syen.12369

De Jong, G.D., Meyer, F. & Goddard, J. (2021). An annotated list of the blow flies and cluster flies (Diptera: Calliphoridae, Polleniidae) of Mississippi. *Transactions of the American Entomological Society, 147*(3), 827-843. doi: https://doi.org/10.3157/061.147.0304

Gisondi, S., Rognes, K., Badano, D., Pape, T. & Cerretti, P. (2020). The world Polleniidae (Diptera, Oestroidea): key to genera and checklist of species. *ZooKeys, 971*, 105-155. doi: https://doi.org/10.3897/zookeys.971.51283

El Husseini, M.M.M. (2019). Endo- or ecto-parasitism with the cluster fly, *Pollenia dasypoda* Portochisky (Diptera: Calliphoridae), based on the diameter of its host body, the earthworm *Allolobophora caligonosa* (Sav.). *Egyptian Journal of Biological Pest Control, 29*(1), 1-5. doi: https://doi.org/10.1186/s41938-019-0150-8

Rognes, K. (1987). The taxonomy of the *Pollenia rudis* species-group in the Holarctic Region (Diptera: Calliphoridae). *Systematic Entomology, 12*(4), 475-502. doi: https://doi.org/10.1111/j.1365-3113.1987.tb00219.x

Szpila, K., Piwczyński, M., Glinkowski, W., Lutz, L., Akbarzadeh, K., Baz, A., Johnston, N.P. & Grzywacz, A. (2023). First molecular phylogeny and species delimitation of West Palaearctic *Pollenia* (Diptera: Polleniidae). *Zoological Journal of the Linnean Society, 197*(1), 267–282. doi: https://doi.org/10.1093/zoolinnean/zlac035

Taleb, M., Tail, G. & Açıkgöz, H.N. (2022). Molecular identification of the potentially forensically relevant cluster flies *Pollenia rudis* (Fabricius) and *Pollenia vagabunda* (Meigen) (Diptera: Polleniidae) - non-recorded species in Algeria. *Forensic Sciences Research, 7*(1), 69-77. doi: https://doi.org/10.1080/20961790.2020.1857937

Thomson, A.J. & Davies, D.M. (1973). The biology of *Pollenia rudis*, the cluster fly (Diptera: Calliphoridae). *The Canadian Entomologist, 105*(7), 985 – 990. doi: https://doi.org/10.4039/Ent105985-7

Vezsenyi, K.A., Langer, S.V., Samkari, B.A. & Beresford, D.V. (2022). The history and current state of cluster flies (Diptera: Polleniidae: *Pollenia*) in North America, with new Canadian provincial records. *Canadian Entomologist, 154*(1). doi: https://doi.org/10.4039/tce.2022.11

Psilidae

Ellis, W.N. (2020). *Parasites.* Plant Parasites of Europe. https://bladmineerders.nl/parasites/

Petherbridge, F.R., Wright, D.W. & Davies, P.G. (1942). Investigations on the biology and control of the carrot fly (*Psila rosae* f.). *Annals of Applied Biology, 29*(4), 380-392. doi: https://doi.org/10.1111/j.1744-7348.1942.tb06142.x

Scudder, G.G.E. & Cannings, R.A. (2006). The Diptera families of British Columbia. http://www.for.gov.bc.ca/hfd/library/FIA/2006/FSP_Y062001b.pdf

Steyskal, G.C. (1987). Psilidae. In McAlpine, J.F., Peterson, B.V., Shewell, G.E., Teskey, H.J., Vockeroth, J.R. & Wood, D.M. (Eds), Manual of Nearctic Diptera. Volume 2. (pp. 781 – 784). Research Branch Agriculture Canada.

Psychodidae

Azmiera, N., Low, V.L. & Heo, C.C. (2021). Colonization of rabbit carcasses by drain fly larvae, *Psychoda* sp. (Diptera: Psychodidae): The first report. *Acta Parasitologica, 66*(2), 706-709. doi: https://doi.org/10.1007/s11686-020-00313-z

Bejarano, E.E. & Estrada, L.G. (2016). Family Psychodidae. *Zootaxa, 4122*(1), 187-238. doi: http://dx.doi.org/10.11646/zootaxa.4122.1.20

Headlee, T.J. & Beckwith, C.S. (1918). Sprinkling sewage filter fly *Psychoda Alternata* Say. *Journal of Economic Entomology, 11*(5), 395–401. doi: https://doi.org/10.1093/jee/11.5.395

Kriska, G. (2013). *Freshwater Invertebrates in Central Europe*. Springer. doi: https://doi.org/10.1007/978-3-7091-1547-3_22

Quate, L.W. & Vockeroth, J.R. (1981). Psychodidae. In McAlpine, J.F., Peterson, B.V., Shewell, G.E., Teskey, H.J., Vockeroth, J.R. & Wood, D.M. (Eds), Manual of Nearctic Diptera. Volume 1. (pp. 293 – 300). Research Branch Agriculture Canada.

Rachesky, S. & Petty, H.B. (1968). Control of *Psychoda alternata* at a Wastewater Sewerage Plant. *Journal of Economic Entomology, 61*(4), 1118–1119. doi: https://doi.org/10.1093/jee/61.4.1118

Salmela, J., Kvifte, G.M. & More, A. (2012). Description of a new *Psychoda* Latreille species from Fennoscandia (Diptera: Psychodidae). *Zootaxa, 3313*(1), 34-43. doi: https://doi.org/10.11646/zootaxa.3313.1.4

Scudder, G.G.E. & Cannings, R.A. (2006). The Diptera families of British Columbia. http://www.for.gov.bc.ca/hfd/library/FIA/2006/FSP_Y062001b.pdf

Ptychopteridae

Alexander, C.P. (1981). Ptychopteridae. In McAlpine, J.F., Peterson, B.V., Shewell, G.E., Teskey, H.J., Vockeroth, J.R. & Wood, D.M. (Eds), Manual of Nearctic Diptera. Volume 1. (pp. 325 – 328). Research Branch Agriculture Canada.

Fasbender, A. (2014). *Phylogeny and diversity of the phantom crane flies (Diptera: Ptychopteridae)*. [Doctoral dissertation, Iowa State University]. Iowa State University ProQuest Dissertations Publishing.

Kriska, G. (2013). *Freshwater Invertebrates in Central Europe*. Springer. doi: https://doi.org/10.1007/978-3-7091-1547-3_22

Mattingly, R.L. (1987). Resource utilization by the freshwater deposit feeder *Ptychoptera townesi* (Diptera: Ptychopteridae). *Freshwater Biology, 18*(2), 241-253. doi: https://doi.org/10.1111/j.1365-2427.1987.tb01311.x

Scudder, G.G.E. & Cannings, R.A. (2006). The Diptera families of British Columbia. http://www.for.gov.bc.ca/hfd/library/FIA/2006/FSP_Y062001b.pdf

Shao, J. & Kang, Z. (2021). New species of the genus *Ptychoptera* Meigen, 1803 (Diptera, Ptychopteridae) from Zhejiang, China with an updated key to Chinese species. *ZooKeys, 1070*(3), 87-99. doi: https://doi.org/10.3897/zookeys.1070.67779

Wiberg-Larsen, P., Hansen, S.B., Rinne, A., Viitanen, E. & Krogh, P.H. (2021). Key to Ptychopteridae (Diptera) larvae of Northern Europe, with notes on distribution and biology. *Zootaxa, 5039*(2), 179-200. doi: https://doi.org/10.11646/zootaxa.5039.2.2

Wolf, B., Zwick, P. & Marxsen, J. (1997). Feeding ecology of the freshwater detritivore *Ptychoptera paludosa* (Diptera, Nematocera). *Freshwater Biology, 38*(2), 375-386. doi: https://doi.org/10.1046/j.1365-2427.1997.00250.x

Rhagionidae

Imada, Y. & Kato, M. (2016). Bryophyte-feeders in a basal brachyceran lineage (Diptera: Rhagionidae: Spaniinae): Adult oviposition behavior and changes in the larval mouthpart morphology accompanied with the diet shifts. *PLOS ONE, 11*(11). doi: https://doi.org/10.1371/journal.pone.0165808

James, M.T. & Turner, W.J. (1981). Rhagionidae. In McAlpine, J.F., Peterson, B.V., Shewell, G.E., Teskey, H.J., Vockeroth, J.R. & Wood, D.M. (Eds), Manual of Nearctic Diptera. Volume 1. (pp. 483 – 488). Research Branch Agriculture Canada.

Lee, J. & Suh, S.J. (2022). A new species of the snipe fly genus *Rhagio* Fabricius (Diptera: Rhagionidae) from Korea. *Journal of Asia-Pacific Biodiversity, 15*(3), 370-374. doi: https://doi.org/10.1016/j.japb.2022.03.005

Roberts, M.J. (1969). Structure of the mouthparts of the larvae of the flies *Rhagio* and *Sargus* in relation to feeding habits. *Journal of Zoology, 159*(3), 381-398. doi: https://doi.org/10.1111/j.1469-7998.1969.tb08453.x

Santos, C.M.D. & Carmo, D.D.D. (2016). Family Rhagionidae. *Zootaxa, 4122*(1), 246-248. doi: http://dx.doi.org/10.11646/zootaxa.4122.1.22

Scudder, G.G.E. & Cannings, R.A. (2006). The Diptera families of British Columbia. http://www.for.gov.bc.ca/hfd/library/FIA/2006/FSP_Y062001b.pdf

Turner, W.J. (1974). A revision of the genus *Symphoromyia* Frauenfeld (Diptera: Rhagionidae): I. Introduction. Subgenera and species-groups. Review of biology. *The Canadian Entomologist, 106*(8), 851 – 868. doi: https://doi.org/10.4039/Ent106851-8

Sarcophagidae

Allen, G.R. & Pape, T. (1996). Description of female and biology of *Blaesoxipha ragg* Pape (Diptera: Sarcophagidae), a parasitoid of *Sciarasaga quadrata* Rentz (Orthoptera: Tettigoniidae) in Western Australia. *Australian Journal of Entomology, 35*(2), 147-151. doi: https://doi.org/10.1111/j.1440-6055.1996.tb01379.x

Brousseau, P-M., Giroux, M. & Handa, I.T. (2020). First record on the biology of *Sarcophaga* (Bulbostyla) (Diptera, Sarcophagidae). *ZooKeys, 909*, 59-66. doi: https://doi.org/10.3897/zookeys.909.46488

Buenaventura, E. (2021). Museomics and phylogenomics with protein-encoding ultraconserved elements illuminate the evolution of life history and phallic morphology of flesh flies (Diptera: Sarcophagidae). *BMC Ecology and Evolution, 21*(1), 70. doi: https://doi.org/10.1186/s12862-021-01797-7

Buenaventura, E., Szpila, K., Cassel, B.k., Wiegmann, B.M. & Pape, T. (2019). Anchored hybrid enrichment challenges the traditional classification of flesh flies (Diptera: Sarcophagidae). *Systematic Entomology, 45*(2), 281-301. doi: https://doi.org/10.1111/syen.12395

Castro, C.V., Buenaventura, E., Sánchez-Rodríguez, J.D. & Wolff, M. (2017). Flesh flies (Diptera: Sarcophagidae: Sarcophaginae) from the Colombian Guajira biogeographic province, an approach to their ecology and distribution. *Zoologia, 34,* 1-11. doi: https://doi.org/10.3897/zoologia.34.e12277

Coupland, J.B. & Barker, G.M. (2004). Diptera as predators and parasitoids of terrestrial gastropods, with emphasis on Phoridae, Calliphoridae, Sarcophagidae, Muscidae and Fanniidae. *Natural Enemies of Terrestrial Molluscs,* 85-158. doi: https://doi.org/10.1079/9780851993195.0085

Danyk, T., Mackauer, M. & Johnson, D.L. (2005). The influence of host suitability on the range of grasshopper species utilized by *Blaesoxipha atlanis* (Diptera: Sarcophagidae) in the field. *Bulletin of Entomological Research, 95*(6), 571-578. doi: https://doi.org/10.1079/BER2005388

de Mello-Patiu, C.A. (2016). Family Sarcophagidae. *Zootaxa, 4122*(1), 884-903. doi: http://dx.doi.org/10.11646/zootaxa.4122.1.75

Falk, S. & Mulley, J.F. (2023). The genome sequence of the lesser worm flesh fly, *Sarcophaga (Sarcophaga) subvicina* (Baranov, 1937). *Wellcome Open Research, 8*, 65. doi: https://doi.org/10.12688/wellcomeopenres.18717.1

Pickens, L.G. (1981). The life history and predatory efficiency of *Ravinia lherminieri* (Diptera: Sarcophagidae) on the face fly (Diptera: Muscidae). *The Canadian Entomologist, 113*(6), 523 – 526. doi: https://doi.org/10.4039/Ent113523-6

Rosenmejer, T. & Pape, T. (2018). Five new species of the flesh fly genus *Boettcheria* (Diptera: Sarcophagidae). *Zootaxa, 4483*(3), 579-590. doi: https://doi.org/10.11646/zootaxa.4483.3.9

Scudder, G.G.E. & Cannings, R.A. (2006). The Diptera families of British Columbia. http://www.for.gov.bc.ca/hfd/library/FIA/2006/FSP_Y062001b.pdf

Shewell, G.E. (1987). Sarcophagidae. In McAlpine, J.F., Peterson, B.V., Shewell, G.E., Teskey, H.J., Vockeroth, J.R. & Wood, D.M. (Eds), Manual of Nearctic Diptera. Volume 2. (pp. 1159 – 1186). Research Branch Agriculture Canada.

Szpila, K., Mądra, A., Jarmusz, M. & Matuszewski, S. (2015). Flesh flies (Diptera: Sarcophagidae) colonising large carcasses in Central Europe. *Parasitology Research, 114*(6), 2341-2348. doi: https://doi.org/10.1007/s00436-015-4431-1

Vairo, K.P., Queiroz, M.M.C., Mendonça, P.M., Barbosa, R.R. & de Carvalho, C.J.B. (2015). Description of immature stages of the flesh fly *Peckia (Sarcodexia) lambens* (Wiedemann) (Diptera:

Sarcophagidae) provides better resolution for taxonomy and forensics. *Tropical Zoology, 28*(3), 114-125. doi: https://doi.org/10.1080/03946975.2015.1057435

Wong, E.S., Dahlem, G.A., Stamper, T.I. & DeBry, R.W. (2014). Discordance between morphological species identification and mtDNA phylogeny in the flesh fly genus *Ravinia* (Diptera : Sarcophagidae). *Invertebrate Systematics, 29*(1), 1-11. doi: https://doi.org/10.1071/IS14018

Yan, L., Buenaventura, E., Pape, T., Kutty, S.N., Bayless, K.M. & Zhang, D. (2020). A phylotranscriptomic framework for flesh fly evolution (Diptera, Calyptratae, Sarcophagidae). *Cladistics, 37*(5), 540-558. doi: https://doi.org/10.1111/cla.12449

Scathophagidae

Blanckenhorn, W.U., Pemberton, A.J., Bussière, L.F., Roembke, J. & Floate, K.D. (2010). A review of the natural history and laboratory culture methods for the yellow dung fly, *Scathophaga stercoraria*. *Journal of Insect Science, 10*(11), 1-17. doi: https://doi.org/10.1673/031.010.1101

Chagnon, M-E. & Sinclair, B.J. (2020). Revision of the Nearctic species of *Gimnomera* Rondani (Diptera: Scathophagidae), with morphological phylogeny and DNA barcodes. *Zootaxa, 4853*(3), 369-403. doi: https://doi.org/10.11646/zootaxa.4853.3.3

Ellis, W.N. (2020). *Parasites*. Plant Parasites of Europe. https://bladmineerders.nl/parasites/

Kutty, S.N., Bernasconi, M.V., Šifner, F. & Meier, R. (2007). Sensitivity analysis, molecular systematics and natural history evolution of Scathophagidae (Diptera: Cyclorrhapha: Calyptratae). *Cladistics, 23*(1), 64-83. doi: https://doi.org/10.1111/j.1096-0031.2006.00131.x

Neff, S.E. (1968). Observations on the immature stages of *Gimnomera cerea* and *G. incisurata* (Diptera: Aanthomyiidae, Scatophaginae). *The Canadian Entomologist, 100*(1), 74 – 83. doi: https://doi.org/10.4039/Ent10074-1

Neff, S.E. & Wallace, J.B. (1969). Observations on the Immature Stages of *Cordilura (Achaetella) deceptiva* and *C. (a.) varipes*. *Annals of the Entomological Society of America, 62*(4), 775–785. doi: https://doi.org/10.1093/aesa/62.4.775

Ovchinnikov, A.N. & Ovtshinnikova, O.G. (2013). The muscular system of the ovipositor of *Spaziphora hydromyzina* (Fallen) (Diptera, Scathophagidae). *Entomological Review*, *93*(8), 987-990. doi: https://doi.org/10.1134/S001387381308006X

Ozerov, A.L. (2017). A review of the genus *Microprosopa* Becker, 1894 (Diptera: Scathophagidae) of Russia. *Russian Entomology, 25*(1), 71-99. doi: https://doi.org/10.15298/rusentj.26.1.10

Ozerov, A.L. (2017). A new species of the genus *Scathophaga* Meigen, 1803 (Diptera, Scathophagidae) from Yakutia (Russia). *Entomological Review, 97*(1), 132-135. doi: https://doi.org/10.1134/S0013873817010134

Scudder, G.G.E. & Cannings, R.A. (2006). The Diptera families of British Columbia. http://www.for.gov.bc.ca/hfd/library/FIA/2006/FSP_Y062001b.pdf

Vockeroth, J.R. (1987). Scathophagidae. In McAlpine, J.F., Peterson, B.V., Shewell, G.E., Teskey, H.J., Vockeroth, J.R. & Wood, D.M. (Eds), Manual of Nearctic Diptera. Volume 2. (pp. 1085 – 1098). Research Branch Agriculture Canada.

Wallace, J.B. & Neff, S.E. (1971). Biology and immature stages of the genus *Cordilura* (Diptera: Scatophagidae) in the Eastern United States. *Annals of the Entomological Society of America 64*(6), 1310–1311. doi: https://doi.org/10.1093/aesa/64.6.1310

### Sciaridae

Babytskiy, A.I., Zuieva, O.A. & Bezsmertna, O.O. (2018). *Peyerimhoffia vagabunda* – new sciarid species (Sciaridae, Diptera) for the entomofauna of Ukraine. *Biosystems diversity, 26*(3), 245-249. doi: https://doi.org/10.15421/011837

Babytskiy, A.I., Moroz, M.S., Kalashnyk, S.O., Bezsmertna, O.O., Dudiak, I.D. & Voitsekhivska, O.V. (2019). New findings of pest sciarid species (Diptera, Sciaridae) in Ukraine, with the first record of *Bradysia* difformis. *Biosystems diversity, 27*(2), 131-141. doi: https://doi.org/10.15421/011918

Chang, S-C., Shentu, H., Shih, H-T. & Lin, S-F. (2004). First record of the pest *Bradysia impatiens* (Diptera: Sciaridae) and overview of *Bradysia* species in Taiwan. *Oriental Insects, 58*(2), 157-171. doi: https://doi.org/10.1080/00305316.2023.2252806

Farsani, N.S., Zamani, A.A., Abbasi, S. & Kheradmand, K. (2013). Effect of temperature and button mushroom varieties on life history of *Lycoriella auripila* (Diptera: Sciaridae). *Journal of Economic Entomology, 106*(1), 115–123. doi: https://doi.org/10.1603/EC12241

Farsani, N.S., Zamani, A.A., Abbasi, S. & Kheradmand, K. (2019). Effect of temperature and mushroom varieties on biology of fungus gnat, *Lycoriella auripila* (Diptera: Sciaridae). *Biologia, 75*(5), 723-731. doi: https://doi.org/10.2478/s11756-019-00340-w

Gou, Y., Quandahor, P., Zhang, Y., Coulter, J.A., Liu, C. & Gao, Y. (2020). Host plant nutrient contents influence nutrient contents in *Bradysia cellarum* and *Bradysia impatiens*. *PLoS One, 15*(4), p.e0226471. doi: https://doi.org/10.1371/journal.pone.0226471

Heiri, O. & Lotter, A.F. (2007). Sciaridae in lake sediments: indicators of catchment and stream contribution to fossil insect assemblages. *Journal of Paleolimnology, 38*(2), 183-189. doi: https://doi.org/10.1007/s10933-006-9068-8

Heller, K., Hippa, H. & Vilkamaa, P. (2015). Taxonomy of *Bradysia* Winnertz (Diptera, Sciaridae) in the Northern Holarctic, with the description of four new species. *European Journal of Taxonomy, 122*. doi: http://dx.doi.org/10.5852/ejt.2015.122

Hussey, N.W. & Gurney, B. (1968). Biology and control of the sciarid *Lycoriella auripila* Winn. (Diptera: Lycoriidae) in mushroom culture. *Annals of Applied Biology, 62*(3), 395-403. doi: https://doi.org/10.1111/j.1744-7348.1968.tb05451.x

Menzel, F., Gammelmo, Ø., Olsen, K.M. & Köhle, A. (2020). The black fungus gnats (Diptera, Sciaridae) of Norway – Part I: species records published until December 2019, with an updated checklist. *ZooKeys, 957*(1), 17- 104. doi: https://doi.org/10.3897/zookeys.957.46528

Frank Menzel, F., Jukka Salmela, J. & Vilkamaa, P. (2020). New species and new records of black fungus gnats (Diptera: Sciaridae) from the Viidumäe Nature Reserve, Estonia. *European Journal of Taxonomy, 720*(1), 62-76. doi: https://doi.org/10.5852/ejt.2020.720.1115

Mohrig, W., Kauschke, E. & Broadley, A. (2017). Black fungus gnats (Diptera: Sciaridae) of Queensland, Australia. Part I. Genera *Chaetosciara* Frey, *Corynoptera* Winnertz, *Cratyna* Winnertz, *Epidapu*s Haliday, *Keilbachia* Mohrig, *Lobosciara* Steffan, *Phytosciara* Frey and *Scatopsciara* Edwards. *Zootaxa, 4303*(4), 451-481. doi: https://doi.org/10.11646/ZOOTAXA.4303.4.1

Salmela, J. & Vilkamaa, P. (2005). Sciaridae (Diptera) from central Finland: Faunistics and taxonomy. *Entomologica Fennica, 16*(4), 287–300. doi: https://doi.org/10.33338/ef.84273

Sawangproh, W. & Cronberg, N. (2016). Life history traits of the liverwort herbivore *Scatopsciara cunicularius* (Diptera: Sciaridae). *Annals of the Entomological Society of America, 109*(3), 343-349. doi: https://doi.org/10.1093/aesa/saw004

Sawangproh, W., Ekroos, J. & Cronberg, N. (2016). The effect of ambient temperature on larvae of *Scatopsciara cunicularius* (Diptera: Sciaridae) feeding on the thallose liverwort Marchantia polymorpha. *European Journal of Entomology, 113*(1), 259-264. doi: https://doi.org/10.14411/eje.2016.030

Scudder, G.G.E. & Cannings, R.A. (2006). The Diptera families of British Columbia. http://www.for.gov.bc.ca/hfd/library/FIA/2006/FSP_Y062001b.pdf

Shamshad, A. (2010). The development of integrated pest management for the control of mushroom sciarid flies, *Lycoriella ingenua* (Dufour) and *Bradysia ocellaris* (Comstock), in cultivated mushrooms. *Pest Management Science, 66*(10), 1063-1074. doi: https://doi.org/10.1002/ps.1987

Shin, S., Jung, S., Menzel, F., Heller, K., Lee, H. & Lee, S. (2013). Molecular phylogeny of black fungus gnats (Diptera: Sciaroidea: Sciaridae) and the evolution of larval habitats. *Molecular Phylogenetics and Evolution, 66*(3), 833-846. doi: https://doi.org/10.1016/j.ympev.2012.11.008

Shin, S., Menzel, F., Heller, K., Lee, H. & Lee, S. (2014). Review of the genus *Cratyna* Winnertz (Diptera: Sciaridae) in Korea, including the description of a new species. *Zootaxa, 3794*(3), 344-354. doi: http://dx.doi.org/10.11646/zootaxa.3794.3.2

Shin, S., Lee, H., Menzel, F. & Lee, S. (2020). Taxonomic study on the *Phytosciara* genus group (Diptera: Sciaridae) in Korea, including the description of a new species. *Journal of Asia-Pacific Entomology, 23*(2), 358-363. doi: https://doi.org/10.1016/j.aspen.2020.01.005

Steffan, W.A. (1981). Sciaridae. In McAlpine, J.F., Peterson, B.V., Shewell, G.E., Teskey, H.J., Vockeroth, J.R. & Wood, D.M. (Eds), Manual of Nearctic Diptera. Volume 1. (pp. 247 – 256). Research Branch Agriculture Canada.

Sueyoshi, M. & Yoshimatsu, S-I. (2019). Pest species of a fungus gnat genus *Bradysia* Winnertz (Diptera: Sciaridae) injuring agricultural and forestry products in Japan, with a review on taxonomy of allied species. *Entomological Science, 22*(3), 317-333. doi: https://doi.org/10.1111/ens.12373

Sueyoshi, M., Nakamura, S. & Menzel, F. (2022). A new species of *Hyperlasion* Schmitz (Diptera: Sciaridae), causing periodic outbreaks in Japan. *Zootaxa, 5168*(4), 451-463. doi: https://doi.org/10.11646/zootaxa.5168.4.5

Vilkamaa, P. & Komonen, A. (2001). Redescription and biology of *Trichosia (Baeosciara) sinuate* Menzel & Mohrig (Diptera: Sciaridae). *Entomologica Fennica, 12*(1), 46–49. doi: https://doi.org/10.33338/ef.84095

Xi, L., Zhou, X., Liu, Y., Li, Y., Shi, H., Wu, H. & Huang, J. (2021). Behavioural response of the fungus gnat, *Bradysia impatiens* (Diptera: Sciaridae) towards certain edible mushrooms and saprophytic fungi. *Journal of Applied Entomology, 145*(5), 458-466. doi: https://doi.org/10.1111/jen.12857

Zhu, G., Luo, Y., Xue, M., Zhao, H., Sun, X. & Wang, X. (2017). Effects of feeding on different host plants and diets on *Bradysia odoriphaga* population parameters and tolerance to heat and insecticides. *Journal of Economic Entomology, 110*(6), 2371-2380. doi: https://doi.org/10.1093/jee/tox242

Sciomyzidae

Ahmed, K.S.D., Stephens, C., Bistline-East, A., Williams, C.D., Mc Donnell, R.J., Carnaghi, M., Huallacháin, D.Ó. & Gormally, M.J. (2019). Biological control of pestiferous slugs using *Tetanocera elata* (Fabricius) (Diptera: Sciomyzidae): Larval behavior and feeding on slugs exposed to *Phasmarhabditis hermaphrodita* (Schneider, 1859). *Biological Control, 135*, 1-8. doi: https://doi.org/10.1016/j.biocontrol.2019.04.003

Bistline-East, A., Carey, J.G.J., Colton, A., Day, M.F. & Gormally, M.J. (2018). Catching flies with honey(dew): adult marsh flies (Diptera: Sciomyzidae) utilize sugary secretions for high-carbohydrate diets. *Environmental Entomology, 47*(6), 1632-1641. doi: https://doi.org/10.1093/ee/nvy155

Bistline-East, A., Williams, C.D. & Gormally, M.J. (2020). Nutritional ecology of predaceous *Tetanocera elata* larvae and the physiological effects of alternative prey utilization. *Biocontrol, 65*(3), 285-296. doi: https://doi.org/10.1007/s10526-020-09997-8

Chapman, E.G., Foote, B.A., Malukiewicz, J. & Hoeh, W.R. (2006). Parallel evolution of larval morphology and habitat in the snail-killing fly genus *Tetanocera*. *Journal of Evolutionary Biology, 19*(5), 1459-1474. doi: https://doi.org/10.1111/j.1420-9101.2006.01132.x

Chapman, E.G., Przhiboro, A.A., Harwood, J.D., Foote, B.A. & Hoeh, W.R. (2012). Widespread and persistent invasions of terrestrial habitats coincident with larval feeding behavior transitions during snail-killing fly evolution (Diptera: Sciomyzidae). *BMC Evolutionary Biology, 12*(1), 175. doi: https://doi.org/10.1186/1471-2148-12-175

Foote, B.A. (2008). Biology and immature stages of snail-killing flies belonging to the genus *Tetanocera* (Diptera: Sciomyzidae). IV. Life histories of predators of land snails and slugs. *Annals of Carnegie Museum, 77*(2), 301-312. doi: https://doi.org/10.2992/0097-4463-77.2.301

Hynes, T.M., Giordani, I., Larkin, M., Mc Donnell, R.J. & Gormally, M.J. (2014). Larval feeding behaviour of *Tetanocera elata* (Diptera: Sciomyzidae): potential biocontrol agent of pestiferous slugs. Biocontrol Science and Technology, 24(9), 1077-1082. doi: https://doi.org/10.1080/09583157.2014.912259

Kazerani, F., Mortelmans, J., Farashiani, M.E. & Thorn, S. (2020). A new species of *Pherbellia* (Diptera: Sciomyzidae) from Iran. *Zootaxa, 4772*(2), 361-370. doi: https://doi.org/10.11646/zootaxa.4772.2.7

Knutson, L.V. (1987). Sciomyzidae. In McAlpine, J.F., Peterson, B.V., Shewell, G.E., Teskey, H.J., Vockeroth, J.R. & Wood, D.M. (Eds), Manual of Nearctic Diptera. Volume 2. (pp. 927 – 940). Research Branch Agriculture Canada.

Kriska, G. (2013). *Freshwater Invertebrates in Central Europe*. Springer. doi: https://doi.org/10.1007/978-3-7091-1547-3_22

Manguin, S. & Vala, J-C. (1989). Prey consumption by larvae of *Tetanocera ferruginea* (Diptera: Sciomyzidae) in relation to number of snail prey species available. *Annals of the Entomological Society of America, 82*(5), 588–592. doi: https://doi.org/10.1093/aesa/82.5.588

Murphy, W.L., Knutson, L.V., Chapman, E.G. Mc Donnell, R.J., Williams, C.D., Foote, B.A. & Vala, J. (2012). Key aspects of the biology of snail-killing Sciomyzidae flies. *Annual Review of Entomology, 57*(1), 425-447. doi: https://doi.org/10.1146/annurev-ento-120710-100702

Murphy, W.L., Mathis, W.N. & Knutson, L.V. (2018). Comprehensive taxonomic, faunistic, biological, and geographic inventory and analysis of the Sciomyzidae (Diptera: Acalyptratae) of the Delmarva region and nearby states in eastern North America. *Zootaxa, 4430*(1), 1-299. doi: https://doi.org/10.11646/zootaxa.4430.1.1

Murphy, W.L., Abercrombie, J., González, C.R. & Knutson, L. (2023). Overview of the Sciomyzidae (Diptera: Sciomyzoidea) of the Americas south of the United States. *Zootaxa, 5345*(1), 1-113. doi: https://doi.org/10.11646/zootaxa.5345.1.1

Nerudová-Horsáková, J., Murphy, W.L. & Vala, J-C. (2016). Biology and immature stages of *Pherbellia limbata* (Diptera: Sciomyzidae), a parasitoid of the terrestrial snail *Granaria frumentum. Zootaxa, 4117*(1), 48-62. doi: http://doi.org/10.11646/zootaxa.4117.1.2

Scudder, G.G.E. & Cannings, R.A. (2006). The Diptera families of British Columbia. http://www.for.gov.bc.ca/hfd/library/FIA/2006/FSP_Y062001b.pdf

Son, Y. & Suh, S.J. (2020). Taxonomic review of the genus *Pherbellia* Robineau-Desvoidy (Diptera: Sciomyzidae) from Korea. *Journal of Asia-Pacific Biodiversity, 13*(4), 577-582. doi: https://doi.org/10.1016/j.japb.2020.09.003

Trelka, D.G. & Foote, B.A. (1970). Biology of slug-killing *Tetanocera* (Diptera: Sciomyzidae). *Annals of the Entomological Society of America, 63*(3), 877–895. doi: https://doi.org/10.1093/aesa/63.3.877

Vala, J-C. & Williams, C.D. (2015). Sciomyzidae Fallén, 1820 (Diptera) collected in the Mercantour National Park, France. *Zoosystema, 37*(4), 611-619. doi: https://doi.org/10.5252/z2015n4a7

Sepsidae

Ang, Y., Meier, R., Su, K. F-Y. & Rajaratnam, G. (2017). Hidden in the urban parks of New York City: *Themira lohmanus*, a new species of Sepsidae described based on morphology, DNA sequences, mating behavior, and reproductive isolation (Sepsidae, Diptera). *ZooKeys, 698*, 95-111. doi: https://doi.org/10.3897/zookeys.698.13411

Pont, A.C. & Meier, R. (2002). *The Sepsidae (Diptera) of Europe*. Brill.

Randall, M., Coulson, J.C. & Butterfield, J. (1981). The distribution and biology of Sepsidae (Diptera) in upland regions of northern England. *Ecological Entomology, 6*(2), 183-190. doi: https://doi.org/10.1111/j.1365-2311.1981.tb00604.x

Scudder, G.G.E. & Cannings, R.A. (2006). The Diptera families of British Columbia. http://www.for.gov.bc.ca/hfd/library/FIA/2006/FSP_Y062001b.pdf

Steyskal, G.C. (1987). Sepsidae. In McAlpine, J.F., Peterson, B.V., Shewell, G.E., Teskey, H.J., Vockeroth, J.R. & Wood, D.M. (Eds), Manual of Nearctic Diptera. Volume 2. (pp. 945 – 950). Research Branch Agriculture Canada.

Simuliidae

Adler, P.H. & Reeves, W.K. (2023). North–South differentiation of black flies in the Western Cordillera of North America: A new species of *Prosimulium* (Diptera: Simuliidae). *Diversity, 5*(2), 212. doi: https://doi.org/10.3390/d15020212

Adler, P.H., Cheke, R.A., Post, R.J. (2010). Evolution, epidemiology, and population genetics of black flies (Diptera: Simuliidae). *Infection, genetics and evolution, 10*(7), 846-865. doi: https://doi.org/10.1016/j.meegid.2010.07.003

Al-Shaer, L., Pierce, A.K., Larson, D. & Hancock, R. (2015). Notes on facultative predation in *Prosimulium* larvae (Diptera: Simuliidae) in alpine and subalpine streams in Colorado. *Journal of the American Mosquito Control Association, 31*(1), 113-116. doi: https://doi.org/10.2987/14-6460.1

Brabrand, Å., Bremnes, T., Koestler, A. G., Marthinsen, G., Pavels, H., Rindal, E., Raastad, J. E., Saltveit, S. J. & Johnsen, A. (2013). Mass occurrence of bloodsucking blackflies in a regulated river reach: localization of oviposition habitat of *Simulium truncatum* using DNA barcoding: mass occurrence bloodsucking blackflies. *River Research and Applications, 30*(5), 602-608. doi: https://doi.org/10.1002/rra.2669

Brannin, M.T., O'Donnell, M.K. & Fingerut, J. (2012). Effects of larval size and hydrodynamics on the growth rates of the black fly *Simulium tribulatum*. *Integrative Zoology, 9*(1), 61-69. doi: https://doi.org/10.1111/1749-4877.12016

Cameron, A.E. (2013). *The morphology and biology of a Canadian cattle-infesting black fly: Simulium simile Mall. (Diptera, Simuliidae).* Agriculture and Agri-Food Canada.

Coscarón, S., Esquivel, M., Moulton, J.K., Coscarónarias, C.L. & Bernal, S.I. (2004). *Simulium* (Hearlea) Vargas, Martinez Palacios, & Diaz Najera (Diptera: Simuliidae): Taxonomic revision and cladistic analysis. *Zootaxa, 396*, 1-52.

Craig, D.A., Currie, D.C., Hunter, F.F. & Spironello, M. (2006). A taxonomic revision of the southwestern Pacific subgenus *Hebridosimulium* (Diptera: Simuliidae: Simulium). *Zootaxa, 1380*(1), 1-90. doi: https://doi.org/10.11646/zootaxa.1380.1.1

Crosskey, R.W. (2007). A new species of *Prosimulium* Roubaud (Diptera: Simuliidae) from Rhodesia. *Journal of Natural History, 2*(4), 487-495. doi: https://doi.org/10.1080/00222936800770981

Currie, D.C. & Hunter, F.F. (2003). A new species of *Stegopterna* Enderlein, and its relationship to the allotriploid species *St. mutata* (Malloch, 1914) (Diptera: Simuliidae). *Zootaxa, 214*(1), 1-11. doi: https://doi.org/10.11646/zootaxa.214.1.1

Diaz, S.A., Moncada, L.I., Murcia, C.H., Lotta, I.A., Matta, N.E. & Adler, P.H. (2015). Integrated taxonomy of a new species of black fly in the subgenus *Trichodagmia* (Diptera: Simuliidae) from the Páramo Region of Colombia. *Zootaxa, 3914*(5), 541-557. doi: https://doi.org/10.11646/zootaxa.3914.5.3

Finn, D.S. & Alder, P.H. (2006). Population genetic structure of a rare high-elevation black fly, *Metacnephia coloradensis*, occupying Colorado lake outlet streams. *Freshwater Biology, 51*(12), 2240-2251. doi: https://doi.org/10.1111/j.1365-2427.2006.01647.x

Gaudreau, C., Larue, B. & Charpentier, G. (2010). Molecular comparison of Quebec and Newfoundland populations of the blackfly, *Simulium vittatum*, species complex. *Medical and Veterinary Entomology, 24*(2), 214-217. doi: https://doi.org/10.1111/j.1365-2915.2009.00844.

Hamada, N., Hernandez, L.M. & Luz, S.L.B. (2006). Taxonomy of *Simulium guaporense* Py-Daniel (Diptera: Simuliidae) from Brazil, with the first description of males and females. *Zootaxa, 1104*(1), 23-34. doi: https://doi.org/10.11646/zootaxa.1104.1.2

Hernandez, L.M., Dias, A.P.A. de L., Maia-Herzog, M. & Shelley, A.J. (2006). Taxonomy of *Simulium (Inaequalium) petropoliense* Coscarón (Diptera: Simuliidae) from Brazil, with the first description of the male and larva. *Zootaxa, 1275*(1), 1-20. doi: https://doi.org/10.11646/zootaxa.1275.1.1

Hunter, F.F., Sutcliffe, J.F. & Downe, A.E.R. (1993). Blood-feeding host preferences of the isomorphic species *Simulium venustum* and *S. truncatum*. *Medical and Veterinary Entomology, 7*(2), 105-110. doi: https://doi.org/10.1111/j.1365-2915.1993.tb00661.x

Kriska, G. (2013). *Freshwater Invertebrates in Central Europe*. Springer. doi: https://doi.org/10.1007/978-3-7091-1547-3_22

Martin, P.J.S. & Edman, J.D. (1993). Assimilation rates of different particulate foods for *Simulium verecundum* (Diptera: Simuliidae). *Journal of Medical Entomology, 30*(4), 805-809. doi: https://doi.org/10.1093/jmedent/30.4.805

McCreadie, J.W. & Colbo, M.H. (1991). Spatial distribution patterns of larval cytotypes of the *Simulium venustum/verecundum* complex (Diptera: Simuliidae) on the Avalon Peninsula, Newfoundland: factors associated with occurrence. *Canadian Journal of Zoology, 69*(10), 2651-2659. doi: https://doi.org/10.1139/z91-373

McCreadie, J.W. & Colbo, M.H. (1993). Larval and pupal microhabitat selection by *Simulium truncatum* Lundström, *S. rostratum* Lundström, and *S. verecundum* AA (Diptera: Simuliidae). *Canadian Journal of Zoology, 71*(2), 358-367. doi: https://doi.org/10.1139/z93-050

McCreadie, J.W., Colbo, M.H. & Bennett, G.F. (1986). The influence of weather on host seeking and blood feeding of *Prosimulium mixtum* and *Simulium venustum/verecundum* Complex (Diptera: Simuliidae). *Journal of Medical Entomology, 23*(3), 289–297. doi: https://doi.org/10.1093/jmedent/23.3.289

Moulton, J.K. & Adler, P.H. (2002). Taxonomy and biology of *Simulium clarkei* Stone & Snoddy (Diptera: Simuliidae), a poorly known black fly of the southeastern United States. *Zootaxa, 31*(1), 1-7. doi: https://doi.org/10.11646/zootaxa.31.1.1

Peterson, B.V. (1981). Simuliidae. In McAlpine, J.F., Peterson, B.V., Shewell, G.E., Teskey, H.J., Vockeroth, J.R. & Wood, D.M. (Eds), Manual of Nearctic Diptera. Volume 1. (pp. 355 – 392). Research Branch Agriculture Canada.

Scudder, G.G.E. & Cannings, R.A. (2006). The Diptera families of British Columbia. http://www.for.gov.bc.ca/hfd/library/FIA/2006/FSP_Y062001b.pdf

Takaoka, H. & Tenedero, V.F. (2019). Two new species of the *Simulium (Simulium) tuberosum* species-group (Diptera: Simuliidae) from Palawan, the Philippines. *Zootaxa, 4568*(2), 383-393. doi: https://doi.org/10.11646/zootaxa.4568.2.12

Zhang, Y. (2006). Balancing food availability and hydrodynamic constraint: phenotypic plasticity and growth in *Simulium noelleri* blackfly larvae. *Oecologia, 147*(1), 39-46. doi: https://doi.org/10.1007/s00442-005-0243-9

Zinchenko, M.O., Sukhomlin, K.B., Zinchenko, O.P. & Tepliuk, V.S. (2021). The biology of *Simulium noelleri* and *Simulium dolini*:  morphological, ecological and molecular data. *Biosystems diversity, 29*(2), 180-184. doi: https://doi.org/10.15421/012122

Sphaeroceridae

Ellis, W.N. (2020). *Parasites*. Plant Parasites of Europe. https://bladmineerders.nl/parasites/

Fredeen, F.J.H. & Glen, G.S. (1970). The survival and development of *Leptocera caenosa* (Diptera: Sphaeroceridae) in laboratory cultures. *The Canadian Entomologist, 102*(2), 164 – 171. doi: https://doi.org/10.4039/Ent102164-2

Jindřich, R. (2013). The fauna of Acalyptrate families Trixoscelididae, Chyromyidae and Sphaeroceridae (Diptera) in the Gemer area (Central Slovakia): supplement 2. *Casopis Slezského Zemského Muzea, 62*(2), 155-172. doi: https://doi.org/10.2478/cszma-2013-0017

Jindřich, R. (2019). First Sphaeroceridae (Diptera) endemic to Madeira – three new terricolous species of *Spelobia* and *Pullimosina*. *Acta Entomologica Musei Nationalis Pragae, 59*(1), 107-124. doi: https://doi.org/10.2478/aemnp-2019-0009

Roháček, J. & Przhiboro, A.A. (2022). *Pullimosina (Pullimosina) turfosa* sp. nov. and other Sphaeroceridae (Diptera) from peat bogs in the North Caucasus (Russia). *ZooKeys, 1132*, 1-49. doi: https://doi.org/10.3897/zookeys.1132.94579

Marshall, S.A. & Richards, O.W. (1987). Sphaeroceridae. In McAlpine, J.F., Peterson, B.V., Shewell, G.E., Teskey, H.J., Vockeroth, J.R. & Wood, D.M. (Eds), Manual of Nearctic Diptera. Volume 2. (pp. 933 – 1006). Research Branch Agriculture Canada.

Pitkin, B.R. (1986). Bait, habitat preferences and the phenology of some lesser dung flies (Diptera: Sphaeroceridae) in Britain. *Journal of Natural History, 20*(6), 1283-1295. doi: https://doi.org/10.1080/00222938600770851

Scudder, G.G.E. & Cannings, R.A. (2006). The Diptera families of British Columbia. http://www.for.gov.bc.ca/hfd/library/FIA/2006/FSP_Y062001b.pdf

Su, L. & Liu, G. (2009). A review of the genus *Terrilimosina* Roháček (Diptera: Sphaeroceridae, Limosininae) from China. *The Pan-Pacific Entomologist, 85*(2), 51-57. doi: https://doi.org/10.3956/2009-10.1

Wheeler, T.A. (1994). Systematics of the New World *Rachispoda* Lioy (Diptera: Sphaeroceridae): morphology, key to species groups, and revisions of the *atra, fuscipennis, limosa* and *vespertina* species groups. Journal of Natural History, 29(1), 159-230. doi: https://doi.org/10.1080/00222939500770091

Stratiomyidae

Brammer, C.A. & Von Dohlen, C.D. (2010). Morphological phylogeny of the variable fly family Stratiomyidae (Insecta, Diptera). *Zoologica Scripta, 39*(4), 363-377. https://doi.org/10.1111/j.1463-6409.2010.00430.x

James, M.T. (1981). Stratiomyidae. In McAlpine, J.F., Peterson, B.V., Shewell, G.E., Teskey, H.J., Vockeroth, J.R. & Wood, D.M. (Eds), Manual of Nearctic Diptera. Volume 1. (pp. 497 – 512). Research Branch Agriculture Canada.

Kriska, G. (2013). *Freshwater Invertebrates in Central Europe*. Springer. doi: https://doi.org/10.1007/978-3-7091-1547-3_22

Lee, J. & Suh, S.J. (2022). First record of the soldier fly genus *Beris* Latreille (Diptera, Stratiomyidae) from Korea, with designation of two new synonyms. *Biodiversity Data Journal, 10*. doi: https://doi.org/10.3897/BDJ.10.e80487

Michalski, M., Gadawski, P., Klemm, J. & Szpila, K. (2021). New species of soldier fly—*Sargus bipunctatus* (Scopoli, 1763) (Diptera: Stratiomyidae), recorded from a human corpse in Europe—A case report. *Insects, 12*(4), 302. doi: https://doi.org/10.3390/insects12040302

Scudder, G.G.E. & Cannings, R.A. (2006). The Diptera families of British Columbia. http://www.for.gov.bc.ca/hfd/library/FIA/2006/FSP_Y062001b.pdf

Torres-Toro, J., Pujol-Luz, J.R. & Wolff, M. (2022). Two new species of *Ptecticus* Loew, 1855 (Diptera: Stratiomyidae), from bat guano in a Colombian cave. *Zootaxa, 5116*(1), 61-88. doi: https://doi.org/10.11646/zootaxa.5116.1.3

Syrphidae

Ball, S. & Morris, R. (2015). *Britain's Hoverflies: A Field Guide - Revised and Updated Second Edition*. Princeton University Press.

Barahona-Segovia, R.M., Riera, P., Pañinao-Monsálvez, L., Guzmán, V.V. & Henríquez-Piskulich, P. (2021). Updating the knowledge of the flower flies (Diptera: Syrphidae) from Chile: Illustrated catalog, extinction risk and biological notes. *Zootaxa, 4969*(1), 1-178. doi: https://doi.org/10.11646/zootaxa.4959.1.1

Berthiaume, R., Hébert, C., Pelletier, G. & Cloutier, C. (2016). Seasonal natural history of aphidophagous Syrphidae (Diptera) attacking the balsam twig aphid in balsam fir (Pinaceae) Christmas tree plantations.
*The Canadian Entomologist, 148*(4), 466 – 475. doi: https://doi.org/10.4039/tce.2015.84

Branquart, E. & Hemptinne, J. (2008). Selectivity in the exploitation of floral resources by hoverflies (Diptera: Syrphinae). *Ecography, 23*(6), 732-742. doi: https://doi.org/10.1111/j.1600-0587.2000.tb00316.x

Campbell, D.R., Bischoff, M., Lord, J.M. & Robertson, A.W. (2010). Flower color influences insect visitation in alpine New Zealand. *Ecology, 91*(9), 2638-2649. doi: https://doi.org/10.1890/09-0941.1

Correa, J. (2019). *Syrphidae: A Guide to Natural History and Identification of Common Genera in Santa Cruz County*. [Senior Project, University of California].

Dziock, F. (2005). Evolution of prey specialization in aphidophagous syrphids of the genera *Melanostoma* and *Platycheirus* (Diptera: Syrphidae) 1. Body size, development and prey traits. European Journal of Entomology, 102(3), 413-421. doi: https://doi.org/10.14411/eje.2005.059

El-Hawagry, M.S. & Gilbert, F. (2019). Catalogue of the Syrphidae of Egypt (Diptera). *Zootaxa, 4577*(2), 201-248. doi: https://doi.org/10.11646/zootaxa.4577.2.1

Ellis, W.N. (2020). *Parasites.* Plant Parasites of Europe. https://bladmineerders.nl/parasites/

Gojkovic, N., Francuski, L., Ludoski, J. & Milankov, V. (2020). DNA barcode assessment and population structure of aphidophagous hoverfly *Sphaerophoria scripta*: Implications for conservation biological control. *Ecology and Evolution, 10*(7), 9428-9443. doi: https://doi.org/10.1002/ece3.6631

Goulson, D. & Wright, N.P. (1998). Flower constancy in the hoverflies *Episyrphus balteatus* (Degeer) and *Syrphus ribesii* (L.) (Syrphidae). *Behavioral Ecology, 9*(3), 213–219. doi: https://doi.org/10.1093/beheco/9.3.213

Gresham, S.D.M., Charles, J.G., Sandanayaka, M.W.R. & Bergh, J.C. (2013). Laboratory and field studies supporting the development of *Heringia calcarata* as a candidate biological control agent for *Eriosoma lanigerum* in New Zealand. *BioControl, 58*(5), 645–656. doi: https://doi.org/10.1007/s10526-013-9530-2

Haslett, J.R. (1989). Adult feeding by holometabolous insects: pollen and nectar as complementary nutrient sources for *Rhingia campestris* (Diptera: Syrphidae). *Oecologia, 81*(3), 361–363. doi: https://doi.org/0.1007/BF00377084

Hawkes, W. & Wotton, K. (2022). The genome sequence of the dumpy grass hoverfly, *Melanostoma mellinum* (Linnaeus, 1758). *Wellcome open research, 7*, 59. doi: https://doi.org/10.12688/wellcomeopenres.17615.1

Jamali, R.A., Memom, N., Shah, M.A., Khan, K. & Ansari, A. (2018). Prevalence of aphidophagous hoverflies (Syrphidae: Syrphinae) in relation to their prey, green aphids on Brassica in Dadu. *Journal of Animal and Plant Sciences, 28*(5), 1447.

Jeong, S-H. & Han, H-Y. (2019). A taxonomic revision of the genus *Xylota* Meigen (Diptera: Syrphidae) in Korea. *Zootaxa, 4661*(3), 457-493. doi: https://doi.org/10.11646/zootaxa.4661.3.3

Jiang, S., Li, H., He, L. & Wu, K. (2022). Population fitness of *Eupeodes corollae* Fabricius (Diptera: Syrphidae) feeding on different species of aphids. *Insects, 13*(6), 494. doi: https://doi.org/10.3390/insects13060494

Koval, A.G., Guseva, O.G. & Shpanev, A.M. (2018). Hoverflies (Diptera, Syrphidae) in Agrolandscapes of St. Petersburg and Leningrad Province. *Entomological Review, 98*(6), 702-708. doi: https://doi.org/10.1134/S0013873818060064

Kriska, G. (2013). *Freshwater Invertebrates in Central Europe*. Springer. doi: https://doi.org/10.1007/978-3-7091-1547-3_22

Krivosheina, N.P. (2020). Ecological relations of the hoverfly larvae (Diptera, Syrphidae, Eristalinae) bark inhabitants with xylobiont insects. *Biology Bulletin of the Russian Academy of Sciences, 47*(6), 605-616. doi: https://doi.org/10.1134/S1062359020060096

Lillo, I., Perez-Bañón, C. & Rojo, S. (2021). Life cycle, population parameters, and predation rate of the hover fly *Eupeodes corollae* fed on the aphid *Myzus persicae*. *Entomologia Experimentalis et Applicata, 169*(11), 1027-1038. doi: https://doi.org/10.1111/eea.13090

López-García, G.P., Roig-Juñet, S.A., Pérez-Bañón, C., Mazzitelli, E., Montoya, A.L., Rojo, S. & Mengual, X. (2022). Description of the third-stage larva and puparium of *Platycheirus (Carposcalis) chalconota* (Philippi) (Diptera: Syrphidae) with new information about the trophic interactions and larval habitats. *Neotropical Entomology, 51*, 81–98. doi: https://doi.org/10.1007/s13744-021-00908-9

Mengual, X., Stahls, G. & Rojo, S. (2008). First phylogeny of predatory flower flies (Diptera, Syrphidae, Syrphinae) using mitochondrial COI and nuclear 28S rRNA genes: conflict and congruence with the current tribal classification. *Cladistics, 24*(4), 543-562. doi: https://doi.org/10.1111/j.1096-0031.2008.00200.x

Mengual, X., Stahls, G. & Rojo, S. (2015). Phylogenetic relationships and taxonomic ranking of Pipizine flower flies (Diptera: Syrphidae) with implications for the evolution of aphidophagy. *Cladistics, 31*(5), 491-508. doi: https://doi.org/10.1111/cla.12105

Nedeljković, Z., Ačanski, J., Vujić, A., Obreht, D., Đan, M., Ståhls, G. & Radenković, S. (2013). Taxonomy of *Chrysotoxum festivum* Linnaeus, 1758 (Diptera: Syrphidae) - an integrative approach. *Zoological Journal of the Linnean Society, 169*(1), 84-102. doi: https://doi.org/10.1111/zoj.12052

Nedeljković, Z., Ačanski, J., Đan, M., Obreht-Vidaković, D., Ricarte, A., Vujić, A. & Biesmeijer, J.C. (2015). An integrated approach to delimiting species borders in the genus *Chrysotoxum* Meigen, 1803 (Diptera: Syrphidae), with description of two new species. *Contributions to Zoology, 84*(4), 285-304. doi: https://doi.org/10.1163/18759866-08404002

Nedeljković, Z., Ricarte, A., Zorić, L.Š., Djan, M., Hayat, R., Vujić, A. & Mª Ángeles Marcos-García, Mª Á. (2020). Integrative taxonomy confirms two new West-Palaearctic species allied with *Chrysotoxum vernale* Loew, 1841 (Diptera: Syrphidae). *Organisms Diversity & Evolution, 20*(4), 821-833. doi: https://doi.org/10.1007/s13127-020-00465-w

Pérez-Bañón, C., Hurtado, P., García-Gras, E. & Rojo, S. (2013). SEM studies on immature stages of the drone flies (Diptera, Syrphidae): *Eristalis similis* (Fallen, 1817) and *Eristalis tenax* (Linnaeus, 1758). *Microscopy Research and Technique, 76*(8), 853-861. doi: https://doi.org/10.1002/jemt.22239

Jones, R. (1958). *Heringia senilis* Sack (Diptera: Syrphidae): a hoverfly new to Britain. *British Journal of Entomology and Natural History, 14*(4), 185-194.

Rank, N.E. & Smiley, J.T. (1994). Host-plant effects on *Parasyrphus melanderi* (Diptera: Syrphidae) feeding on a willow leaf beetle *Chrysomela aeneicollis* (Coleoptera: Chrysomelidae). *Ecological Entomology, 19*(1), 31-38. doi: https://doi.org/10.1111/j.1365-2311.1994.tb00387.x

Rizza, A., Campobasso, G., Dunn, P.H. & Stazi, M. (1988). *Cheilosia corydon* (Diptera: Syrphidae), a candidate for the biological control of musk thistle in North America. *Annals of the Entomological Society of America, 81*(2), 225-232. doi: https://doi.org/10.1093/aesa/81.2.225

Rotheray, G.E. (1988). Larval morphology and feeding patterns of four *Cheilosia* species (Diptera: Syrphidae) associated with *Cirsium palustre* L. Scopoli (Compositae) in Scotland. *Journal of natural history, 22*(1), 17-25. doi: https://doi.org/0.1080/00222938800770031

Rotheray, G. & Lyszkowski, R. (2014). Diverse mechanisms of feeding and movement in Cyclorrhaphan larvae (Diptera). Journal of Natural History, 49(35-36), 2139-2211. doi: https://doi.org/10.1080/00222933.2015.1010314

Scudder, G.G.E. & Cannings, R.A. (2006). The Diptera families of British Columbia. http://www.for.gov.bc.ca/hfd/library/FIA/2006/FSP_Y062001b.pdf

Skevington, J.H., Young, A.D., Locke, M.M. & Moran, K.M. (2019). New Syrphidae (Diptera) of North-eastern North America. *Biodiversity Data Journal, 7*. doi: https://doi.org/10.3897/BDJ.7.e36673

Sturza, V.S., Dequech, S.T.B., Toebe, M., Silveira, T.R., Filho, C.A. & Bolzan, A. (2014). *Toxomerus duplicatus* Wiedemann, 1830 (Diptera: Syrphidae) preying on *Microtheca* spp. (Coleoptera: Chrysomelidae) larvae. *Brazilian journal of biology, 74*(3), 656. doi: https://doi.org/10.1590/bjb.2014.0071

van Steenis, J., Young, A.D., Ssymank, A.M., Wu, T-H., Shiao, S-F. & Skevington, J.H. (2019). The species of the genus *Platycheirus* Lepeletier & Serville, 1828 (Diptera, Syrphidae) from Taiwan, with a discussion on intersex specimens. *Journal of Asia-Pacific Entomology, 22*(1), 281-295. doi: https://doi.org/10.1016/j.aspen.2018.12.004

Tijana, N., D.R., Dubravka, M., V.M., Sonja, T., Snežana, J., Smiljka, S. & A.V. (2013). Models of the potential distribution and habitat preferences of the genus *Pipiza* (Syrphidae: Diptera) on the Balkan peninsula. *Archives of Biological Sciences, 65*(3), 1037-1052. doi: https://doi.org/10.2298/ABS1303037N

Vanhaelen, N., Francis, F. & Haubruge, E. (2004). Purification and characterization of glutathione S-transferases from two syrphid flies (*Syrphus ribesii* and *Myathropa florae*). *Comparative Biochemistry and Physiology Part B: Biochemistry and Molecular Biology, 137*(1), 95-100. doi: https://doi.org/10.1016/j.cbpc.2003.10.006

Vockeroth, J.R. & Thompson, F.C. (1987). Syrphidae. In McAlpine, J.F., Peterson, B.V., Shewell, G.E., Teskey, H.J., Vockeroth, J.R. & Wood, D.M. (Eds), Manual of Nearctic Diptera. Volume 2. (pp. 713 – 744). Research Branch Agriculture Canada.

Tabanidae

Hine, J.S. (1903). *Tabanidae of Ohio with a catalogue and bibliography of the species from America north of Mexico.* Press of Spahr & Glenn.

Jones, C.M. (1953). Biology of Tabanidae in Florida. *Journal of Economic Entomology, 46*(6), 1108–1109. doi: https://doi.org/10.1093/jee/46.6.1108

Kelly-Hope, L., Paulo, R., Thomas, B., Brito, M., Unnasch, T.R. & Molyneux, D. (2017). Loa loa vectors *Chrysops* spp.: perspectives on research, distribution, bionomics, and implications for elimination of lymphatic filariasis and onchocerciasis. *Parasites & Vectors, 10*(1), 172. doi: https://doi.org/10.1186/s13071-017-2103-y

Kriska, G. (2013). *Freshwater Invertebrates in Central Europe*. Springer. doi: https://doi.org/10.1007/978-3-7091-1547-3_22

Leprince, D.J. & Bigras-Poulin, M. (1990). Gonotrophic status, follicular development, sperm presence, and sugar-feeding patterns in a *Hybomitra lasiophthalma* population (Diptera: Tabanidae). *Journal of Medical Entomology, 27*(1), 31–35. doi: https://doi.org/10.1093/jmedent/27.1.31

Leprince, D.J. & Lewis, D.J. (1983). Aspects of the biology of female *Chrysops univittatus* (Diptera: Tabanidae) in southwestern Quebec. *The Canadian Entomologist, 115*(4), 421 – 425. doi: https://doi.org/10.4039/Ent115421-4

Lewis, L.F. (1959). On the Biology of *Chrysops flavida* in the Yazoo- Mississippi Delta. *Journal of Economic Entomology, 52*(5), 884–888. doi: https://doi.org/10.1093/jee/52.5.884

Ossowski, A. & Hunter, F.F. (2012). Distribution patterns, body size, and sugar-feeding habits of two species of *Chrysops* (Diptera: Tabanidae). *The Canadian Entomologist, 132*(2), 213-221. doi: https://doi.org/10.4039/Ent132213-2

Pechuman, L.L. & Teskey, H.J. (1981). Tabanidae. In McAlpine, J.F., Peterson, B.V., Shewell, G.E., Teskey, H.J., Vockeroth, J.R. & Wood, D.M. (Eds), Manual of Nearctic Diptera. Volume 1. (pp. 463 – 478). Research Branch Agriculture Canada.

Scudder, G.G.E. & Cannings, R.A. (2006). The Diptera families of British Columbia. http://www.for.gov.bc.ca/hfd/library/FIA/2006/FSP_Y062001b.pdf

Sofield, R.K. & Hansens, E.J. (1982). Notes on Biology *Hybomitra daeckei* (Hine) (Diptera: Tabanidae). *Entomological News, 93*, 67-69.

Teskey, H.J., Shemanchuk, J.A. & Weintraub, J. (1987). *Hybomitra agora*, a new species of Tabanidae (Diptera) from Western North America. *Canadian Entomologist, 119*(12), 1117-1122. doi: https://doi.org/10.4039/Ent1191117-12

Tachinidae

Al-Dobai, S., Reitz, S. & Sivinski, J. (2012). Tachinidae (Diptera) associated with flowering plants: estimating floral attractiveness. Biological Control, 61(3), 230-239. doi: https://doi.org/10.1016/j.biocontrol.2012.02.008

Allen, H.W. (1925). *Biology of the red-tailed Tachina-fly, Winthemia quadripustulata Fabr*. Mississippi Agricultural Experiment Station.

Bora, D. & Deka, B. (2014). Role of visual cues in host searching behaviour of *Exorista sorbillans* Widemann, a parasitoid of Muga Silk Worm, *Antheraea assama* Westwood. *Journal of Insect Behavior, 27*(1), 92-104. doi: https://doi.org/10.1007/s10905-013-9409-1

Dindo, M.L., Rezaei, M. & De Clercq, P. (2019). Improvements in the rearing of the Tachinid parasitoid *Exorista larvarum* (Diptera: Tachinidae): Influence of adult food on female longevity and reproduction capacity. *Journal of Insect Science, 19*(2). doi: https://doi.org/10.1093/jisesa/iey122

Foerster, L.A. & Doetzer, A.K. (2002). Host instar preference of *Peleteria robusta* (Wiedman) (Diptera: Tachinidae) and development in relation to temperature. *Neotropical Entomology, 31*(3), 405-409. doi: https://doi.org/10.1590/S1519-566X2002000300009

Lutovinovas, E., Malenovský, I., Tóthová, A., Ziegler, J. & Vaňhara, J. (2013). Taxonomic approach to the tachinid flies *Dinera carinifrons* (Fallén) (Diptera: Tachinidae) and *Dinera fuscata* Zhang and Shima using molecular and morphometric data. *Journal of Insect Science, 13*(1), 139. doi: https://doi.org/10.1673/031.013.13901

Martel, V., Thireau, J. & Régnière, J. (2021). Manual inoculation of host larvae with first instar maggots as a rearing technique for the larval parasitoid *Actia interrupta* (Diptera: Tachinidae). *Biocontrol Science and Technology, 32*(1), 110-113. doi: https://doi.org/10.1080/09583157.2021.1967292

Morewood, D.W. & Wood, M.D. (2002). Host utilization by *Exorista thula* Wood (sp. nov.) and *Chetogena gelida* (Coquillett) (Diptera: Tachinidae), parasitoids of arctic *Gynaephora* species (Lepidoptera: Lymantriidae). *Polar Biology, 25*(8), 575-582. doi: https://doi.org/10.1007/s00300-002-0382-y

Prebble, M.L. (1935). *Actia diffidens* Curran, a parasite of *Peroneavariana* (Fernald) in Cape Breton, Nova Scotia. *Canadian Journal of Research, 12*(2), 216-227. doi: https://doi.org/10.1139/cjr35-016

Régnière, J., Thireau, J., Saint-Amant, R. & Martel, V. (2021). Modeling climatic influences on three parasitoids of low-density spruce budworm populations. Part 3: *Actia interrupta* (Diptera: Tachinidae). *Forests, 12*(11), 1471. doi: https://doi.org/10.3390/f12111471

Scaramozzino, P.L., Di Giovanni, F., Loni, A., Gisondi, S., Lucchi, A. & Cerretti, P. (2020). Tachinid (Diptera, Tachinidae) parasitoids of *Lobesia botrana* (Denis & Schiffermüller, 1775) (Lepidoptera, Tortricidae) and other moths. *ZooKeys, 934*, 111-140. doi: https://doi.org/10.3897/zookeys.934.50823

Scudder, G.G.E. & Cannings, R.A. (2006). The Diptera families of British Columbia. http://www.for.gov.bc.ca/hfd/library/FIA/2006/FSP_Y062001b.pdf

Tachi, T. (2010). Three new species of *Exorista* Meigen (Diptera: Tachinidae), with a discussion of the evolutionary pattern of host use in the genus. *Journal of Natural History, 45*(19-20), 1165-1197. doi: https://doi.org/10.1080/00222933.2011.552803

Wood, D.M. (1987). Tachinidae. In McAlpine, J.F., Peterson, B.V., Shewell, G.E., Teskey, H.J., Vockeroth, J.R. & Wood, D.M. (Eds), Manual of Nearctic Diptera. Volume 2. (pp. 1193 – 1270). Research Branch Agriculture Canada.

Tipulidae

Alexander, C.P. & Byers, G.W. (1981). Tipulidae. In McAlpine, J.F., Peterson, B.V., Shewell, G.E., Teskey, H.J., Vockeroth, J.R. & Wood, D.M. (Eds), Manual of Nearctic Diptera. Volume 1. (pp. 153 – 190). Research Branch Agriculture Canada.

Canhoto, C. & and Graça, M.A.S. (1995). Food value of introduced eucalypt leaves for a Mediterranean stream detritivore: *Tipula lateralis*. *Freshwater Biology, 34*(2), 209-214. doi: https://doi.org/10.1111/j.1365-2427.1995.tb00881.x

Canhoto, C. & and Graça, M.A.S. (2006). Digestive tract and leaf processing capacity of the stream invertebrate *Tipula lateralis*. *Canadian Journal of Zoology, 84*(8), 1087-1095. doi: https://doi.org/10.1139/Z06-092

Jo, J. (2017). Description of larval and pupal stages of *Tipula (Nippotipula) sinica* (Diptera, Tipulidae) from South Korea with ecological notes. *Animal Systematics, Evolution and Diversity, 33*(1), 56-59. doi: https://doi.org/10.5635/ASED.2017.33.1.036

Levente-Péter Kolcsár, L-P., Oosterbroek, P., Olsen, K.M., Paramonov, N.M., Gavryushin, D.I., Pilipenko, V.E., Polevoi, A.V., Eiroa, E., Andersson, M., Dufour, C., Syratt, M., Kurina, O., Lindström, M., Starý, J., Lantsov, V.I., Wiedeńska, J., Pape, T., Friman, M., Peeters, K., Gritsch, W., JSalmela, J., Viitanen, E., Aristophanous, M., Janević, D. and Watanabe, K. (2023). Contribution to the Knowledge of Cylindrotomidae, Pediciidae and Tipulidae (Diptera: Tipuloidea): First records of 86 species from various European countries. *Diversity, 15*(3), 336. doi: https://doi.org/10.3390/d15030336

Kostina, N.V., Chernysheva, A.N., Vecherskii, M.V. & Kuznetsova, T.A. (2020). Microbial nitrogen fixation in the intestine of Tipulidae *Tipula maxima* Larvae. *Biology Bulletin, 47,* 35–39. doi: https://doi.org/10.1134/S1062359020010069

Kriska, G. (2013). *Freshwater Invertebrates in Central Europe*. Springer. doi: https://doi.org/10.1007/978-3-7091-1547-3_22

Petersen, M.J. (2013). Evidence of a climatic niche shift following North American introductions of two crane flies (Diptera; genus *Tipula*). *Biological Invasions, 15*(4), 885-897. doi: https://doi.org/10.1007/s10530-012-0337-3

Podeniene, V., Gelhaus, J.K. & Yadamsuren, O. (2006). The last instar larvae and pupae of *Tipula (Arctotipula)* (Diptera, Tipulidae) from Mongolia. *Proceedings of the Academy of Natural Sciences of Philadelphia, 155*(1), 79-105. doi: https://doi.org/10.1635/i0097-3157-155-1-79.1

Pritchard, G. (1976). Growth and development of larvae and adults of *Tipula sacra* Alexander (Insecta: Diptera) in a series of abandoned beaver ponds. *Canadian Journal of Zoology, 54*(2), 266-284. doi: https://doi.org/10.1139/z76-030

Pritchard, G. (1983). Biology of Tipulidae. *Annual Review of Entomology, 28*(1), 1-22. doi: https://doi.org/10.1146/annurev.en.28.010183.000245

Rao, S., Liston, A., Crampton, L. & Takeyasu, J. (2006). Identification of larvae of exotic *Tipula paludosa* (Diptera: Tipulidae) and *T. oleracea* in North America using mitochondrial cytB sequences. *Annals of the Entomological Society of America, 99*(1), 33-40. doi: https://doi.org/10.1603/0013-8746(2006)099[0033:IOLOET]2.0.CO2

Salmela, J. & Petrašiūnas, A. (2014). Checklist of the infraorder Tipulomorpha (Trichoceridae, Tipuloidea) (Diptera) of Finland. *ZooKeys, 441*(441), 21-36. doi: https://doi.org/10.3897/zookeys.441.7533

Scudder, G.G.E. & Cannings, R.A. (2006). The Diptera families of British Columbia. http://www.for.gov.bc.ca/hfd/library/FIA/2006/FSP_Y062001b.pdf

Sivell, O. & Sivell, D. (2023). The genome sequence of a Tiger Cranefly, *Nephrotoma flavescens* (Linnaeus, 1758). *Wellcome open research, 8*, 148. doi: https://doi.org/10.12688/wellcomeopenres.19203.1

Smith, R.M., Young, M.R. & Marquiss, M. (2008). Bryophyte use by an insect herbivore: does the crane-fly *Tipula montana* select food to maximise growth? *Ecological Entomology, 26*(1), 83-90. doi: https://doi.org/10.1046/j.1365-2311.2001.00297.x

White, J.H. (1951). Observations on the life history and biology of *Tipula Lateralis* Meig. *Annals of Applied Biology, 38*(4), 847-858. doi: https://doi.org/10.1111/j.1744-7348.1951.tb07855.x

Trichoceridae

Dong, F., Shih, C. & Ren, D. (2014). Two new species of Trichoceridae from the Middle Jurassic Jiulongshan Formation of Inner Mongolia, China. *ZooKeys, 411*, 145-160. doi: https://doi.org/10.3897/zookeys.411.6858

Laurence, B.R. (1955). On the life history of *Trichocera saltator* (Harris) (Diptera, Trichoceridae). *Proceedings of the Zoological Society of London, 126*(2), 235-244. doi: https://doi.org/10.1111/j.1096-3642.1956.tb00434.x

Potocka, M. & Krzemińska, E. (2018). *Trichocera maculipennis* (Diptera)—an invasive species in Maritime Antarctica. *PeerJ*. doi: https://doi.org/10.7717/peerj.5408

Scudder, G.G.E. & Cannings, R.A. (2006). The Diptera families of British Columbia. http://www.for.gov.bc.ca/hfd/library/FIA/2006/FSP_Y062001b.pdf

Volonterio, O., de León, R.P., Convey, P. & Krzemińska, E. (2013). First record of Trichoceridae (Diptera) in the maritime Antarctic. *Polar Biology, 36*(8), 1125-1131. doi: https://doi.org/10.1007/s00300-013-1334-4

Ulidiidae

Brunel, O. & Rull, J. (2010). The natural history and unusual mating behavior of *Euxesta bilimeki* (Diptera: Ulidiidae). *Annals of the Entomological Society of America, 103*(1), 111–119. doi: https://doi.org/10.1603/008.103.0114

El-Hawagry, M.S. (2021). The family Ulidiidae in Egypt (Diptera: Tephritoidea). *African Entomology, 29*(2), 445-462. doi: https://doi.org/10.4001/003.029.0445

Han, H-Y. (2013). A Checklist of the families Lonchaeidae, Pallopteridae, Platystomatidae, and Ulidiidae (Insecta: Diptera: Tephritoidea) in Korea with notes on 12 species new to Korea. *Animal Systematics, Evolution and Diversity, 29*(1), 56-69. doi: http://dx.doi.org/10.5635/ASED.2013.29.1.56

Kameneva, E.P. & Korneyev, V.A. (2012). A new species of *Herina* Robineau-Desvoidy, 1830 (Diptera: Ulidiidae) from Turkey, with the key to species of *oscillans* group. *Zootaxa, 3548*(1), 69-74. doi: https://doi.org/10.11646/zootaxa.3548.1.5

Xylophagidae

James, M.T. (1981). Xylophagidae. In McAlpine, J.F., Peterson, B.V., Shewell, G.E., Teskey, H.J., Vockeroth, J.R. & Wood, D.M. (Eds), Manual of Nearctic Diptera. Volume 1. (pp. 489 – 492). Research Branch Agriculture Canada.

Scudder, G.G.E. & Cannings, R.A. (2006). The Diptera families of British Columbia. http://www.for.gov.bc.ca/hfd/library/FIA/2006/FSP_Y062001b.pdf
